# Targeted Kinase Degradation via the KLHDC2 Ubiquitin E3 Ligase

**DOI:** 10.1101/2022.12.17.520883

**Authors:** Younghoon Kim, Christina Seo, Eunhye Jeon, Inchul You, Kyubin Hwang, Namkyoung Kim, Ha-Soon Choi, Stephen M. Hinshaw, Nathanael S. Gray, Taebo Sim

## Abstract

Chemically induced protein degradation is a powerful strategy for perturbing cellular biochemistry. The predominant mechanism of action for protein degrader drugs involves induced proximity between the cellular ubiquitin conjugation machinery and the target. Unlike traditional small molecule enzyme inhibition, targeted protein degradation can clear an undesired protein from cells. We demonstrate here the use of peptide ligands for Kelch-Like Homology Domain Containing protein 2 (KLHDC2), a substrate adaptor protein and member of the cullin-2 (CUL2) ubiquitin ligase complex, for targeted protein degradation. Peptide-based bivalent compounds that can induce proximity between KLHDC2 and target proteins cause degradation of the targeted factors. The cellular activity of these compounds depends on KLHDC2 binding. This work demonstrates the utility of KLHDC2 for targeted protein degradation and exemplifies a strategy for the rational design of new peptide-based ligands useful for this purpose.

## Introduction

Targeted protein degradation is an alternative to conventional small molecule-based enzyme inhibition. Compounds that induce protein degradation can cause near-complete clearance of targets despite sub-stoichiometric occupancy by acting catalytically (*1, 2*). These molecules comprise three components: *i*) a ligand for a ubiquitin E3 ligase, *ii*) a ligand for a degradation target (the so-called warhead), and *iii*) a linker to connect the first two components. Degrader drugs are commonly called PROTACs, for PROteolysis Targeting Chimeras (*3*). PROTAC-induced proximity between a ubiquitin E3 ligase and a target (often called a neo-substrate) catalyzes the transfer of ubiquitin to the target, thus marking this species for degradation.

Of the estimated ∼600 human ubiquitin E3 ligases, only two are commonly used for targeted protein degradation: VHL and CRBN. Chemical ligands for these E3 ligases are synthetically tractable, have well-described modes of target engagement, and have favorable pharmacological properties (*4-7*). Several other ubiquitin E3 ligase proteins have been used, but the corresponding chemical ligands have poorly understood modes of E3 ligase engagement, produce only modest degradation of target substrates, or induce degradation of the repurposed E3 ligase itself (*8*).

The paucity of available small molecules that bind ubiquitin E3 ligases limits the development of therapeutically useful compounds for targeted protein degradation. PROTACs with matching substrate binding components and different E3 ligase ligands (VHL versus CRBN) often show dramatic differences in potency for the targeted substrate proteins, a quality reflected in their proteome-wide target profiles (*9, 10*).

Perhaps more troubling, it is anticipated that resistance to PROTAC drugs will arise from downregulation or mutation of the ubiquitin ligase proteins required for their activities (*11*). Such resistance is less likely to occur if multiple ligases can be exploited simultaneously or in series. Thus, expanding the arsenal of ubiquitin E3 ligases available for targeted protein degradation is of major interest.

The C-end degron pathway is a protein homeostasis pathway dedicated to the clearance of substrates with defined carboxy-terminal motifs (*12-15*). KLHDC2 is one of several substrate recognition components in this pathway. Recognition depends on a tight interaction (<20 nM dissociation constant in biochemically reconstituted systems) between the KLHDC2 beta propeller domain and a 5-10 amino acid substrate peptide ending with Gly-Gly (Figure 1A) (*14*). Recently, high-throughput chemically induced protein proximity experiments have identified KLHDC2 as a tractable ubiquitin E3 ligase for targeted protein degradation (*16*). SelK is a prototypical KLHDC2 substrate that terminates in Gly-Gly when the cellular selenomethoinine pool is depleted. The minimal C-terminal degrons that specify proteins as C-end pathway substrates make the responsible ubiquitin E3 ligases attractive targets for use in targeted protein degradation. We focus on KLHDC2 and describe here new PROTAC molecules that target its activity to kinases.

**Figure 1.**
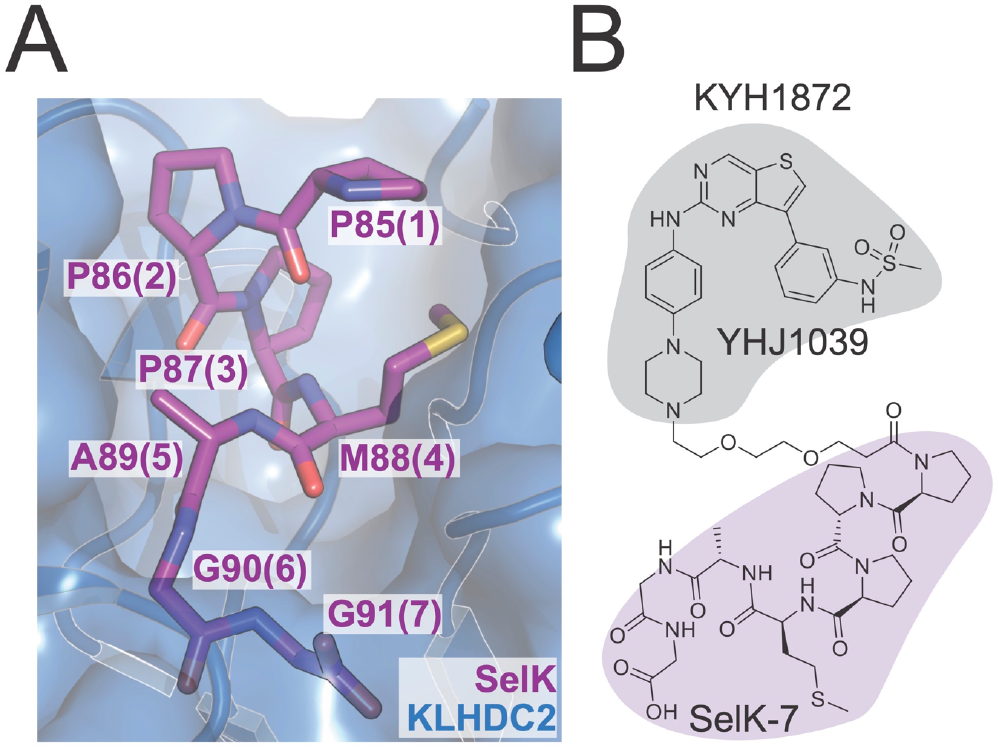
A) Crystal structure showing the KLHDC2-SelK interface (PDB 6D03). SelK is shown in magenta, and individual amino acid residues are labeled at their alpha carbons according to their positions in SelK. Positions in KYH1872 are shown as parentheticals. B) Structure of the KYH1872 compound. The kinase and KLHDC2 ligands are shaded gray and magenta, respectively.

## Results

We have synthesized PROTAC molecules by fusing KLHDC2 substrate peptides derived from SelK with the broad-spectrum kinase inhibitor YHJ1039 (*17*) (Figure 1B). The synthesized molecules consisted of *i*) a C-terminal fragment of the SelK protein, *ii*) a polyethylene glycol (PEG) linker, and *iii*) a promiscuous kinase inhibitor, YHJ1039 (*18*). This inhibitor was previously found to induce kinase degradation when conjugated to gluteramide-containing CRBN recruiters (*18*). In our PROTAC molecules, the most potent of which is KYH1872 (see below), the linkers connect to the N-terminus of the SelK fragment such that the Gly-Gly degron remains available for KLHDC2 binding. Though short linear peptides generally have unfavorable cell permeability properties, there is precedent for using peptide degrons in this way (*19*). The anticipated catalytic mechanism of action and tight KLHDC2-SelK interaction suggests that even trace amounts of cytoplasmic drug availability might drive observable target degradation.

To test whether these compounds induce kinase degradation, we treated MOLM-14 cells with them and detected protein levels by Western blotting. To select degradation targets for the Western blot experiments, we referred to reported inhibition and proteomics profiles of YHJ1039 (*17*) and the related CRBN-based PROTAC DB0614 (*18*), respectively. KYH1872, in which a two-PEG linker connects YHJ1039 with a seven amino acid SelK peptide, induced the degradation of NEK9, FAK, CDK4, CDK6, and WEE1 in cells treated at 1 and 10 μM for 24 hours (Figure 2A). The previously reported CRBN-based PROTAC, DB0614 (*18*), was a more effective kinase degrader than KYH1872 at 1 μM. KYH1872 displayed slightly improved degradation activity when compared with KYH1886, which differs only in the length of its SelK fragment (five versus seven amino acids; Figure S1). Biochemical binding experiments confirmed that the lead degrader compound KYH1872 binds to KLHDC2 protein. Shorter SelK fragments are reported to possess substantially weaker biochemical affinity for KLHDC2, (*14*), which is consistent with the Western blot results described above. However, we could not discriminate the KLHDC2 binding affinities of KYH1872 and KYH1886 in the binding assay (Figure S2A-B; Table 1), as both have dissociation constants that are tighter than the limit of detection.

**Table 1.** Properties of reported compounds. Biochemical binding and cell proliferation values are reported as log_10_[compound] (M), and the assays are described in the Supporting Information. Cell viability measurements were taken 72 hours after treatment of MOLM-14 cells.

| Compound | KLHDC2 binding ($K_D$) | Cell viability ($EC_{50}$) |
| --- | --- | --- |
| YHJ1039 |  | -8.30 |
| DB0625 |  | -8.14 |
| KYH1872 | < -7.7 | -6.19 |
| KYH2011 | -6.78 | -4.34 |
| KYH1996 | < -7.7 | -5.48 |
| KYH1886 | < -7.7 | -5.86 |

**Figure 2.**
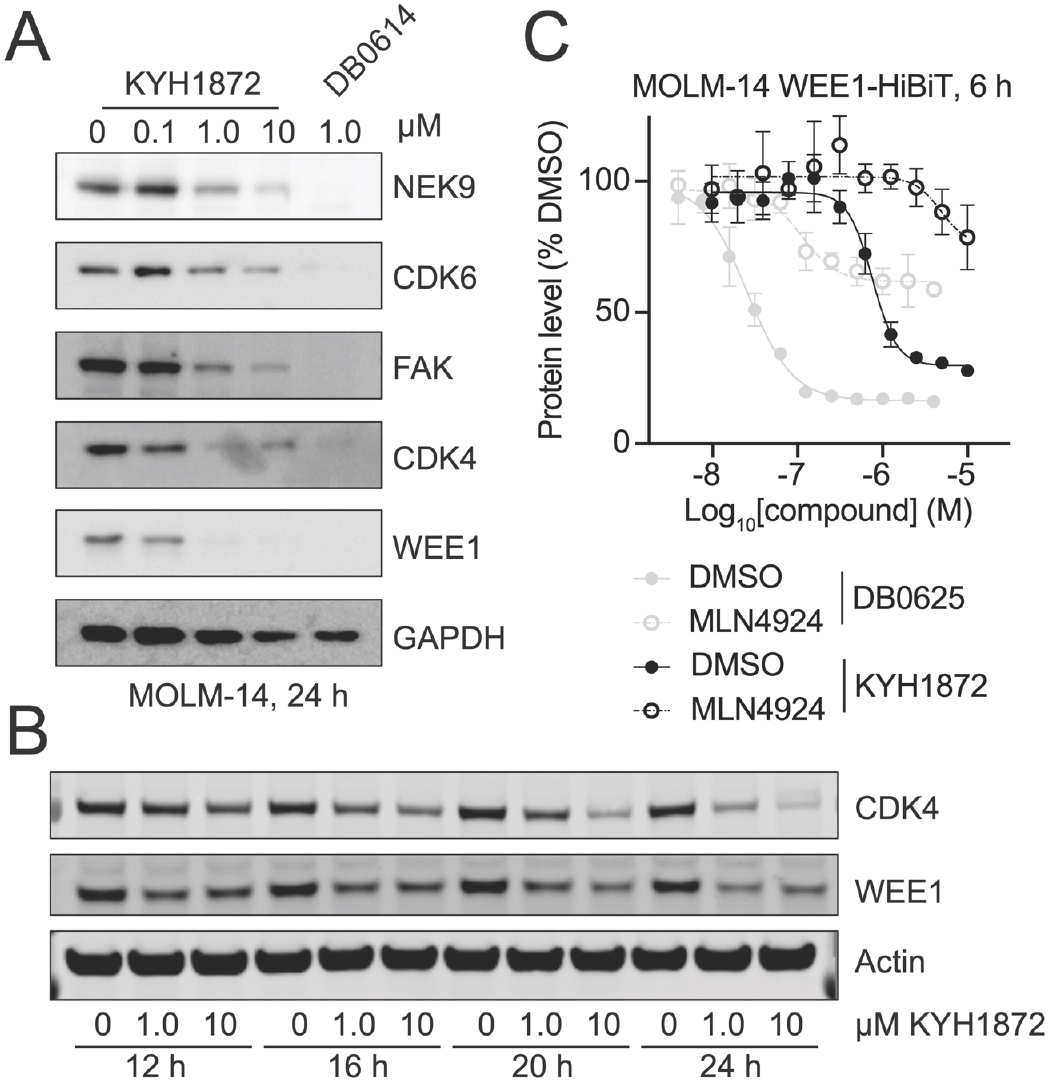
A) Kinase degradation was assessed by Western blotting for the indicated proteins after treatment of MOLM-14 cells for 24 hours with the indicated compounds. B) Time course Western blot experiment performed as in panel A. Cells were collected after the treatment times indicated below. C) HiBiT experiment showing WEE1 kinase domain protein levels at the indicated compound concentrations. DB0625 is identical to DB0614 except for the linker (*18*). The compounds have similar substrate profiles. Cells were pretreated with 1 μM MLN4924 for two hours before adding the indicated degrader compounds. Three independent replicates were made for each measurement. Error bars show +/- SD.

To determine the kinetics of kinase degradation, we carried out time-course experiments and observed degradation of representative KYH1872 targets by Western blotting. For these experiments, we selected WEE1 and CDK4, kinase substrates that were readily degraded in the 24-hour experiments described above. Degradation of WEE1 and CDK4 was observable as early as 12 hours after treatment and continued until 24 hours after treatment, when the experiment was stopped (Figure 2B). Quantitative Western blotting indicated half maximal degradation of several kinase substrates of at approximately 1 μM KYH1872 in a 24-hour experiment (Table 2). We used HiBiT assays (*20*), which can provide more sensitive detection of early degradation events than Western blotting, to detect degradation of kinases six hours after treating MOLM-14 cells with KYH1872. Half maximal kinase degradation occurred at KYH1872 concentrations comparable to those derived from Western blotting experiments (Figure 2C, Table 3).

**Table 2.** Degradation efficiencies for the indicated kinases as determined by Western blotting. DC_50_ values, or the concentration of compound at which 50% target degradation is observed, are expressed as log_10_[KYH1872] (M). MOLM-14 cells were treated with the indicated compounds for 24 hours.

|  | CDK4 | CDK6 | WEE1 | NEK9 | FAK |
| --- | --- | --- | --- | --- | --- |
| KYH1872 | -6.34 | -5.97 | -6.52 | -6.00 | -6.03 |

**Table 3.** Degradation efficiencies for the indicated kinases as determined by HiBit assays. DC_50_ values are expressed as log_10_[compound] (M). D_max_ values, or the maximum total fraction of the target degraded, is given as a fraction in parentheses. MOLM-14 cells were treated with the indicated compounds for six hours. The assays are described in the Supporting Information.

| Compound | CDK4 | CDK6 | WEE1-KD* |
| --- | --- | --- | --- |
| DB0625 | -7.17 (0.69) | -7.40 (0.55) | -7.28 (0.82) |
| KYH1872 | -6.27 (0.31) | -6.46 (0.15) | -6.20 (0.73) |
| KYH2011 | n.d. | n.d. | n.d. |

\* WEE1 kinase domain (residues 290-576)
n.d. – not determined (no degradation)

The promiscuous kinase inhibitor YHJ1039 and the previously reported CRBN-targeted PROTAC based on this molecule, DB0614, are cytotoxic (*18*). To distinguish between non-specific kinase destruction as a secondary effect of cell cytotoxicity and kinase degradation that is a direct consequence of ubiquitin transfer to targets specified by KYH1872, we pre-treated cells with the Neddylation Activating Enzyme (NAE) inhibitor, MLN4924, for 2 hours before treating with KYH1872 for 24 hours. (*21, 22*). NAE inhibition, which inactivates cullin-dependent ubiquitin transfer to substrate proteins, rescued the degradation of CDK4 and WEE1 in cells treated for 24 hours with 1 μM KYH1872 (Figure 3A), confirming that this assay reports on ubiquitin-dependent induced protein degradation. MLN4924 also prevented WEE1 and CDK4 degradation in a six-hour experiment carried out in the HiBiT system described above (Figure 2C, S3A). Consistent with these findings, co-treatment with increasing concentrations of MLN4924 blunted the negative effect of KYH1872 on cell proliferation (Figure S3B), indicating that protein degradation is at least partially responsible for the anti-proliferative effects of KYH1872. RT-PCR showed that WEE1 and CDK4 RNA levels were not changed in cells treated with 1 μM KYH1872 for 24 hours (Figure S3C).

**Figure 3.**
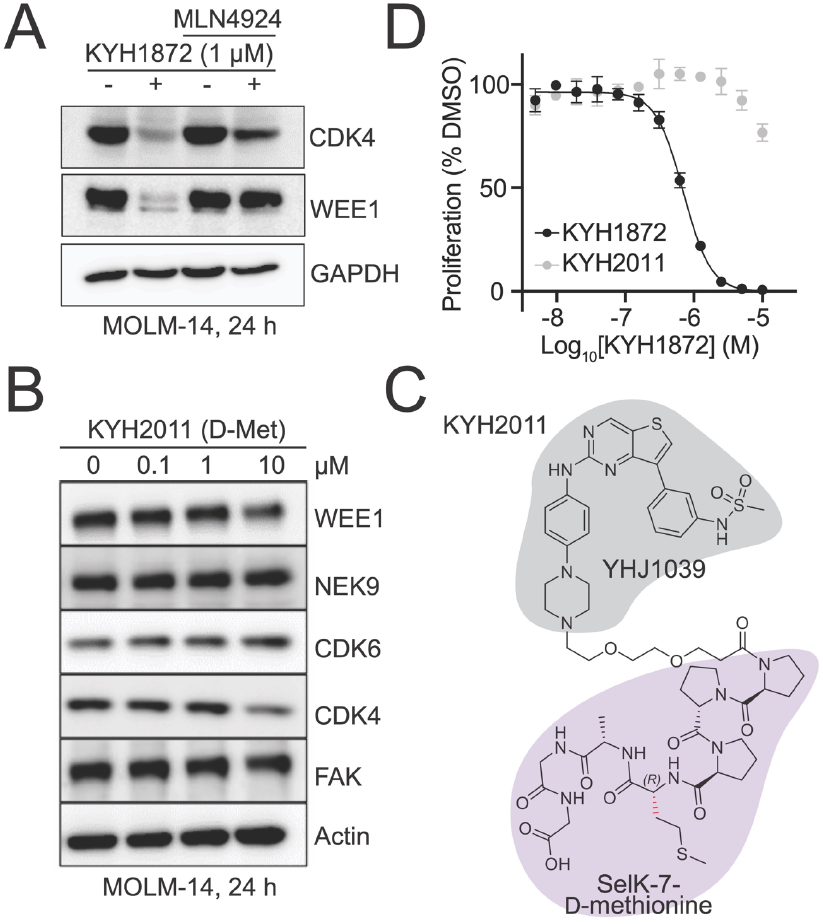
A) Kinase degradation after treatment with KYH1872 in the presence or absence of the NAE inhibitor MLN4924. MOLM-14 cells were treated for 24 hours with the indicated compounds. B) Structure of the KLHDC2-interacting fragment of the KYH2011 compound. The distinguishing D-Methionine residue is colored in red. C) Kinase degradation was assessed by Western blotting for the indicated proteins after treatment of MOLM-14 cells for 24 hours with the KYH2011 compound. D) Cell measurements after MOLM-14 cells were treated for 72 hours with the indicated compounds. Error bars show +/- SD.

To further study the mechanism of KYH1872-mediated kinase degradation, we synthesized PROTAC molecules with altered SelK fragments. KYH2011 is identical to KYH1872 except that the alpha carbon stereocenter of the methionine at position four in the SelK peptide is reversed (L-Met to D-Met; Figure 3B). KYH2011-KLHDC2 binding was more than 10-fold weaker than KYH1872-KLHDC2 binding in biochemical experiments (Table 1), and KYH2011 did not cause kinase degradation or inhibit cell proliferation (Figure 3C-D). We also synthesized KYH1996, in which the methionine residue at position four has been substituted by *O*-methyl homoserine (Figure S4A-B). KYH1996 was indistinguishable from KYH1872 in the KLHDC2 binding assay; both have dissociation constants tighter than the limit of detection for the assay, which we estimate to be 20 nM. Nevertheless, KYH1996 displayed slightly weaker activity than KYH1872 in cellular kinase degradation experiments and cell proliferation assays (Table 1). Therefore, tight binding to KLHDC2 is required for the degradation activity of KYH1872.

To determine how broadly useful KLHDC2 might be as an E3 ligase for targeted protein degradation, we assessed kinase degradation by Western blotting in a panel of cancer cell lines (Figure S5A). MOLM-14, MDA-MB-231, and MV4-11 cancer cells displayed WEE1 and CDK4 degradation in response to treatment with KYH1872 at 1 μM, while other cell lines tested did not. To determine whether variable KYH1872 responsiveness could be explained by different KLHDC2 protein levels, we assessed its expression by Western blot (Figure S5B). Nearly all tested cancer cell lines express KLHDC2 at detectable levels, and MOLM-14 cells do not overexpress this protein.

## Conclusion

The results presented here indicate that KLHDC2 is ubiquitin E3 ligase useful for the design of new PROTAC molecules. A seven amino acid fragment of the endogenous KLHDC2 substrate, SelK, suffices to redirect KLHDC2 activity towards proteins of interest. These are then targeted for proteolysis. Linking this peptide with a promiscuous kinase inhibitor demonstrated the utility of this approach. Rescue experiments conducted with a chemical inhibitor of cullin-dependent ubiquitin ligation indicated that kinase degradation depends on ubiquitination. Cellular inactivity of a minimally altered PROTAC molecule with weakened KLHDC2 binding affinity in biochemical experiments (KYH2011) indicated that KLHDC2 is the ubiquitin ligase responsible for degradation.

Dependence of KYH1872 on KLHDC2 affinity suggests but does not prove that KLHDC2 is the sole E3 ubiquitin ligase required for the degradation activity of this compound. Indeed, ubiquitin E3 ligase members of the C-end degradation pathway share similar but non-overlapping recognition motifs (*12*). Stringent proof that KLHDC2 is the sole responsible ubiquitin E3 ligase will require experiments in engineered cell lines lacking this factor. Thus far, we have failed to generate these cell lines. That KLHDC2 expression level does not appear to predict KYH1872-mediated kinase degradation indicates that low levels of this E3 ligase are sufficient for PROTAC-mediated targeting in susceptible cell lines, a notion consistent with the anticipated catalytic mechanism of action of the lead degrader and with the tight biochemical affinity between compound and ligase.

The most potent peptide-based KLHDC2-targeting degrader reported here (KYH1872) is substantially less potent than CRBN-targeting compounds using the same promiscuous kinase ligand (*18*). We suspect this is due to poor cellular permeability of the SelK peptide. Given this, it is surprising that KYH1872 is more active than KYH1886. Binding assays suggest a threshold dissociation constant of <20 nM (PROTAC-KLHDC2 binding), above which KLHDC2-targeting PROTACs are inactive. Extremely poor cellular permeability coupled with tight KLHDC2 binding (a low equilibrium binding constant) once inside cells could explain the very different responses of MOLM-14 cells to KYH1872 and KYH2011; inefficient cell penetration could produce cellular concentrations of the free drug that are at or below the dissociation constant.

It remains to be seen whether optimization of the SelK peptide from KYH1872 to improve its cell-permeability will result in PROTACs that relax the requirement for tight KLHDC2 binding. Such peptide modifications could include removal of amide bonds, cyclization, or modification of amino acid side chains. Introduction of a covalent-acting chemical group could also improve the activity of the compound, though given tight SelK-KLHDC2 binding, this is a secondary priority. Substantial improvements could obviate the need for a concerted small molecule screening campaign targeting KLHDC2.

Extensive investigation of CRBN- and VHL-based PROTACs has demonstrated the importance of induced direct contact between E3 ligase and neo-substrate proteins for the most potent degraders (*23, 24*). Whether similar adventitious KLHDC2-neo-substrate interactions can drive the potency of KLHDC2-based degrader molecules will be of great interest.

## Acknowledgments

This work was supported by the following grants: R01 CA218278 from NIH, the Basic Science Research Program (NRF-2021R1A2C3011992) from the National Research Foundation in Korea, the Brain Korea 21 Project, and the KU-KIST Graduate School of Converging Science and Technology Program.

