## Supplemental Information for "Targeted Kinase Degradation via the KLHDC2 Ubiquitin E3 Ligase"

**Supporting Information**

**Table of Contents:**

Supplementary Figures…………………………………………………………2-6

Chemistry Methods……………………………………………………………3-19

Biology and Biochemistry Methods……………………………………..…20-21

$^{1}H$ and $^{13}C$ NMR spectra of intermediates…………………………………...22-37

$^{1}H$, $^{13}C$ NMR, HPLC traces, and ESI-HRMS spectra of final compounds.….38-49

Supplemental References……………………………………..……………50

**Supplementary Figures**

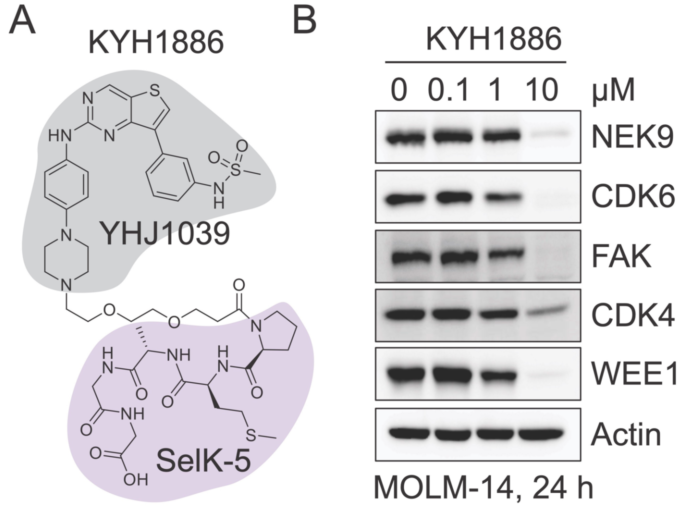


**Figure S1.** A) Structure of the KYH1886 PROTAC. B) Western blot result showing kinase degradation upon treatment of MOLM-14 cells with KYH1886 for 24 hours at the indicated concentrations.

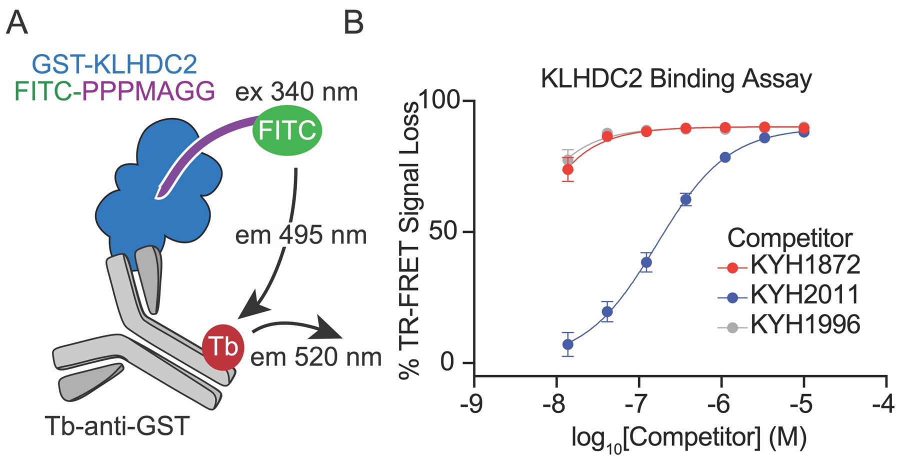


**Figure S2.** A) Schematic showing the biochemical binding assay principle, which is based on TR-FRET (Time-Resolved Förster Resonance Energy Transfer). Upon incubation with compounds that engage KLHDC2, the FITC-labeled SelK peptide (purple) is displaced from recombinant GST-KLHDC2. B) KLHDC2 biochemical binding assay showing the ability of the indicated compounds to compete with FITC-SelK for KLHDC2 binding. Error bars show +/- SD.

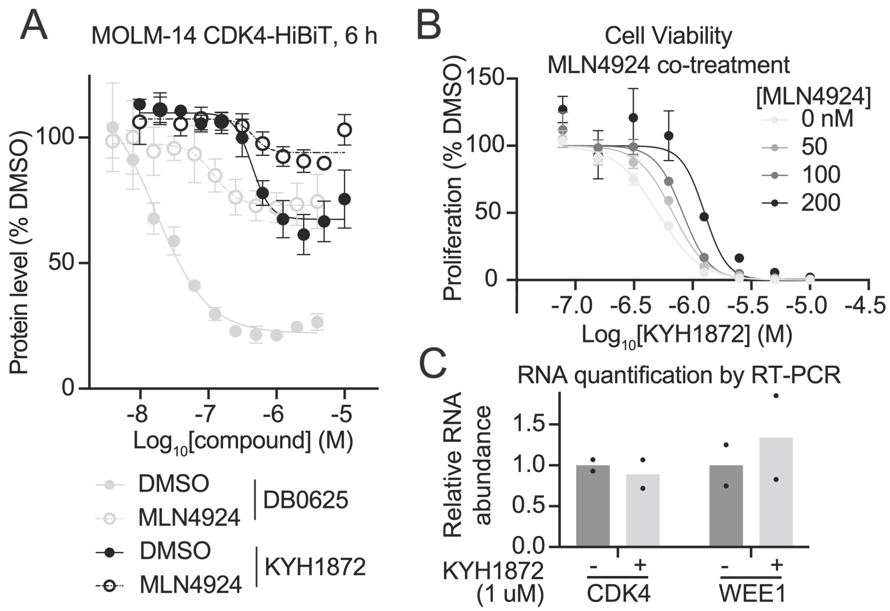


**Figure S3.** A) HiBiT experiment showing levels of CDK4-HiBiT protein after treatment with the indicated compounds. Cells were pretreated for two hours with either DMSO or 1 μM MLN4924. B) Cell viability experiment showing the effects of increasing doses of MLN4924. Maximal cell viability was calculated separately for each MLN4924 dose (right). Error bars show +/- SD. C) RNA levels were determined by RT-PCR. GAPDH- and DMSO-normalized values from two biological replicates are shown with measurements from independent experiments plotted as dots. RNA levels were determined by densitometric quantification of RT-PCR band intensity.

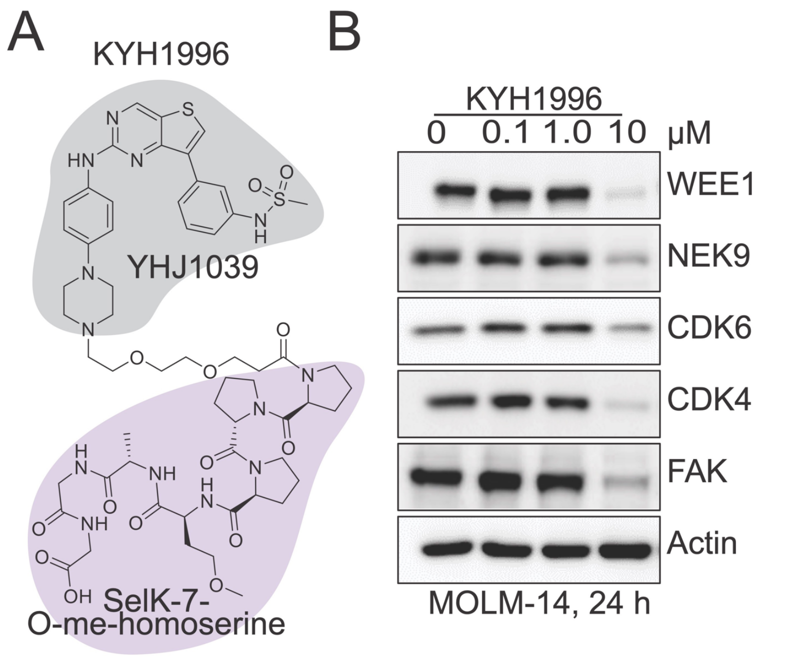


**Figure S4.** A) Structure of the KYH1996 compound. B) Western blot showing kinase levels in MOLM-14 cells treated with the indicated concentrations of KYH1996.

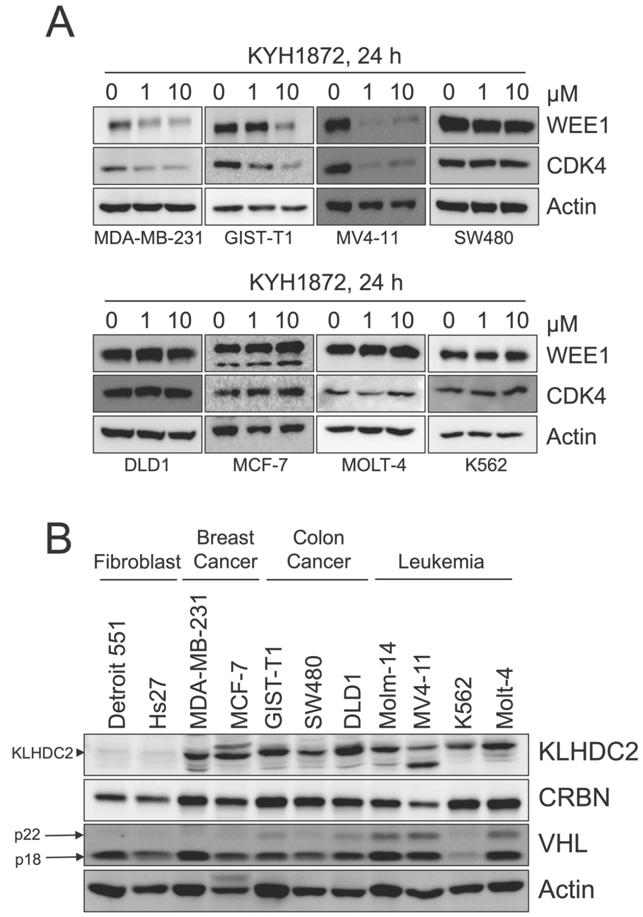


**Figure S5.** A) Western blots showing kinase degradation (WEE1 and CDK4) in the indicated cell lines treated for 24 h with KYH1872. B) KLHDC2, VHL, and CRBN expression levels assessed by Western blot in the indicated cell lines. The arrow (left) points to the KLHDC2 band.

*Chemistry Materials and Methods*

**General information.** Unless otherwise described, all commercial reagents and solvents were purchased from commercial suppliers and used without further purification. All reactions were performed under a $N_2$ atmosphere in flame-dried glassware. Reactions were monitored by using TLC with 0.25 mm E. Merck pre-coated silica gel plates (60 F254). Reaction progress was monitored by using TLC analysis using a UV lamp, ninhydrin, or p-anisaldehyde stain for detection purposes. All solvents were purified by using standard techniques. Purification of reaction products was carried out by using silica gel column chromatography with Kieselgel 60 Art. 9385 (230–400 mesh). Purities of all compounds were ≥ 95%. Mass spectra and purities of all compounds were assessed using LC/mass spec analysis with Waters LCMS system (Waters 2998 Photodiode Array Detector, Waters 3100 Mass Detector, Waters SFO System Fluidics Organizer, Water 2545 Binary Gradient Module, Waters Reagent Manager, and Waters 2767 Sample Manager) using SunFireTM C18 column (4.6 × 50 mm, 5 $\mu$m particle size): solvent gradient = 30% B at 0.00 min, 100% B at 7.00 min, 100% B at 8.50 min, 30% B at 8.51 min, 30% B at 10.00 min. Solvent A = 0.1% formic acid in $H_2O$; Solvent B = 0.1% formic acid in MeOH; flow rate = 0.8 mL/min. $^1H$ and $^{13}C$ NMR spectra were obtained using Bruker 400 MHz FT-NMR (400 MHz for $^1H$, and 100 MHz for $^{13}C$) spectrometer. Standard abbreviations are used for denoting the signal multiplicities.

Pathways for synthesis of PROTAC molecules having a broad-spectrum kinase inhibitor (YHJ1039) fused with KLHDC2 substrate peptides are outlined in Schemes 1-7. The synthetic route employed to prepare the linker-attached warhead **6**, given in Scheme 1, begins with the treatment of commercially available acrylate **l** and 2-(2-hydroxyethoxy)ethanol with Triton B, followed by iodination using $I_2$ to yield the iodide **3** (85%). The kinase inhibitor **4** prepared as previously reported was subjected to alkylation with iodide **3** using NaI and DIPEA, followed by hydrolysis of the *tert*-butyl ester to yield the key intermediate **6** (*1*).

**Scheme 1.** Synthesis of Linker-attached Warhead **6**.

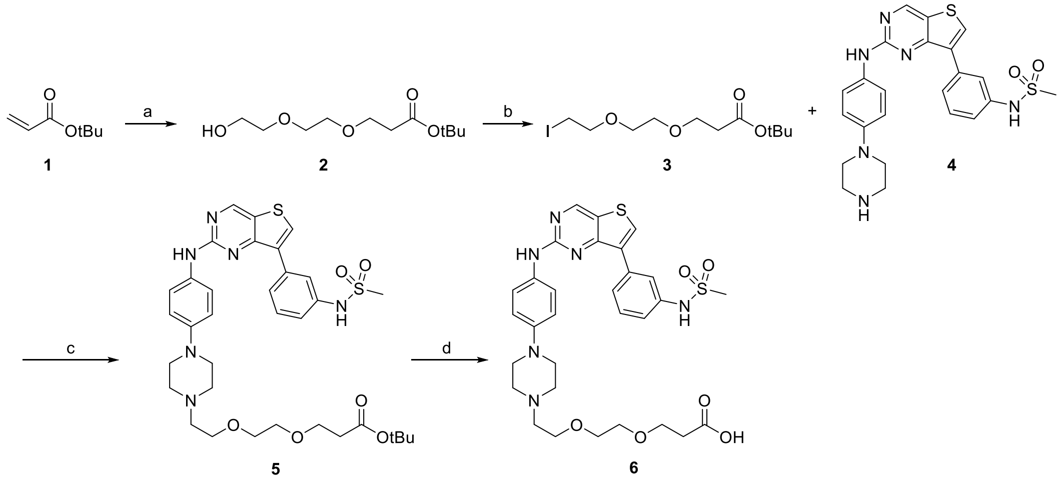


Reagent and condition: (a) 2-(2-hydroxyethoxy)ethanol, Triton B, ACN, rt, 12 h, 44%; (b) $I_2$, triphenylphosphine, imidazole, DCM, rt, 12 h, 85%; (c) NaI, DIPEA, DMF, 60 °C, 3 h, 43%; (d) 4 M HCl in dioxane, DCM, 0°C to rt, 6 h, 96%.

Peptide fragments **10** and **17-19** were prepared by the synthetic strategy described in Schemes 2-3. Amide coupling reactions using 1-ethyl-3-(3-dimethylaminopropyl carbodiimide hydrochloride and 1-hydroxybenzotriazole carried out on the commercially available amino acid **7** with glycine methyl ester hydrochloride yielded the dipeptide **8** (87%). The hydrolysis of the methyl ester, followed by another amide coupling reaction formed **9** (91%), which upon removal of *tert*-butyloxycarbonyl group formed the tripeptide **10** (91%). The same synthetic route as for **9** by using the commercially available amino acid **11** and *L*-proline methyl ester hydrochloride was employed to synthesize the tripeptide **13** (79%), which upon hydrolysis of methyl ester and amide coupling with the corresponding amino acid methyl ester hydrochloride formed **17-19** (75-82%).

**Scheme 2.** Synthesis of Tripeptide Fragment **10**.

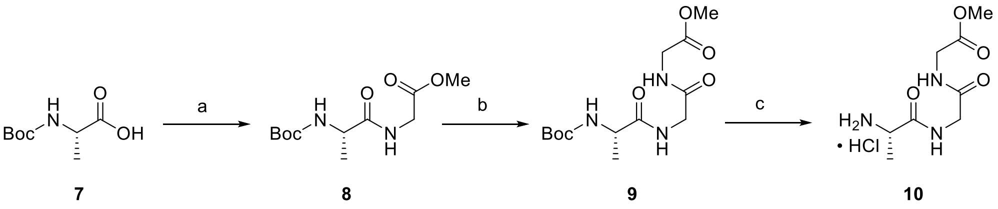


Reagent and condition: (a) glycine methyl ester hydrochloride, 1-ethyl-3-(3-dimethylaminopropyl carbodiimide hydrochloride, 1-hydroxybenzotriazole, DIPEA, DCM, rt, 3 h, 87%; (b) i) LiOH, THF/$H_2O$ (1:1), 0 °C to rt, 3 h; ii) glycine methyl ester hydrochloride, 1-ethyl-3-(3-dimethylaminopropyl carbodiimide hydrochloride, 1-hydroxybenzotriazole, DIPEA, DCM, rt, 3 h, 91% over two steps; (c) 4 M HCl in dioxane, DCM, 0°C to rt, 3 h, 91%.

**Scheme 3.** Synthesis of Tetrapeptide Fragments **17-19** .

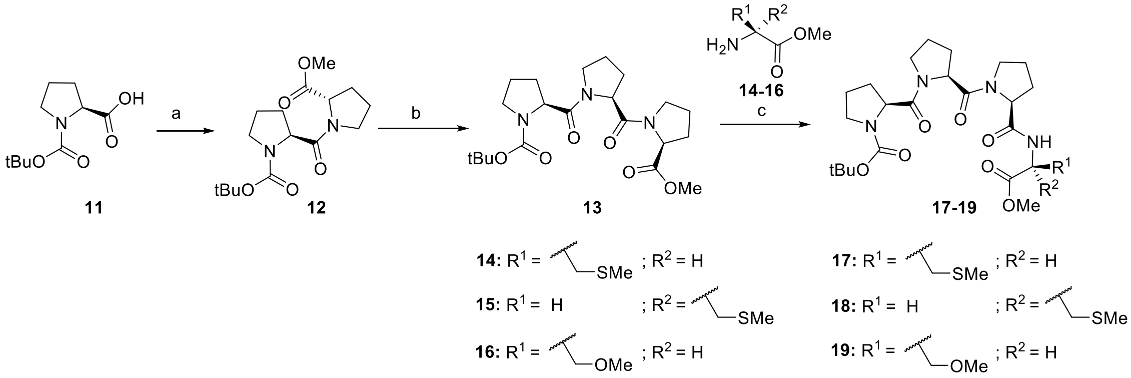


Reagent and condition: (a) amino acid methyl ester hydrochloride, 1-ethyl-3-(3-dimethylaminopropyl)carbodiimide hydrochloride, 1-hydroxybenzotriazole, DIPEA, DCM, rt, 3 h, 86%; (b) i) LiOH, THF/$H_2O$ (1:1), 0 °C to rt, 3 h; ii) *L*-proline methyl ester hydrochloride, 1-ethyl-3-(3-dimethylaminopropyl carbodiimide hydrochloride, 1-hydroxybenzotriazole, DIPEA, DCM, rt, 3 h, 79% over two steps; (c) i) LiOH, THF/$H_2O$ (1:1), 0 °C to rt, 3 h; ii) corresponding amino acid methyl ester hydrochloride, 1-ethyl-3-(3-dimethylaminopropyl carbodiimide hydrochloride, 1-hydroxybenzotriazole, DIPEA, DCM, rt, 3 h, 75-82% over two steps.

The synthesis of penta- and heptapeptide KLHDC2 binders is shown in Schemes 4-5. Amide coupling reactions using 1-ethyl-3-(3-dimethylaminopropyl carbodiimide hydrochloride and 1-hydroxybenzotriazole carried out on the commercially available amino acid **11** with *L*-methionine methyl ester hydrochloride yielded the dipeptide **20** (65%), which upon hydrolysis of the methyl ester and amide coupling reaction with the tripeptide **10** formed the pentapeptide KLHDC2 binder **21** (32%). The hydrolysis of ester moieties of tetrapeptides **17-19**, followed by amide coupling reaction with the tripeptide **10** generated heptapeptide KLHDC2 binders **22-24** (30-45%).

**Scheme 4.** Synthesis of Pentapeptide KLHDC2 Binder **21**.

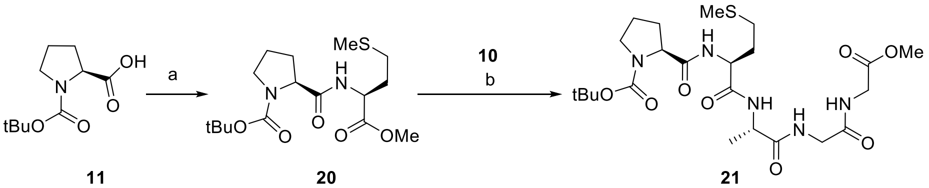


**11** **20** **21**

Reagent and condition: (a) amino acid methyl ester hydrochloride, 1-Ethyl-3-(3-dimethylaminopropyl)carbodiimide hydrochloride, 1-hydroxybenzotriazole, DIPEA, DCM, rt, 3 h, 65%; (b) i) LiOH, THF/$H_2O$ (1:1), 0 °C to rt, 3 h; ii) **10**, 1-Ethyl-3-(3-dimethylaminopropyl carbodiimide hydrochloride, 1-hydroxybenzotriazole, DIPEA, DCM, rt, 3 h, 32% over two steps.

**Scheme 5.** Synthesis of Heptapeptide KLHDC2 Binders **22-24** .

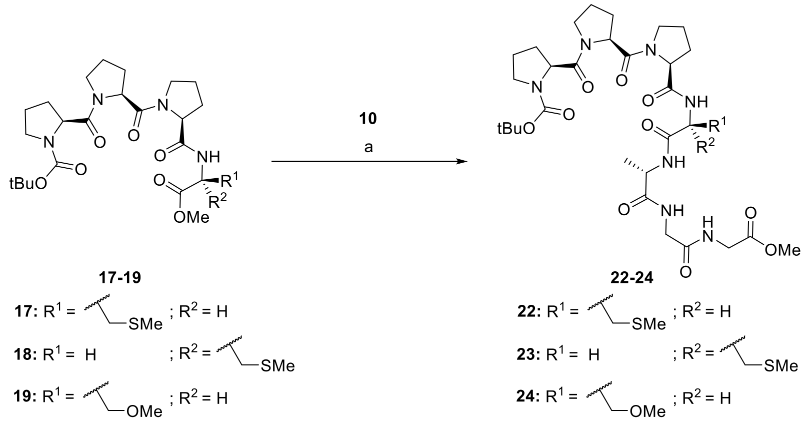


Reagent and condition: (a) i) LiOH, THF/$H_2O$ (1:1), 0 °C to rt, 3 h; ii) amino acid methyl ester hydrochloride, 1-Ethyl-3-(3-dimethylaminopropyl carbodiimide hydrochloride, 1-hydroxybenzotriazole, DIPEA, DCM, rt, 3 h, 30-45% over two steps.

The synthetic routes towards the various KLHDC2 PROTAC molecules are shown in Schemes 6-7. The peptide key intermediates **21** and **22-24** were treated with HCl (4.0 M in 1,4-dioxane) to deprotect *tert*-butyloxycarbonyl group, followed by amide coupling reactions with the linker-attached warhead **6** using 1-propanephosphonic anhydride (50% w/w in EA). The synthesis involved, as a final step, removal of the methyl ester to yield KLHDC2 PROTAC molecules **25-28** (33-57%).

**Scheme 6.** Synthesis of Pentapeptide KLHDC2 PROTAC **25**.

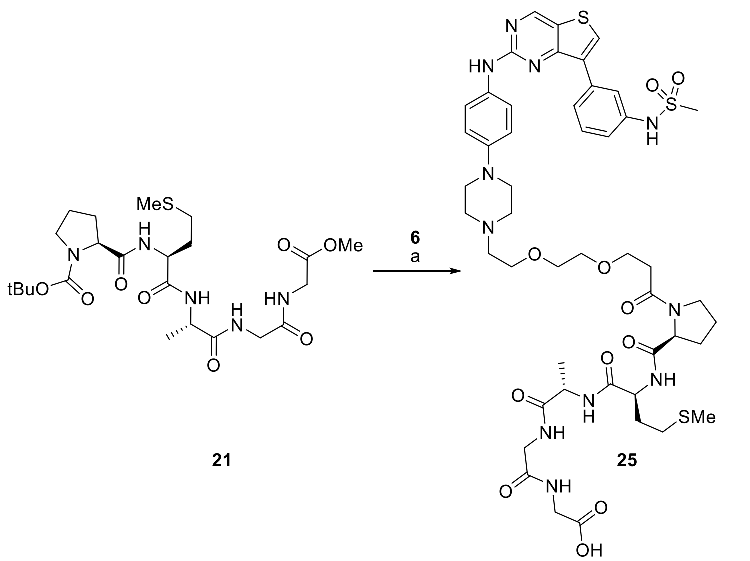


Reagent and condition: (a) i) 4 M HCl in dioxane, DCM, 0°C to rt, 3 h ii) **6**, triethylamine, 1-propanephosphonic anhydride (50% w/w in EA), DMF, -10 °C, 0.5 h; iii) LiOH, THF/$H_2O$ (1:1), -10 °C, 0.5 h, 43%.

**Scheme 7.** Synthesis of Heptapeptide KLHDC2 PROTACs **26-28**.

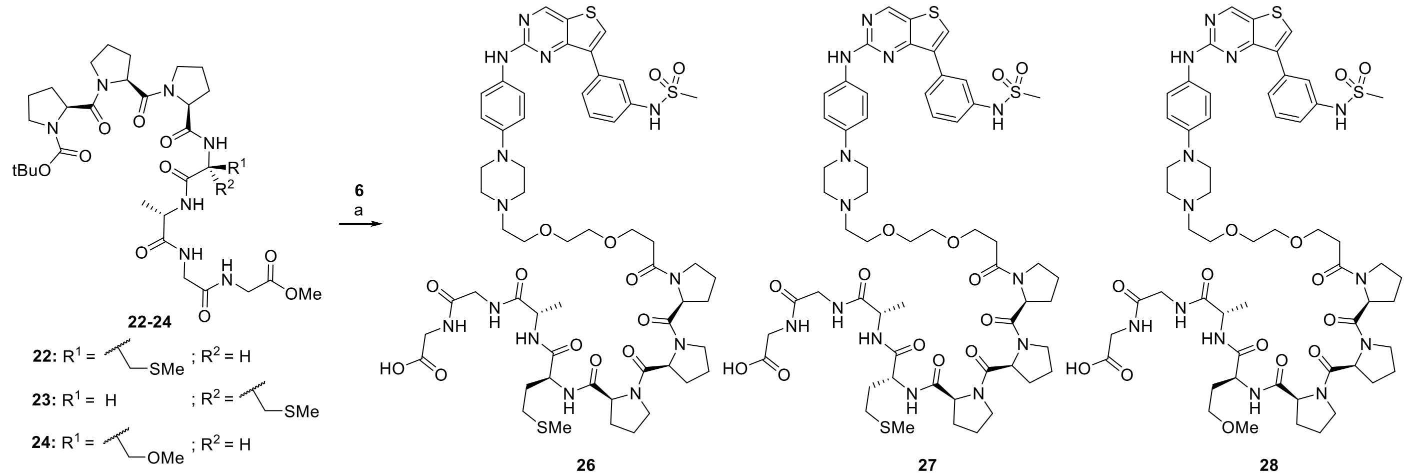


Reagent and condition: (a) i) 4 M HCl in dioxane, DCM, 0°C to rt, 3 h ii) **6**, triethylamine, 1-propanephosphonic anhydride (50% w/w in EA), DMF, -10 °C, 0.5 h; iii) LiOH, THF/$H_2O$ (1:1), -10 °C, 0.5 h, 33-57%.

*tert*-butyl 3-(2-(2-hydroxyethoxy)ethoxy)propanoate **(2)**.

To a solution of *tert*-butyl acrylate (3 ml, 20.7 mmol) and 2-(2-hydroxyethoxy)ethanol (2 mL, 20.7 mmol) in acetonitrile (30 mL) was added Triton B (0.3 mL, 0.75 mmol) and stirred at room temperature for 12 h. The reaction mixture was concentrated and diluted with ethyl acetate and water. The organic phase was washed with $NaHCO_3$ solution and brine, dried over $Na_2SO_4$, filtered, and concentrated. The residue was subjected to chromatography on silica gel (30% to 50% EA/hexane) to afford **2** (2.12 g, 44%).$^1$H NMR (400 MHz, $CDCl_3$): $\delta$ 3.75 – 3.68 (m, 4H), 3.66 – 3.56 (m, 6H), 2.57 – 2.44 (m, 3H), 1.44 (s, 9H); $^{13}$C NMR (101 MHz, $CDCl_3$) $\delta$ 171.07, 80.81, 72.59, 70.46, 70.44, 66.94, 61.88, 36.27, 28.19. LRMS (ESI) *m/z* 257 $[M + H]^+$.

*tert*-butyl 3-(2-(2-iodoethoxy)ethoxy)propanoate **(3)**.
To a solution of **2** (1.00 g, 4.27 mmol) and triphenylphosphine (1.17 g, 4.50 mmol) in DCM (30 mL), imidazole (363 mg, 5.30 mmol) was added followed by $I_2$ (1.19 g, 4.70 mmol). The mixture was stirred at room temperature for 12 h. The reaction mixture was quenched by addition of saturated aqueous sodium thiosulfate solution and extracted with DCM. The organic phase was washed with brine, dried over $Na_2SO_4$, filtered, and concentrated. The residue was subjected to chromatography on silica gel (0% to 20% EA/hexane) to afford **3** (1.25 g, 85%) as yellow oil. $^1$H NMR (400 MHz, $CDCl_3$): $\delta$ 3.74 (t, $J$ = 5.9 Hz, 2H), 3.71 (t, $J$ = 5.5 Hz, 2H), 3.66 – 3.59 (m, 4H), 3.24 (t, $J$ = 6.9 Hz, 2H), 2.50 (t, $J$ = 6.5 Hz, 2H), 1.44 (s, 9H); $^{13}$C NMR (101 MHz, $CDCl_3$) $\delta$ 171.00, 80.68, 72.07, 70.46, 70.24, 67.06, 36.34, 28.22, 3.03. LRMS (ESI) *m/z* 367 [M + Na]$^+$.

*tert*-butyl 3-(2-(2-(4-(4-((7-(3-(methylsulfonamido)phenyl)thieno[3,2-d]pyrimidin-2-yl)amino)phenyl)piperazin-1-yl)ethoxy)ethoxy)propanoate **(5)**.
To a solution of **4** (335 mg, 0.70 mmol) in DMF (5.0 mL) was added **3** (288 mg, 0.84 mmol), NaI (524 mg, 3.49 mmol), DIPEA (0.49 ml, 2.79 mmol) at room temperature. The reaction mixture was then stirred at 60 $^o$C for 3 h, quenched with water and diluted with EtOAc. The organic layer was washed with brine, dried over $Na_2SO_4$, filtered, and concentrated under reduced pressure. The residue was purified by flash column chromatography on silica gel (50% to 70% THF/hexane) to afford **5** (210 mg, 43%) as a yellow solid.
$^1$H NMR (400 MHz, DMSO-$d_6$): $\delta$ 9.84 (s, 1H), 9.44 (s, 1H), 9.16 (s, 1H), 8.45 (s, 1H), 7.83 (t, $J$ = 1.74, 2H), 7.82 – 7.79 (m, 1H), 7.70 (d, $J$ = 9.1, 2H), 7.48 (t, $J$ = 7.9, 1H), 7.27 (ddd, $J$ = 0.9, 2.2, 8.1 1H), 6.91 (d, $J$ = 9.1, 2H), 3.60 (t, $J$ = 6.2, 2H), 3.54 (t, $J$ = 5.9, 2H), 3.50 (s, 4H), 3.06 (t, $J$ = 4.4, 4H), 3.02 (s, 3H), 2.56 (t, $J$ = 4.4, 4H), 2.53 – 2.50 (m, 2H), 2.42 (t, $J$ = 6.2, 2H), 1.40 (s, 9H); $^{13}$C NMR (101 MHz, DMSO-$d_6$) $\delta$ 170.47, 158.41, 157.95, 154.00, 146.12, 138.57, 134.60, 134.38, 133.69, 132.73, 129.34, 123.81, 122.19, 120.07, 119.97, 119.10, 115.89, 79.75, 69.69, 69.66, 68.43, 66.27, 57.31, 53.28, 49.07, 35.89, 27.79. LRMS (ESI) *m/z* 697 [M + H]$^+$.

3-(2-(2-(4-(4-((7-(3-(methylsulfonamido)phenyl)thieno[3,2-d]pyrimidin-2-yl)amino)phenyl)piperazin-1-yl)ethoxy)ethoxy)propanoic acid **(6)**.
To a solution of **5** (500 mg, 0.72 mmol) in DCM (1.44 mL) was added HCl (4.0 M in 1,4-dioxane, 1.80 mL) at 0 °C. After stirring at room temperature for 6 h, the solution was concentrated under reduced pressure. The residue was solidified by swirling in DCM/diethyl ether (1:4) and separation of the formed solid by filtration followed by drying yielded **6** (443 mg, 0.69 mmol, 96%) as a yellow solid that was used for next reaction without any further purification. LRMS (ESI) *m/z* 641 [M + H]$^+$.

**General Procedure A for the Synthesis of Dipeptides.**
To a solution of Boc-protected amino acid (1.0 equiv) in DCM (0.2 M) was added 1-ethyl-3-(3-dimethylaminopropyl carbodiimide hydrochloride (1.5 equiv) and 1-hydroxybenzotriazole hydrate (1.5 equiv). The mixture was stirred for 5 min, treated slowly with a solution of amino acid methyl ester hydrochloride (1.2 equiv) and DIPEA (4 equiv) in DCM (0.4 M). The reaction mixture was stirred at room temperature for 2 h, quenched by addition of ice water, and extracted with IPA/CHCl3 (1:4). The organic layer was washed with $NH_4Cl$ solution and $NaHCO_3$ solution, dried over $Na_2SO_4$, filtered, and concentrated under reduced pressure. The residues were subjected to flash column chromatography on silica gel.

**General Procedure B for the Hydrolysis of Ester Group and Amide Bond Formation.**
To a solution of Boc-protected amino acid methyl ester (1.0 equiv) in THF/$H_2O$ (1:1, 0.1 M) was added aqueous LiOH solution (3.0 equiv) at 0 °C. The reaction mixture was stirred at room temperature for 2 h and pH was adjusted to 2 by addition of 1 N aqueous HCl solution. The mixture was then extracted with IPA/$CHCl_3$ (1:4), dried over $Na_2SO_4$, filtered, and concentrated under reduced pressure. To the solution of crude acid in DCM (0.2 M) was added 1-ethyl-3-(3-dimethylaminopropyl carbodiimide hydrochloride (1.5 equiv) and 1-hydroxybenzotriazole hydrate (1.5 equiv). The mixture was stirred for 5 min, treated slowly with a solution of corresponding amino acid methyl ester hydrochloride (1.2 equiv) and DIPEA (4 equiv) in DCM (0.4 M). The reaction mixture was stirred at room temperature for 2 h, quenched by addition of ice water and extracted with IPA/CHCl3 (1:4). The organic layer was washed with $NH_4Cl$ solution and $NaHCO_3$ solution, dried over $Na_2SO_4$, filtered, and concentrated under reduced pressure. The residues were subjected to flash column chromatography on silica gel.

Methyl (*tert*-butoxycarbonyl)-*L*-alanylglycinate **(8)**.
(*tert*-Butoxycarbonyl)-*L*-alanine (2.00 g, 10.6 mmol) was converted to the target compound using general procedure A. The crude product was purified using flash column chromatography (0% to 5% MeOH/$CH_2Cl_2$) on silica gel to afford **8** (2.39 g, 9.2 mmol, 87%) as a pale yellow oil. $^1$H NMR (400 MHz, DMSO-$d_6$): $\delta$ 8.30 – 8.13 (m, 2H), 6.96 (d, $J$ = 7.6 Hz, 1H), 4.06 – 3.94 (m, 1H), 3.83 (ddd, $J$ = 6.56, 17.5 36.9 Hz, 2H), 3.62 (s, 3H), 1.37 (s, 9H), 1.18 (d, $J$ = 6.8 Hz, 3H); $^{13}$C NMR (100 MHz, DMSO-$d_6$): $\delta$ 173.34, 170.31, 155.04, 78.02, 51.70, 49.48, 40.54, 28.23, 18.19. LRMS (ESI) *m/z* 283 $[M + Na]^+$.

Methyl (*tert*-butoxycarbonyl)-*L*-alanylglycylglycinate **(9)**.
Compound **8** (2.39 g, 9.2 mmol) was converted to the target compound using general procedure B. The crude product was purified using flash column chromatography (5% to 10% MeOH/$CH_2Cl_2$) on silica gel to afford **9** (2.64 g, 8.33 mmol, 91%) as a pale yellow oil. $^1$H NMR (400 MHz, DMSO-$d_6$): $\delta$ 8.21 (t, $J$ = 5.57, 1H), 8.13 (t, $J$ = 5.45 Hz, 1H), 7.05 (d, $J$ = 6.95 Hz, 1H), 4.02 – 3.91 (m, 1H), 3.90 – 3.78 (m, 2H), 3.72 (d, $J$ = 5.7 Hz, 2H), 3.62 (s, 3H), 1.37 (s, 9H), 1.17 (d, $J$ = 7.12 Hz, 3H); $^{13}$C NMR (100 MHz, DMSO-$d_6$): $\delta$ 173.05, 170.16, 169.40, 155.34, 78.25, 51.76, 49.84, 41.82, 40.53, 28.23, 17.95. LRMS (ESI) *m/z* 340 $[M + Na]^+$.

Methyl *L*-alanylglycylglycinate hydrochloride **(10)**.
To a solution of **9** (1.40 g, 4.42 mmol) in DCM (4.4 mL) was added HCl (4.0 M in 1,4-dioxane, 11.0 mL) at 0 °C. After stirring at room temperature for 3 h, the solution was concentrated under reduced pressure. The residue was solidified by swirling in DCM/diethyl ether (1:4) and separation of the formed solid by filtration, followed by drying yielded **10** (1.02 g, 4.03 mmol, 91%) as a white solid that was used for next reaction without any further purification. LRMS (ESI) *m/z* 240 $[M + Na]^+$.

*tert*-Butyl (*S*)-2-((*S*)-2-(methoxycarbonyl)pyrrolidine-1-carbonyl)pyrrolidine-1-carboxylate **(12)**.
(*tert*-Butoxycarbonyl)-*L*-proline (10.0 g, 46.4 mmol) was converted to the target compound using general procedure A. The crude product was purified using flash column chromatography (0% to 50% THF/hexane) on silica gel to afford **12** (13.1 g, 40.1 mmol, 86%) as a pale yellow oil.

$^1$H NMR (400 MHz, DMSO-$d_6$): $\delta$ two rotamers 4.43 (dd, $J$ = 3.1, 8.7 Hz, minor), 4.39 (dd, $J$ = 4.3, 8.3 Hz, major, 1H), 4.35 -4.28 (m, 1H), 3.7 – 3.46 (m, 2H), 3.61 (s, major, 3H), 3.60 (s, minor), 3.35 – 3.24 (m, 2H), 2.27 – 2.06 (m, 2H), 2.00 – 1.89 (m, 2H), 1.87 – 1.68 (m, 4H), 1.36 (s, minor), 1.30 (s, major, 9H);$^{13}$C NMR (100 MHz, DMSO-$d_6$): $\delta$ two rotamers 172.34 (minor), 172.20 (major), 170.94 (major), 170.35 (minor), 153.33 (minor), 152.95 (major), 78.40 (minor), 78.30 (major), 58.33 (major), 58.32 (minor), 57.33 (minor), 57.26 (major), 51.76 (major), 51.70 (minor), 46.51 (minor), 46.37 (major), 46.23 (major), 46.16 (minor), 29.36 (major), 28.46 (minor), 28.41 (major), 28.33 (minor), 28.15 (minor), 27.92 (major), 24.65 (major), 24.63 (minor), 23.59 (minor), 23.11 (major). LRMS (ESI) *m/z* 349 $[M + Na]^+$.

*t*ert-Butyl (*S*)-2-((*S*)-2-((*S*)-2-(methoxycarbonyl)pyrrolidine-1-carbonyl)pyrrolidine-1-carbonyl)pyrrolidine-1-carboxylate **(13)**.
Compound **12** (6.00 g, 18.4 mmol) was converted to the target compound using general procedure B. The crude product was purified using flash column chromatography (30% to 50% THF/hexane) on silica gel to afford **13** (6.13 g, 14.5 mmol, 79%) as a colorless oil.
$^1$H NMR (400 MHz, DMSO-$d_6$): $\delta$ two rotamers 4.61 (dd, $J$ = 4.0, 8.0 Hz, minor), 4.58 (dd, $J$ = 4.0, 8.4 Hz, major, 1H), 4.45 -4.34 (m, 1H), 4.33 – 4.24 (m, 1H), 3.72 – 3.38 (m, 4H), 3.60 (s, 3H), 3.30 – 3.25 (m, 2H), 2.24 – 2.05 (m, 3H), 2.00 – 1.86 (m, 4H), 1.86 – 1.67 (m, 5H), 1.37 (s, minor), 1.31 (s, major, 9H); $^{13}$C NMR (100 MHz, DMSO-$d_6$): $\delta$ two rotamers 172.30, 170.41(major), 169.95 (minor), 169.83 (minor), 169.79 (major), 153.31 (minor), 153.02 (major), 78.29 (minor), 78.21 (major), 58.32 (major), 58.27 (minor), 57.35, 57.29, 51.72, 46.51 (minor), 46.41 (major), 46.36, 46.25 (major), 46.20 (minor), 29.24 (major), 28.34 (minor), 28.37, 28.17 (minor), 27.96 (major), 27.60 (major), 27.41 (minor), 24.60 , 24.34 (major), 24.33 (minor), 23.56 (minor), 23.04 (major). LRMS (ESI) *m/z* 446 $[M + Na]^+$.

Methyl *O*-methyl-*L*-homoserinate hydrochloride **(16)**.
To a solution of *N*-(*tert*-butoxycarbonyl)-*L*-homoserine (1 g, 4.3 mmol) in methanol (4.3 mL) was added trimethylsilyl chloride (2.8 mL, 21.5 mmol) slowly and stirred at room temperature for 2 h. The solution was concentrated under reduced pressure. The residue was solidified by swirling in DCM/diethyl ether (1:4) and separation of the formed solid by filtration, followed by drying yielded **16** (625 mg, 3.41 mmol, 79%) as a white solid.
$^1$H NMR (400 MHz, DMSO-$d_6$): $\delta$ 5.59 (brs, 3H), 4.02 (t, $J$ = 6.1 Hz, major, 1H), 3.73 (s, 3H), 3.44 (t, $J$ = 6. Hz, 2H), 3.22 (s, major, 3H), 2.12 – 1.98 (m, 2H); $^{13}$C NMR (100 MHz, DMSO-$d_6$): $\delta$ 169.85, 66.92, 58.03, 52.72, 49.61, 29.93. LRMS (ESI) *m/z* 148 $[M + H]^+$.

*tert*-Butyl (*S*)-2-((*S*)-2-((*S*)-2-(((*S*)-1-methoxy-4-(methylthio)-1-oxobutan-2-yl)carbamoyl)pyrrolidine-1-carbonyl)pyrrolidine-1-carbonyl)pyrrolidine-1-carboxylate **(17)**.
Compound **13** (3.80 g, 8.98 mmol) was converted to the target compound using general procedure B. The crude product was purified using flash column chromatography (30% to 50% THF/hexane) on silica gel to afford **17** (4.02 g, 7.24 mmol, 81%) as a white solid.
$^1$H NMR (400 MHz, DMSO-$d_6$) δ diastereomeric mixtures (85:10) 8.70 (dd, $J$ = 7.6, 29.8 Hz, minor), 8.24 (dd, $J$ = 4.0, 7.6 Hz, major, 1H), two rotamers 4.58 (dd, $J$ = 4.1, 8.9 Hz, minor), 4.55 (dd, $J$ = 3.7, 8.6 Hz, major, 1H), 4.47 – 4.24 (m, 3H), 3.63 (s, 3H), 3.62 – 3.39 (m, 4H), 3.32 – 3.27 (m, 2H), 2.58 – 2.43 (m, 2H), 2.33 – 2.05 (m, 3H), 2.04 (s, 3H), 1.98 – 1.67 (m, 11H), 1.38 (s, minor), 1.31 (s, major, 9H); $^{13}$C NMR (100 MHz, DMSO-$d_6$) δ two rotamers 172.19, 171.98 (minor), 171.97 (major), 170.39 (major), 169.82 (minor), 169.77 (minor), 169.61

(major), 153.31 (minor), 153.04 (major), 78.31 (minor), 78.22 (major), 58.89 (major), 58.84 (minor), 57.41 (major), 57.30 (minor), 57.36, 51.92, 50.85 (minor), 50.84 (major), 46.54, 46.50 (minor), 46.44 (major), 46.38, 30.82, 29.38, 29.24 (major), 28.33 (minor), 28.80, 28.19 (minor), 27.97 (major), 27.70 (major), 27.51 (minor), 24.46, 24.32 (major), 24.29 (minor), 23.60 (minor), 23.07 (major), 14.58. LRMS (ESI) *m/z* 577 $[M + Na]^+$.

*tert*-Butyl (*S*)-2-((*S*)-2-((*S*)-2-(((*R*)-1-methoxy-4-(methylthio)-1-oxobutan-2-yl)carbamoyl)pyrrolidine-1-carbonyl)pyrrolidine-1-carbonyl)pyrrolidine-1-carboxylate **(18)**.
Compound **13** (1.50 g, 3.55 mmol) was converted to the target compound using general procedure B. The crude product was purified using flash column chromatography (30% to 50% THF/hexane) on silica gel to afford **18** (1.61 g, 2.91 mmol, 82%) as a white solid.
$^1$H NMR (400 MHz, DMSO-$d_6$) δ diastereomeric mixtures (85:13) 8.26 (dd, *J* = 7.2, 18.7 Hz, minor), 8.05 (d, *J* = 8.2 Hz, major, 1H), two rotamers 4.57 (dd, *J* = 3.8, 8.6 Hz, minor), 4.53 (dd, *J* = 3.7, 8.3 Hz, major, 1H), 4.47 – 4.34 (m, 2H), 4.33 – 4.22 (m, 1H), 3.69 – 3.34 (m, 4H), 3.63 (s, 3H), 3.32 – 3.24 (m, 2H), 2.49 – 2.32 (m, 2H), 2.22 – 1.99 (m, 3H), 2.02 (s, 3H), 2.00 – 1.71 (m, 11H), 1.37 (s, minor), 1.31 (s, major, 9H); $^{13}$C NMR (100 MHz, DMSO-$d_6$) δ two rotamers 172.13, 171.75 (minor), 171.71 (major), 170.54 (major), 170.17 (minor), 170.01 (major), 169.92 (major), 153.33 (minor), 153.04 (major), 78.34 (minor), 78.25 (major), 59.57 (major), 59.51 (minor), 57.70, 57.39 (minor), 57.34 (major), 51.98, 50.56, 46.66 (major), 46.63 (minor), 46.58 (minor), 46.55 (major), 46.48 (minor), 46.40 (major), 30.72, 29.42, 29.31 (major), 28.38 (minor), 29.11, 28.21 (minor), 28.00 (major), 27.81 (major), 27.66 (minor), 24.48, 24.47 (major), 24.43 (minor), 23.62 (minor), 23.12 (major), 14.59. LRMS (ESI) *m/z* 577 $[M + Na]^+$.

*tert*-Butyl (*S*)-2-((*S*)-2-((*S*)-2-(((*S*)-1,4-dimethoxy-1-oxobutan-2-yl)carbamoyl)pyrrolidine-1-carbonyl)pyrrolidine-1-carbonyl)pyrrolidine-1-carboxylate **(19)**.
Compound **13** (1.00 g, 2.36 mmol) was converted to the target compound using general procedure B. The crude product was purified using flash column chromatography (30% to 50% THF/hexane) on silica gel to afford **19** (950 mg, 1.76 mmol, 75%) as a white solid.
$^1$H NMR (400 MHz, DMSO-$d_6$) δ diastereomeric mixtures (84:12) 8.67 (dd, *J* = 7.5, 13.5 Hz, minor), 8.17 (dd, *J* = 7.5, 4.4 Hz, major, 1H), two rotamers 4.58 (dd, *J* = 3.6, 8.4 Hz, minor), 4.55 (dd, *J* = 3.7, 8.5 Hz, major, 1H), 4.51 – 4.20 (m, 3H), 3.68 – 3.42 (m, 4H), 3.63 (s, 3H), 3.39 – 3.33 (m, 2H), 3.30 – 3.24 (m, 2H), 3.20 (s, 3H), 2.22 – 1.98 (m, 3H), 1.97 – 1.68 (m, 11H), 1.37 (s, minor), 1.31 (s, major, 9H); $^{13}$C NMR (100 MHz, DMSO-$d_6$) δ two rotamers 172.45, 171.85 (minor), 171.83 (major), 170.38 (major), 169.81 (minor), 169.78 (minor), 169.63 (major), 153.31 (minor), 153.03 (major), 78.30 (minor), 78.21 (major), 67.86, 58.84 (major), 58.79 (minor), 57.88, 57.40 (minor), 57.30 (major), 57.35, 51.81, 49.12 (minor), 49.11 (major), 46.53, 46.49 (minor), 46.44 (major), 46.37, 31.07, 29.24 (major), 28.32 (minor), 28.81, 28.18 (minor), 27.97 (major), 27.71 (major), 27.52 (minor), 24.41, 24.34 (major), 24.30 (minor), 23.59 (minor), 23.06 (major). LRMS (ESI) *m/z* 561 $[M + Na]^+$.

*tert*-Butyl (*S*)-2-(((*S*)-1-methoxy-4-(methylthio)-1-oxobutan-2-yl)carbamoyl)pyrrolidine-1-carboxylate **(20)**.
(*tert*-Butoxycarbonyl)-*L*-proline (2.00 g, 9.30 mmol) was converted to the target compound using general procedure A. The crude product was purified using flash column chromatography (0% to 50% THF/hexane) on silica gel to afford **20** (2.19 g, 6.08 mmol, 65%) as a pale yellow oil.

$^{1}H$ NMR (400 MHz, DMSO-$d_6$): $\delta$ two rotamers 8.27 (d, $J$ = 7.5 Hz, major, 1H), 8.23 (d, $J$ = 7.3 Hz, minor), 4.45 – 4.35 (m, 1H), 4.17 – 4.07 (m, 1H), 3.63 (s, 3H), 3.41 – 3.34 (m, 1H), 3.30 – 3.22 (m, 1H), 2.60 – 2.40 (m, 2H), 2.18 – 2.07 (m, 1H), 2.03 (s, 3H), 2.00 – 1.85 (m, 2H), 1.84 – 1.72 (m, 3H), 1.38 (s, minor), 1.33 (s, major 9H); $^{13}C$ NMR (100 MHz, DMSO-$d_6$): $\delta$ two rotamers 172.85 (major), 172.48 (minor), 172.28 (minor), 172.24 (major), 153.51 (minor), 153.29 (major), 78.55 (minor), 78.41 (major), 59.30 (major), 59.05 (minor), 51.95, 50.75 (minor), 50.71 (major), 46.64 (minor), 46.49 (major), 30.92 (major), 30.75 (minor), 30.25 (major), 29.75 (minor), 29.64 (major), 29.45 (minor), 28.14 (minor), 27.98 (major), 23.89 (minor), 23.02 (major), 14.65 (minor), 14.43 (major). LRMS (ESI) $m/z$ 383 $[M + Na]^+$.

*tert*-Butyl (*S*)-2-(((10*S*,13*S*)-10-methyl-3,6,9,12-tetraoxo-2-oxa-16-thia-5,8,11-triazaheptadecan-13-yl)carbamoyl)pyrrolidine-1-carboxylate **(21)**.
Compound **20** (500 mg, 1.38 mmol) was converted to the target compound using general procedure B. The crude product was purified using flash column chromatography (30% to 50% THF/hexane) on silica gel to afford **21** (240 mg, 0.44 mmol, 32%) as a colorless oil.
$^{1}H$ NMR (400 MHz, DMSO-$d_6$): $\delta$ two rotamers 8.27 – 8.15 (m, 2H), 8.06 (d, $J$ = 6.9 Hz, major, 1H), 7.99 (brd, minor), 8.00 (d, $J$ = 7.6, 1H), 4.39 – 4.29 (m, 1H), 4.28 – 4.21 (m, 1H), 4.17 – 4.09 (m, 1H), 3.85 (dd, $J$ = 4.3, 5.73 Hz, 2H), 3.78 – 3.69 (m, 2H), 3.63 (s, 3H), 3.40 – 3.33 (m, 1H), 3.30 – 3.21 (m, 1H), 2.48 – 2.40 (m, 1H), 2.15 – 2.06 (m, 1H), 2.03 (s, 3H), 1.98 – 1.87 (m, 1H), 1.86 – 1.70 (m, 4H), 1.39 (s, minor), 1.32 (s, major, 9H), 1.22 (d, $J$ = 7.05 Hz, 3H); $^{13}C$ NMR (100 MHz, DMSO-$d_6$): $\delta$ two rotamers 172.57 (major), 172.32 (minor), 172.46, 170.97 (major), 170.90 (minor), 170.94 (minor), 170.21 (major), 169.29, 153.74 (minor), 153.31 (major), 78.72 (minor), 78.40 (major), 59.26 (minor), 59.23 (major), 51.76, 51.71 (minor), 51.61 (major), 48.41(minor), 48.34 (major), 46.72 (minor), 46.52 (major), 41.73, 40.54, 31.87 (major), 31.01 (minor), 29.74, 29.58 (major), 29.46 (minor), 28.16 (minor), 28.02 (major), 23.99 (minor), 23.06 (major), 18.02 (major), 17.90 (minor), 14.68 (minor), 14.54 (major). LRMS (ESI) $m/z$ 546 $[M + H]^+$.

*tert*-Butyl (*S*)-2-((*S*)-2-((*S*)-2-(((10*S*,13*S*)-10-methyl-3,6,9,12-tetraoxo-2-oxa-16-thia-5,8,11-triazaheptadecan-13-yl)carbamoyl)pyrrolidine-1-carbonyl)pyrrolidine-1-carbonyl)pyrrolidine-1-carboxylate **(22)**.
Compound **17** (3.5 g, 6.31 mmol) was converted to the target compound using general procedure B. The crude product was purified using flash column chromatography (0% to 10% $MeOH/CHCl_3$) on silica gel to afford **22** (2.01 g, 2.84, 45%) as a white solid.
$^{1}H$ NMR (400 MHz, DMSO-$d_6$) $\delta$ two rotamers 8.19 (dd, $J$ = 5.4, 6.7 Hz, 2H), 8.02 – 7.95 (m, 2H), 4.58 (dd, $J$ = 4.0, 8.7 Hz, minor), 4.55 (dd, $J$ = 3.7, 8.5 Hz, major, 1H) 4.43 – 4.18 (m, 4H), 3.86 (dd, $J$ = 3.2, 5.8 Hz, 2H), 3.73 (dd, $J$ = 6.0, 8.4 Hz, 2H), 3.63 (s, 3H), 3.66 – 3.40 (m, 4H), 3.31 – 3.26 (m, 2H), 2.48 – 2.37 (m, 2H), 2.20 – 1.98 (m, 3H), 2.03 (s, 3H), 1.98 – 1.68 (m, 11H), 1.37 (s, minor), 1.31 (s, major, 9H), 1.22 (d, $J$ = 7.2, 3H); $^{13}C$ NMR (100 MHz, DMSO-$d_6$) $\delta$ two rotamers 172.45, 171.89 (minor), 171.87 (major), 170.91, 170.51 (major), 170.09 (minor), 170.20, 169.93 (major), 169.91 (minor), 169.31, 153.34 (minor), 153.06 (major), 78.37 (minor), 78.28 (major), 59.33 (major), 59.27 (minor), 57.54, 57.38 (minor), 57.31 (major), 51.88, 51.76, 48.39, 46.67 (major), 46.62 (minor), 46.57 (minor), 46.50 (major), 46.41, 41.76, 40.54, 31.86, 29.45, 29.28 (major), 28.37 (minor), 28.82 (minor), 28.22 (major), 28.01(major), 27.76 (minor), 27.57 (major), 24.62, 24.40 (major), 24.36 (minor), 23.66 (minor), 23.15 (major), 17.87, 14.65.

LRMS (ESI) *m/z* 740 $[M + H]^+$. HRMS (ESI) m/z calculated for $C_{57}H_{77}N_{13}O_{14}S_2^+$ $[M + H]^+$: 740.3575. Found: 740.3612.

*tert*-Butyl (*S*)-2-((*S*)-2-((*S*)-2-(((10*S*,13*R*)-10-methyl-3,6,9,12-tetraoxo-2-oxa-16-thia-5,8,11-triazaheptadecan-13-yl)carbamoyl)pyrrolidine-1-carbonyl)pyrrolidine-1-carbonyl)pyrrolidine-1-carboxylate **(23)**.
Compound **18** (1.47 g, 2.65 mmol) was converted to the target compound using general procedure B. The crude product was purified using flash column chromatography (0% to 10% MeOH/$CHCl_3$) on silica gel to afford **23** (750 mg, 1.01 mmol, 38%) as a white solid.
$^1$H NMR (400 MHz, DMSO-*d$_6$*) δ two rotamers 8.31 (d, *J* = 8.3 Hz, major, 1H), 8.30 (d, *J* = 8.0 Hz, minor, 1H), 8.21 – 8.10 (m, 2H), 7.80 (d, *J* = 6.6 Hz, major, 1H), 7.77 (d, *J* = 6.8 Hz, minor, 1H), 4.59 (dd, *J* = 4.5, 8.5 Hz, minor), 4.55 (dd, *J* = 4.3, 8.4 Hz, major, 1H) 4.44 – 4.35 (m, 1H), 4.31 – 4.17 (m 3H), 3.90 – 3.80 (m, 2H), 3.77 – 3.70 (m, 2H), 3.63 (s, 3H), 3.69 – 3.37 (m, 4H), 3.45 – 3.37 (m, 1H) 3.31 – 3.25 (m, 2H), 2.48 – 2.34 (m, 2H), 2.21 – 1.97 (m, 4H), 2.01 (s, 3H), 1.95 – 1.65 (m, 10H), 1.37 (s, minor), 1.31 (s, major, 9H), 1.23 (d, *J* = 7.1, 3H); $^{13}$C NMR (101 MHz, DMSO-*d$_6$*) δ two rotamers 172.83, 172.60 (minor), 172.56 (major), 171.58 (major), 171.55 (minor), 170.92 (major), 170.80 (minor), 170.63, 170.61 (major), 170.31 (minor), 169.79, 153.76 (minor), 153.47 (major), 78.79 (minor), 78.71 (major), 60.36 (major), 60.27 (minor), 57.96, 57.74 (minor), 57.67 (major), 52.18, 52.05, 49.13 (major), 49.09 (minor), 47.33, 46.98 (major), 46.96 (minor), 46.89 (minor), 46.83 (major), 42.18, 40.94, 31.46, 30.01, 29.67 (major), 28.77 (minor), 29.36 (minor), 29.33 (major), 28.64 (minor), 28.43 (major), 28.14 (major), 27.97 (minor), 25.32, 24.99 (major), 24.97 (minor), 24.12 (minor), 23.60 (major), 18.01 (minor), 17.98 (major), 15.05. LRMS (ESI) *m/z* 740 $[M + H]^+$.

*tert*-Butyl (*S*)-2-((*S*)-2-((*S*)-2-(((10*S*,13*S*)-10-methyl-3,6,9,12-tetraoxo-2,16-dioxa-5,8,11-triazaheptadecan-13-yl)carbamoyl)pyrrolidine-1-carbonyl)pyrrolidine-1-carbonyl)pyrrolidine-1-carboxylate **(24)**.
Compound **19** (1.0 g, 1.86 mmol) was converted to the target compound using general procedure B. The crude product was purified using flash column chromatography (0% to 10% MeOH/$CHCl_3$) on silica gel to afford **24** (405 mg, 0.56 mmol, 30%) as a white solid.
$^1$H NMR (400 MHz, DMSO-*d$_6$*) δ two rotamers 8.17 (t, *J* = 5.9 Hz, 1H), 8.12 (t, *J* = 5.9 Hz, 1H), 7.95 (dd, *J* = 3.7, 7.6 Hz), 7.90 (d, *J* = 7.0 Hz), 4.59 (dd, *J* = 4.2, 8.8 Hz, minor), 4.55 (dd, *J* = 3.7, 8.5 Hz, major, 1H), 4.45 – 4.35 (m, 1H), 4.34 – 4.28 (m, 1H), 4.27 – 4.16 (m, 2H), 3.85 (dd, *J* = 2.8, 5.8 Hz, 2H), 3.73 (dd, *J* = 6.0, 8.4 Hz, 2H), 3.65 – 3.41 (m, 4H), 3.62 (s, 3H), 3.36 – 3.27 (m, 4H), 3.18 (s, 3H), 2.18 – 1.98 (m, 3H), 1.97 – 1.67 (m, 11H), 1.37 (s, minor), 1.31 (s, major, 9H), 1.21 (d, *J* = 7.1, 3H); $^{13}$C NMR (101 MHz, DMSO-*d$_6$*) δ two rotamers 172.41, 171.86 (minor), 171.84 (major), 171.22, 170.54 (major), 170.20 (minor), 170.21, 170.03 (major), 169.94 (minor), 169.32, 153.36 (minor), 153.07 (major), 93.89, 78.38 (minor), 78.29 (major), 68.51, 59.42 (major), 59.35 (minor), 57.91, 57.60, 57.38 (minor), 57.32 (major), 51.76, 50.04, 48.38, 46.69 (major), 46.65 (minor), 46.58 (minor), 46.51 (major), 46.42, 41.78, 40.54, 31.60, 29.30 (major), 28.38 (minor), 28.85, 28.22 (minor), 28.01 (major), 27.77 (major), 27.59 (minor), 24.60, 24.44 (major), 24.41 minor), 23.68 (minor), 23.16 (major), 17.87. LRMS (ESI) *m/z* 724 $[M + H]^+$.

**General Procedure C for the Synthesis of KLHDC2 PROTACs.**

To a solution of corresponding peptide intermediates (1.0 equiv) in DCM (0.5 M) was added HCl (4.0 M in 1,4-dioxane, 10 equiv) at 0 °C. After stirring at room temperature for 3 h, the solution was concentrated. To the crude peptide in DMF (0.1 M) was added **6** (0.8 equiv), TEA (4.0 equiv) followed by 1-propanephosphonic anhydride (50% w/w in EA, 1.6 equiv) at – 10 °C. The reaction mixture was then stirred at -10 °C for 30 min, quenched with MeOH and concentrated under reduced pressure. To the solution of crude ester in THF (0.1 M) was added aqueous LiOH solution (5.0 equiv) at -10 °C. The reaction mixture was then stirred at -10 °C for 30 min and diluted with DMF. The resulting mixture was subjected to ACCQPREP HP150 system (0% to 50% ACN/$H_2O$ (0.1% formic acid) to afford compound **25-28** as yellow solid (33-57%).

(3-(2-(2-(4-(4-((7-(3-(methylsulfonamido)phenyl)thieno[3,2-d]pyrimidin-2-yl)amino)phenyl)piperazin-1-yl)ethoxy)ethoxy)propanoyl)-*L*-prolyl-*L*-methionyl-*L*-alanylglycylglycine **(25)**.
Compound **21** (50 mg, 0.07 mmol) was converted to the target compound using general procedure C. The product was purified with ACCQPREP HP150 system using XBridge BEH Shield RP18 column (19 × 250 mm, 10 $\mu$m particle size; 0% to 50% ACN/$H_2O$ (0.1% formic acid) to afford compound **25** as a yellow solid (41.0 mg, 0.03 mmol, 43%).
$^1$H NMR (400 MHz, DMSO-$d_6$): $\delta$ two rotamers 9.84 (s, 1H), 9.44 (s, 1H), 9.16 (s, 1H), 8.46 (s, 1H), 8.29 (d, $J$ = 8.2 Hz, minor), 8.19 (t, $J$ = 5.8 Hz, minor), 8.13 – 7.99 (m, 3H), 7.86 (d, $J$ = 7.0 Hz, 1H), 7.83 (s, 1H), 7.80 (d, $J$ = 7.8 Hz, 1H), 7.69 (d, $J$ = 9.0 Hz, 2H), 7.48 (t, $J$ = 7.9 Hz, 1H), 7.27 (dd, $J$ = 1.3, 8.1 Hz, 1H), 6.91 (d, $J$ = 9.1 Hz, 2H), 4.45 (dd, $J$ = 2.7, 8.5, minor), 4.41 – 4.34 (m, minor), 4.32 – 4.27 (m, 2H), 4.26 – 4.19 (m, 1H), 3.77 – 3.70 (m, 4H), 3.67 – 3.53 (m, 6H), 3.50 – 3.35 (m, 6H), 3.07 (brt, 4H), 3.02 (s, 3H), 2.64 – 2.52 (m, 6H), 2.48 – 2.37 (m, 2H), 2.23 – 2.11 (m, 1H), 2.04 (s, minor), 2.03 (s, major, 3H), 2.00 – 1.71 (m, 5H), 1.24 (d, $J$ = 7.1 Hz, major, 3H), 1.22 (d, $J$ = 7.0 Hz, minor); $^{13}$C NMR (100 MHz, DMSO-$d_6$): $\delta$ two rotamers 172.45 (minor), 172.01 (major), 172.40, 172.05 (minor), 171.18 (major), 170.97 (major), 170.95 (minor), 169.43 (major), 169.41 (minor), 167.00, 158.41, 157.95, 154.01, 146.05, 138.56, 134.60, 134.40, 133.69, 132.77, 129.35, 123.82, 122.20, 120.08, 120.00, 119.13, 115.93, 69.67, 68.21, 66.48 (minor), 66.32 (major), 59.51 (minor), 59.44 (major), 57.15, 53.16, 51.79 (major), 51.74 (minor), 48.93, 48.44 (major), 48.36 (minor), 47.12 (minor), 47.02 (major), 41.85 (major), 41.79 (minor), 40.77, 34.46, 31.45, 29.67 (minor), 29.33 (major), 29.63, 24.34, 18.06 (minor), 17.87 (major), 14.68. LRMS (ESI) *m/z* 528 [M/2 + H]$^+$; 1054 [M + H]$^+$. HRMS (ESI) m/z calculated for 1/2 $C_{47}H_{63}N_{11}O_{11}S_3^+$ [M/2 + H]$^+$: 527.6936. Found: 527.6990; $C_{47}H_{63}N_{11}O_{11}S_3^+$ [M + H]$^+$: 1054.3871. Found: 1054.3889.

(3-(2-(2-(4-(4-((7-(3-(methylsulfonamido)phenyl)thieno[3,2-d]pyrimidin-2-yl)amino)phenyl)piperazin-1-yl)ethoxy)ethoxy)propanoyl)-*L*-prolyl-*L*-prolyl-*L*-prolyl-*L*-methionyl-*L*-alanylglycylglycine **(26)**.
Compound **22** (50 mg, 0.07 mmol) was converted to the target compound using general procedure C. The product was purified with ACCQPREP HP150 system using XBridge BEH Shield RP18 column (19 × 250 mm, 10 $\mu$m particle size; 0% to 50% ACN/$H_2O$ (0.1% formic acid) to afford compound **26** as a yellow solid (31.5 mg, 0.03 mmol, 43%).
$^1$H NMR (400 MHz, DMSO-$d_6$): $\delta$ two rotamers 9.84 (brs, 1H), 9.44 (s, 1H), 9.16 (s, 1H), 8.46 (s, 1H), 8.16 – 8.13 (m, 1H), 8.06 – 8.00 (m, 1H), 7.99 – 7.93 (m, 2H), 7.84 (s, 1H), 7.80 (d, $J$ = 7.9 Hz 1H), 7.69 (d, $J$ = 9.0 Hz, 2H), 7.48 (t, $J$ = 7.9 Hz, 1H), 7.27 (dd, $J$ = 1.3, 8.1 Hz, 1H), 6.91

(d, $J$ = 9.0 Hz , 2H), 4.78 (dd, $J$ = 2.7, 8.5 Hz, minor), 4.62 (dd, $J$ = 4.5, 8.5 Hz, minor), 4.57 – 4.49 (m, 2H, major), 4.34 – 4.29 (m, 1H), 4.29 – 4.19 (m, 2H), 3.78 – 3.69 (m, 4H), 3.68 – 3.51 (m, 10 H), 3.47 – 3.33 (m, 6H), 3.05 (brt, 4H), 3.02 (s, 3H), 2.60 – 2.56 (brt, 4H), 2.55 – 2.52 (m, 2H), 2.48 – 2.38 (m, 2H), 2.24 – 2.01 (m, 3H), 2.02 (s, 3H), 1.97 – 1.71 (m, 11H), 1.22 (d, $J$ = 7.1 Hz, 3H); $^{13}$C NMR (100 MHz, DMSO-$d_6$): $\delta$ two rotamers 172.44, 171.89 (minor), 171.82 (major), 171.87 (minor), 171.17 (major), 170.87, 170.12 (minor), 169.62 (major), 169.66 (minor), 169.55 (major), 169.00, 168.24, 158.41, 157.95, 154.03, 146.10, 138.57, 134.60, 134.42, 133.69, 132.76, 129.37, 123.83, 122.21, 120.07, 120.02, 119.14, 115.92. 69.70, 69.66, 68.33 (major), 68.31 (minor), 66.49 (minor), 66.30 (major), 59.25 (major), 59.23 (minor), 57.49, 57.23, 57.13, 53.22, 51.88, 49.01, 48.36, 46.86, 46.65 (minor), 46.63 (major), 46.53 (major), 46.48 (minor), 41.81, 40.79, 34.34, 31.88, 29.45, 28.84, 28.79 (minor), 27.91 (major), 27.69 (major), 27.63 (minor), 24.60, 24.58, 24.38 (major), 24.14 (minor), 17.93, 14.66. LRMS (ESI) *m/z* 625 [M/2 + H]$^+$; 1248 [M + H]$^+$. HRMS (ESI) m/z calculated for 1/2 $C_{57}H_{77}N_{13}O_{13}S_3^+$ [M/2 + H]$^+$: 624.7463. Found: 624.7516; $C_{57}H_{77}N_{13}O_{13}S_3^+$ [M + H]$^+$: 1248.4926. Found: 1248.4956.

(3-(2-(2-(4-(4-((7-(3-(methylsulfonamido)phenyl)thieno[3,2-d]pyrimidin-2-yl)amino)phenyl)piperazin-1-yl)ethoxy)ethoxy)propanoyl)-*L*-prolyl-*L*-prolyl-*L*-prolyl-*D*-methionyl-*L*-alanylglycylglycine **(27)**.

Compound **23** (50 mg, 0.07 mmol) was converted to the target compound using general procedure C. The product was purified with ACCQPREP HP150 system using XBridge BEH Shield RP18 column (19 × 250 mm, 10 $\mu$m particle size; 0% to 50% ACN/$H_2O$ (0.1% formic acid) to afford compound **27** as a yellow solid (42.8 mg, 0.04 mmol, 57%).

$^1$H NMR (400 MHz, DMSO-$d_6$): $\delta$ two rotamers 9.85 (brs, 1H), 9.45 (s, 1H), 9.17 (s, 1H), 8.47 (s, 1H), 8.35 – 8.27 (m, 1H), 8.12 (t, $J$ = 5.9 Hz, 1H), 8.03 – 7.95 (m, 1H), 7.84 (s, 1H), 7.83 – 7.75 (m, 2H), 7.70 (d, $J$ = 9.0 Hz, 2H), 7.49 (t, $J$ = 7.9 Hz, 1H), 7.27 (dd, $J$ = 1.3, 8.1 Hz, 1H), 6.92 (d, $J$ = 9.07 Hz, 2H), 4.79 (dd, $J$ = 2.8, 8.6 Hz, minor), 4.63 (dd, $J$ = 5.0, 8.4 Hz, minor), 4.58 – 4.50 (m, 2H), 4.33 – 4.17 (m, 3H), 3.77 – 3.71 (m, 4H), 3.69 – 3.53 (m, 9H), 3.50 – 3.36 (m, 7H), 3.07 (brt, 4H), 3.03 (s, 3H), 2.63 – 2.53 (m, 6H), 2.48 – 2.35 (m, 2H), 2.24 – 1.96 (m, 5H), 2.02 (s, 3H), 1.94 – 1.68 (m, 9H), 1.24 (d, $J$ = 7.1 Hz, 3H); $^{13}$C NMR (100 MHz, DMSO-$d_6$): $\delta$ two rotamers 172.36, 172.31 (minor), 172.11 (major), 172.13 (major), 171.19 (minor), 171.07 (major), 171.03 (minor), 170.40 (minor), 169.61 (major), 169.65 (minor), 169.51 (major), 169.03, 168.22, 158.40, 157.94, 154.02, 146.07, 138.56, 134.60, 134.41, 133.68, 132.76, 129.36, 123.82, 122.20, 120.67, 120.01, 119.13, 115.92, 69.70, 69.66, 68.29, 66.49 (minor), 66.30 (major), 59.88 (major), 59.81 (minor), 57.51, 57.20, 57.08, 53.19, 51.60, 48.97, 48.65 (major), 48.56 (minor), 46.87, 46.86, 46.55, 41.82, 40.77, 34.34, 31.03, 29.59, 28.92, 28.89 (minor), 28.05 (major), 27.65 (major), 27.52 (minor), 24.88, 24.84, 24.54 (minor), 24.18 (major), 17.89 (minor), 17.62 (major), 14.64. LRMS (ESI) *m/z* 625 [M/2 + H]$^+$; 1248 [M + H]$^+$. HRMS (ESI) m/z calculated for 1/2 $C_{57}H_{77}N_{13}O_{13}S_3^+$ [M/2 + H]$^+$: 624.7463. Found: 624.7515; $C_{57}H_{77}N_{13}O_{13}S_3^+$ [M + H]$^+$: 1248.4926. Found: 1248.4953.

*O*-methyl-*N*-(3-(2-(2-(4-(4-((7-(3-(methylsulfonamido)phenyl)thieno[3,2-d]pyrimidin-2-yl)amino)phenyl)piperazin-1-yl)ethoxy)ethoxy)propanoyl)-*L*-prolyl-*L*-prolyl-*L*-prolyl-*L*-homoseryl-*L*-alanylglycylglycine **(28)**.

Compound **24** (50 mg, 0.09 mmol) was converted to the target compound using general procedure C. The product was purified with ACCQPREP HP150 system using XBridge BEH Shield RP18 column (19 × 250 mm, 10 $\mu$m particle size; 0% to 50% ACN/$H_2O$ (0.1% formic acid) to afford compound **28** as a yellow solid (28.3 mg, 0.03 mmol, 33%).
$^1$H NMR (400 MHz, DMSO-$d_6$): $\delta$ two rotamers 9.84 (s, 1H), 9.44 (s, 1H), 9.16 (s, 1H), 8.46 (s, 1H), 814 – 8.08 (m, 1H), 8.03 (m, 1H), 7.97 – 7.88 (m, 2H), 7.84 (s, 1H), 7.80 (d, *J* = 7.81 Hz, 1H), 7.70 (d, *J* = 7.8 Hz, 2H), 7.48 (t, *J* = 7.9 Hz, 1H), 7.27 (dd, *J* = 1.3, 8.1 Hz, 1H), 6.91 (d, *J* = 1.3, 8.1 Hz, 2H), 4.78 (dd, *J* = 8.7, 2.6 Hz, minor), 4.62 (dd, *J* = 4.5, 8.6 Hz, minor), 4.58 – 4.45 (m, 2H), 4.36 – 4.26 (m, 1H), 4.26 – 4.16 (m, 2H), 3.77 – 3.69 (m, 4H), 3.68 – 3.52 (m, 7H), 3.49 – 3.27 (m, 11H), 3.19 (s, 3H), 3.06 (brt, 4H), 3.02 (s, 3H), 2.62 – 2.52 (m, 6H), 2.24 – 1.96 (m, 3H), 1.97 – 1.64 (m, 11H), 1.21 (d, *J* = 7.1 Hz, 3H) $^{13}$C NMR (100 MHz, DMSO-$d_6$): $\delta$ two rotamers 172.37, 171.80 (minor), 171.16 (major), 171.74 (major), 171.63 (minor), 171.15, 170.20 (minor), 169.62 (major), 169.90 (major), 169.55 (minor), 169.00, 168.23, 158.40, 157.94, 154.01, 146.08, 138.56, 134.60, 134.40, 133.68, 132.76, 129.36, 123.81, 122.20, 120.06, 120.01, 119.13, 115.91, 69.70, 69.66, 68.49, 68.31, 66.48 (minor), 66.29 (major), 59.29 (major), 59.28 (minor), 57.90, 57.49, 57.22, 57.12, 53.21, 50.01, 48.99, 48.34, 46.68, 46.64 (major), 46.63 (minor), 46.52 (major), 46.32 (minor), 41.82, 40.74, 34.34, 31.62, 28.84, 28.76 (minor), 27.91 (major), 27.69 (major), 27.52 (minor), 24.56, 24.53 (major), 24.51 (minor), 24.39 (minor), 24.14 (major), 17.91. LRMS (ESI) *m/z* 617 $[M/2 + H]^+$; 1232 $[M + H]^+$. HRMS (ESI) m/z calculated for 1/2 $C_{57}H_{77}N_{13}O_{14}S_2^+$ $[M/2 + H]^+$: 616.7577. Found: 616.7630; $C_{57}H_{77}N_{13}O_{14}S_2^+$ $[M + H]^+$: 1232.5154. Found: 1232.5182.

*Cell Culture and Proliferation Assays*
Cell lines were cultured in RPMI-1640 (MOLT-4, MOLM-14, DLD1), Leibovitz's L-15 medium supplemented with 10% FBS (MDA-MB-231 and SW480), Eagle's Minimum Essential Medium (MCF7), Iscove's Modified Dulbecco's Medium (MV-4-11), or Dulbecco's Modified Essential Medium (GIST-T1). All growth media were supplemented with 10% fetal bovine serum (FBS) and 1% penicillin-streptomycin. Cells were tested periodically for mycoplasma contamination.

For cell proliferation assays, MOLM-14 cells were plated at low density in 384-well white plates and treated in triplicate or quadruplicate with the indicated compounds. DMSO concentration was balanced for all conditions. Compounds were dispensed from 10 mM stock solutions using a D300 drug printing robot (Tecan). 72 hours after application of compounds, Cell Titer Glo reagent (Promega) was applied and incubated for 10-15 min at room temperature before reading luminescence on a PHERAStar plate reader (BMG LABTECH). For all treatments, luminescence was normalized to DMSO-only wells or DMSO-MLN4924 at the matching MLN4924 concentration.

*HiBiT Assays*
The indicated kinases were fused with the HiBiT sequence (VSGWRLFKKIS) linked to the protein of interest via a GSGS linker. Codon-optimized coding sequences were ordered as gBlock fragments (IDT) and cloned into vector pN103 under the control of the EF1a promoter. Corresponding lentivirus was used to transduce MOLM-14 cells cultured as described above, and tranductants were selected by growth in 1 µg/ml puromycin added directly to the culture medium. Pooled selectants were verified for kinase expression using the lytic HiBiT assay as described by the manufacturer (Promega). For HiBiT measurements, cells were dispensed into 384-well white plates and treated with compounds in triplicate or quadruplicate. Six hours after treatment, a lytic HiBiT assay was performed, and luminescence signal was measured as described above. For MLN4924 experiments, cells were pretreated for two hours with 1 µM MLN4924 before the six hour incubation with test compound. For these experiments, HiBiT signal was normalized to the Cell Titer Glo cell viability signal for each matching condition and to DMSO for each treatment (DMSO and MLN4924). We noticed that DB0625 and DB0614, which differ only in the length of their linkers (*2*), performed similarly or identically in this assay for the kinases tested. We show DB0625, for which we have the more complete dataset.

*Western Blotting*
Compound treatments and times are indicated in the figures. After treatment, cells were collected, washed once in PBS, and resuspended in cell lysis buffer supplemented with protease inhibitors. Equal amounts of protein from each lysate were prepared for Western blotting by boiling in SDS-PAGE sample buffer buffer. For blotting, membranes were blocked in TBST/5% milk (*w:v*) for at least one hour before probing with the following primary antibodies: CDK4, WEE1, GAPDH, NEK9, CDK6, FAK, Actin, KLHDC2, CRBN, Tubulin. Secondary antibodies (goat anti-mouse IgG or anti-rabbit IgG) were used at 1:10,000 concentration.

*Protein Purification*
Codon-optimized KLHDC2- or KLHDC2-K147A-1-362 (coding for the propellor domain) was amplified by PCR from a gBlock fragment (IDT) and cloned into an N-terminal His6-TEV-GST fusion vector by ligation-independent cloning. The cloned gene was verified by Sanger sequencing and transformed into Rosetta2(DE3)pLysS chemically competent *E. coli* cells

(Novagene) and grown in Luria Broth supplemented with chloramphenicol and carbenicillin overnight. Saturated overnight cultures were distributed to 1 L flasks containing 2XYT medium supplemented with antibiotics. Protein expression was induced by addition of 400 μM IPTG (final concentration). The temperature was adjusted from 37° to 18° Celsius. After incubation overnight, cells were harvested by centrifugation. Cell pellets were resuspended in ~4 ml/L D800 buffer (20 mM HEPES, pH 7.5; 800 mM NaCl; 10 mM imidazole, pH 8.0; 10 % glycerol, 2 mM beta mercaptoethanol) supplemented with protease inhibitors (1 mM PMSF, 1 mM benzamidine, ~20 ug/ml pepstatin, aprotinin, and leupeptin) and frozen at -80°C.

Cell pellets were thawed briefly in warm water and lysed by sonication and addition of solid lysozyme before centrifugation at 16,233 rcf for 1 hr at 12 °C. Clarified lysate was mixed with ~0.5 ml/L of growth cobalt resin (TaKaRa) for one hour before centrifugation at low speed to separate the beads, which were subsequently washed by gravity flow with ~25 column volumes ice cold D800 buffer before a final wash with B50 (D800 with 50 mM NaCl) and elution with C50 (B50 with 400 mM imidazole, pH 8.0). Cobalt eluate was applied to a 5 ml anion exchange column (Q HP, Cytiva) and eluted with an 8-column volume gradient from B50 to D800. Peak fractions were concentrated by ultrafiltration before application to a 24 ml gel filtration column (S200 increase, Cytiva) primed with GF150 buffer (20 mM Tris-HCl, pH 8.5, 150 mM NaCl, 1 mM TCEP). Peak fractions were again concentrated by ultrafiltration, supplemented with 5% glycerol (*v:v,* final), and aliquoted and frozen at -80° C.

*TR-FRET Binding Assay*

FITC-PPPMAGG probe (Elim) was first resuspended in DMSO before verification of the stock concentration by absorbance at 488 upon dilution in PBS. A stock solution of Tb-anti-GST antibody (CSBio) was made according to the manufacturer's recommendation in TR-FRET buffer (20 mM HEPES, pH 7.5, 150 mM NaCl, 1 mM TCEP, 0.1% NP-40 substitute, 0.1% BSA). Recombinant His6-GST-KLHDC2 and FITC-PPPMAGG probe were titrated systematically to identify TR-FRET conditions with i) minimal probe concentration, ii) a suitable assay window, and iii) a KLHDC2 concentration giving maximum responsiveness to changing protein concentration (~20% maximum signal). Final concentrations were as follows: 2 nM FITC-PPPMAGG, 3.2 nM His6-GST-KLHDC2, and 0.83 nM Tb-anti-GST.

For binding assays, a TR-FRET assay mix was prepared according to concentrations above and dispensed into a 384-well black assay plate. The indicated compounds were dispensed as above. After a one hour incubation at room temperature, the plate was read on a PHERAStar plate reader (BMG LABTECH, TRF 337 520 490 Optic Module).

**$^1$H and $^{13}$C NMR spectra of intermediates**

*tert*-butyl 3-(2-(2-hydroxyethoxy)ethoxy)propanoate **(2)**.

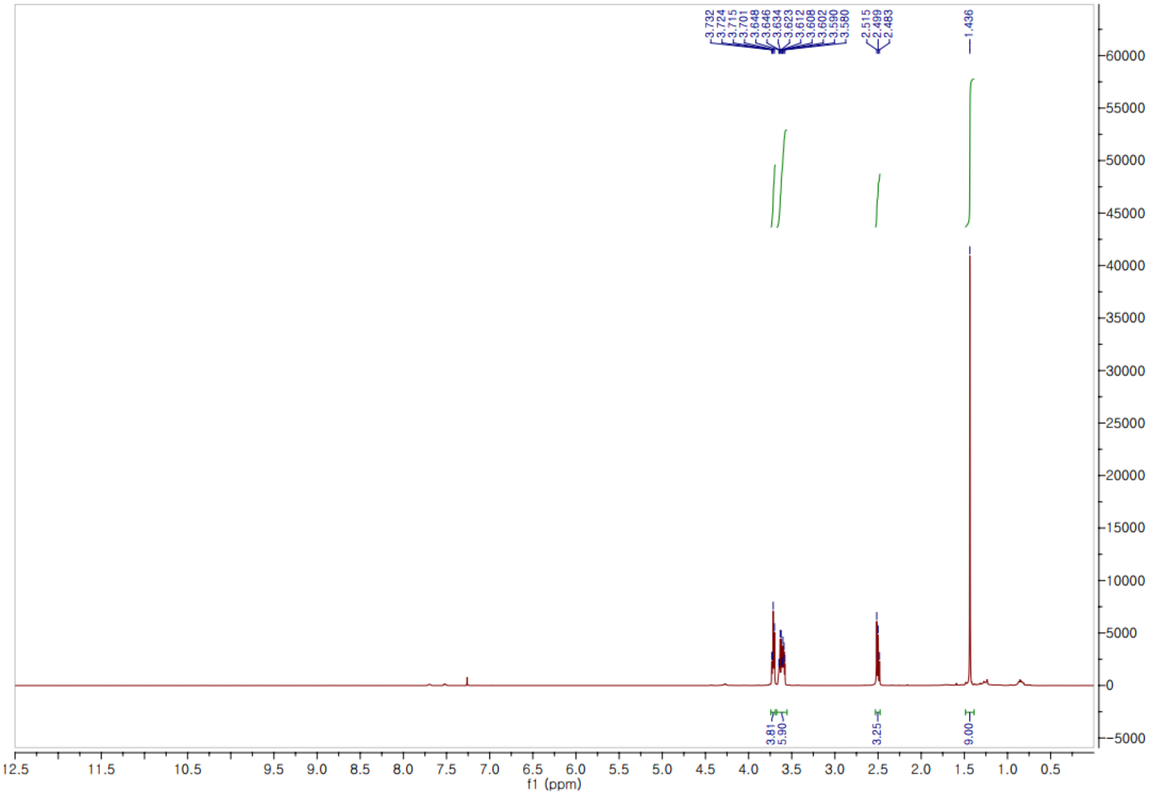


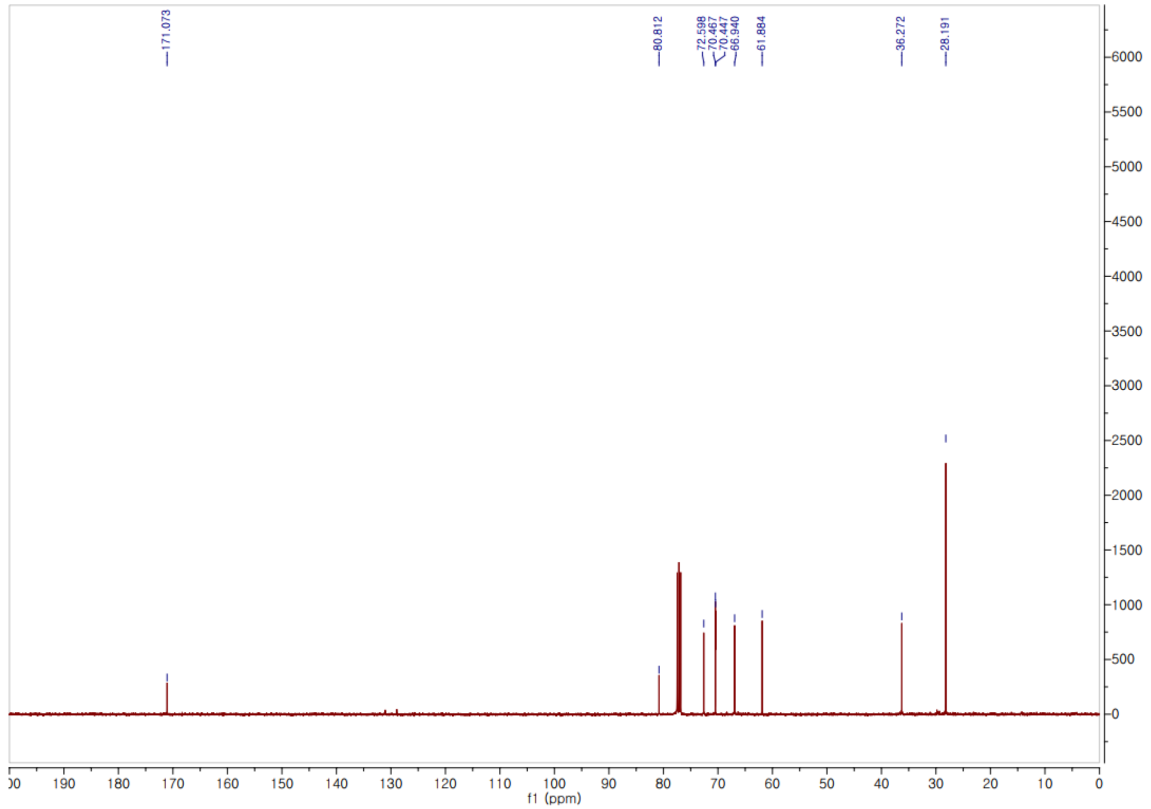

*tert*-butyl 3-(2-(2-iodoethoxy)ethoxy)propanoate **(3)**.

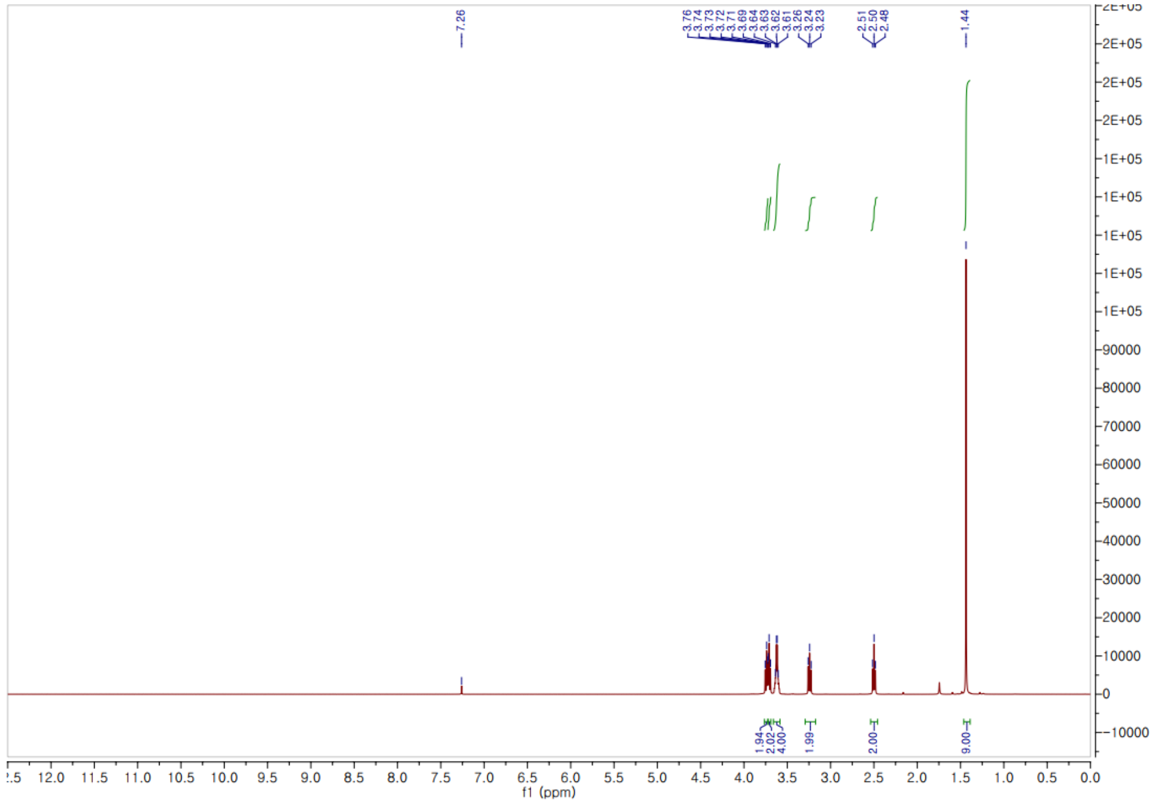

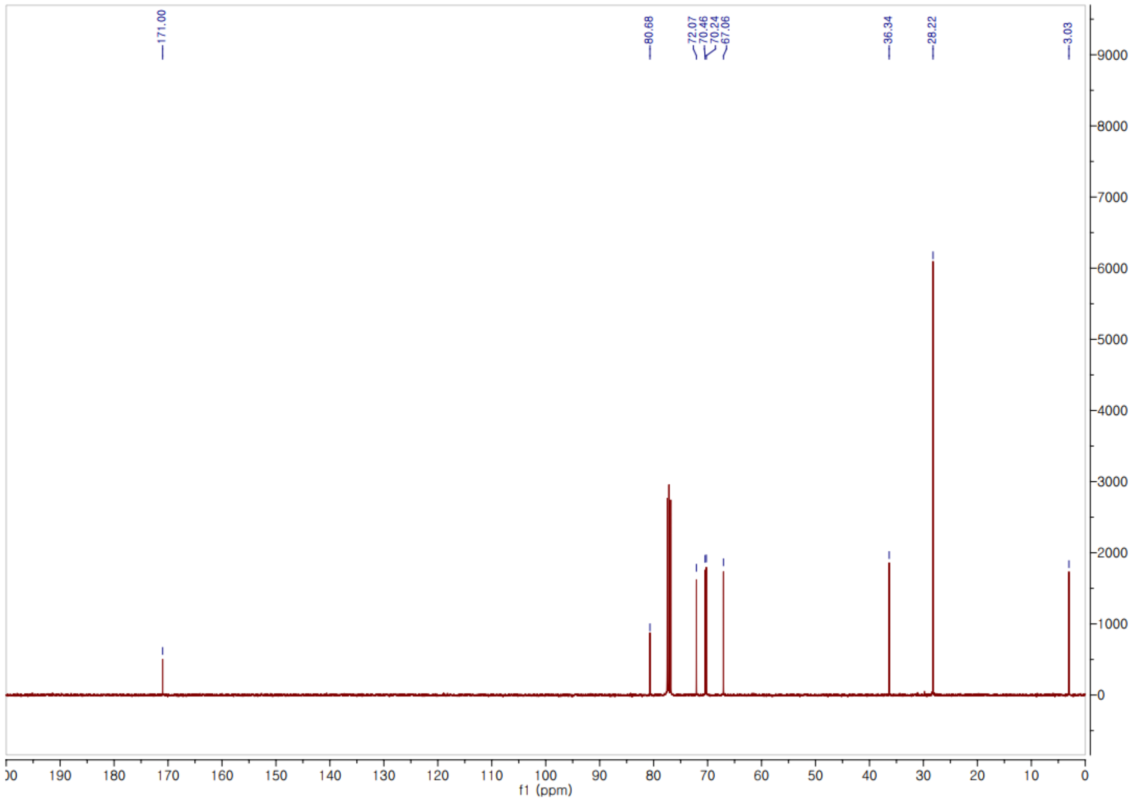

*tert*-butyl 3-(2-(2-(4-(4-((7-(3-(methylsulfonamido)phenyl)thieno[3,2-d]pyrimidin-2-yl)amino)phenyl)piperazin-1-yl)ethoxy)ethoxy)propanoate **(5)**.

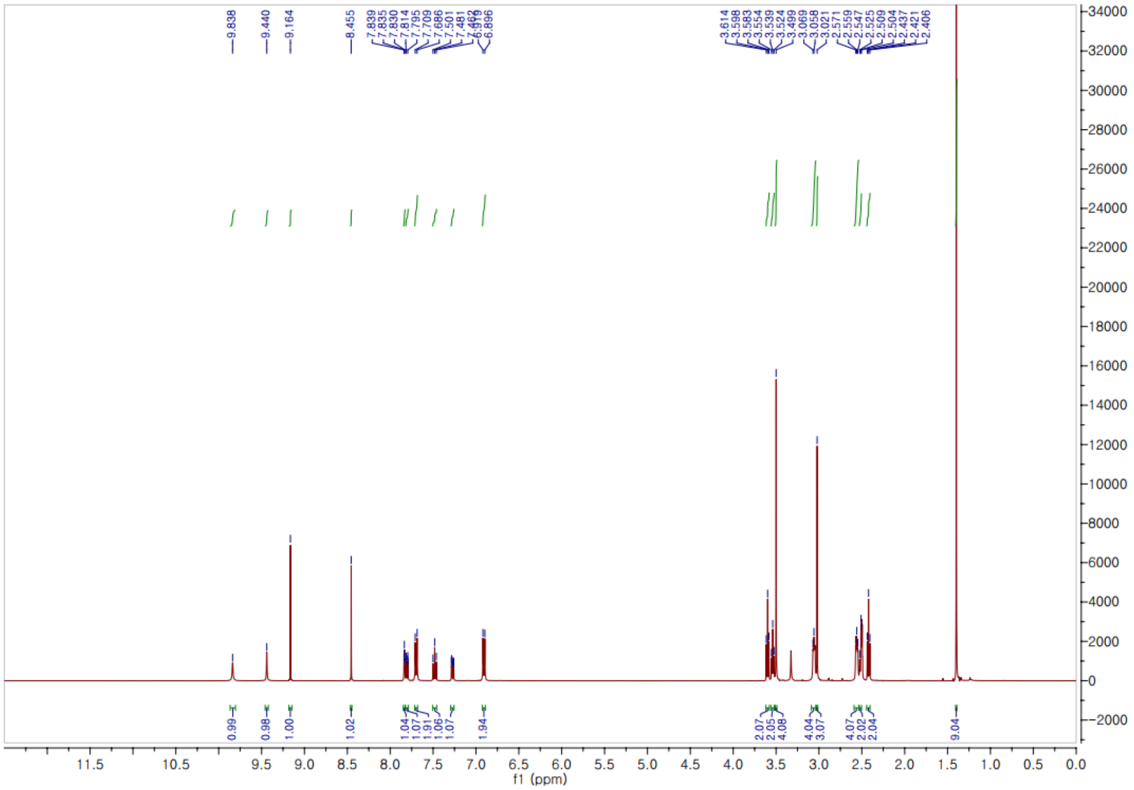


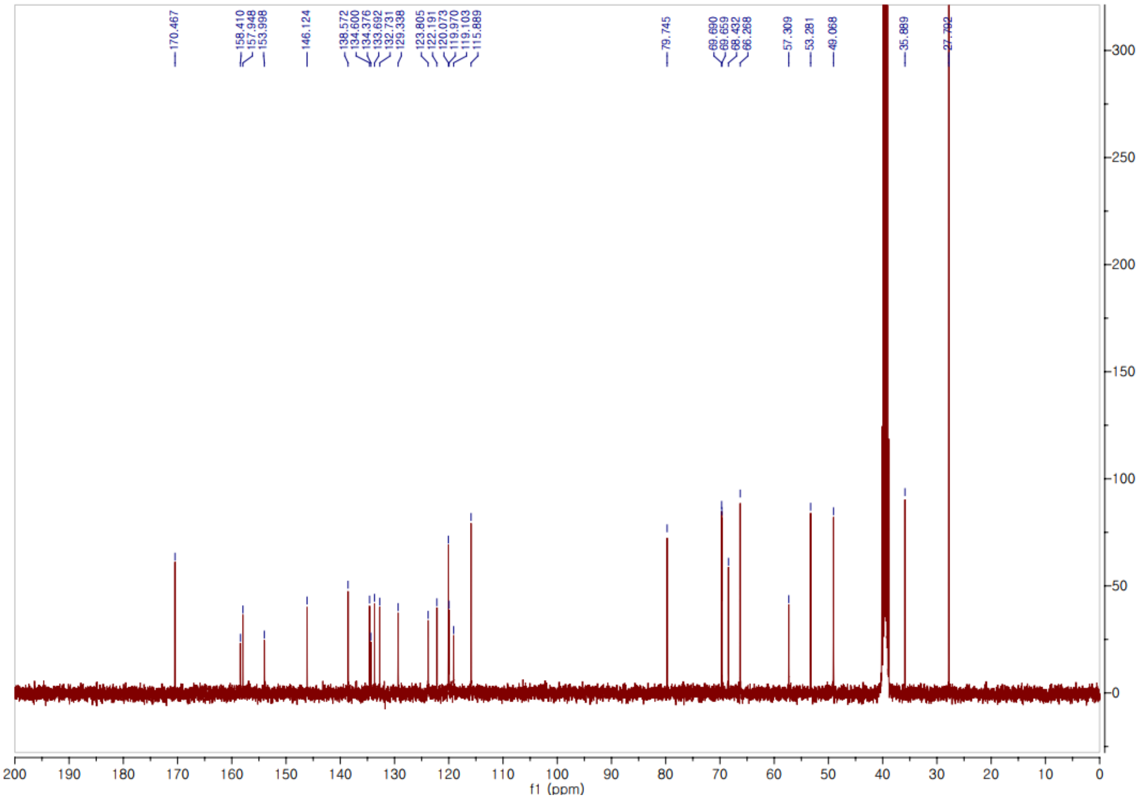

Methyl (*tert*-butoxycarbonyl)-*L*-alanylglycinate **(8)**.

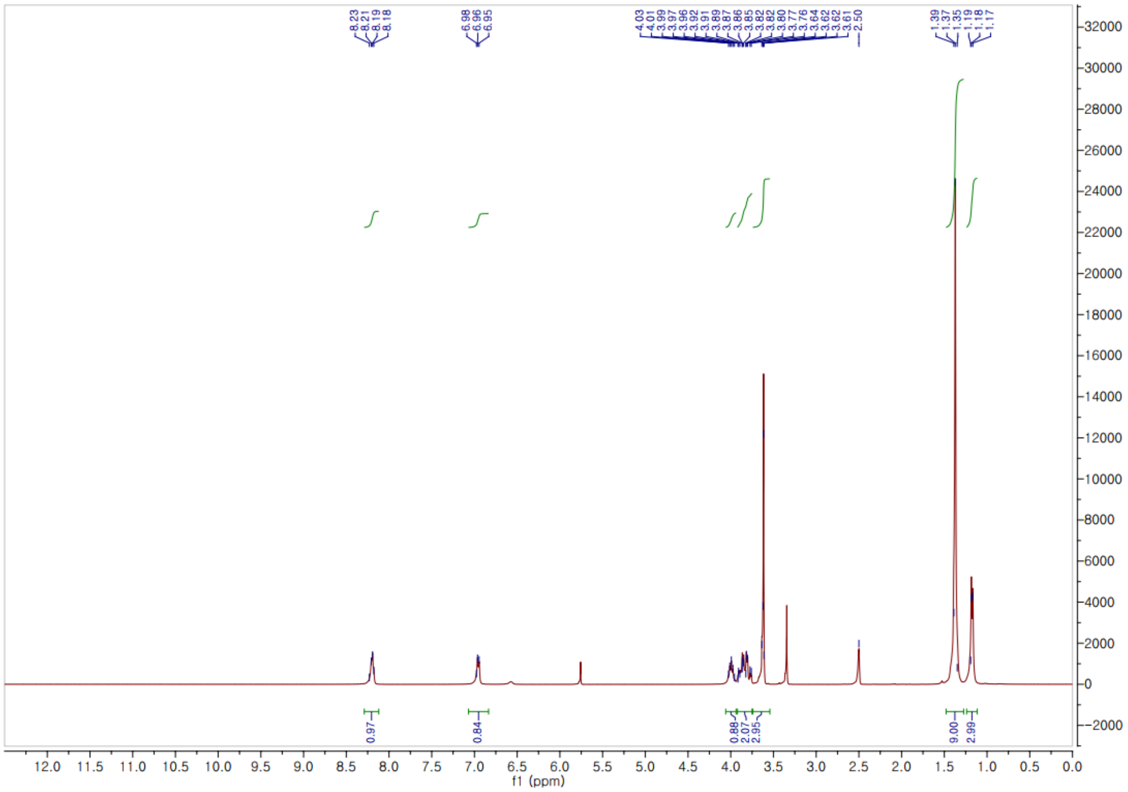


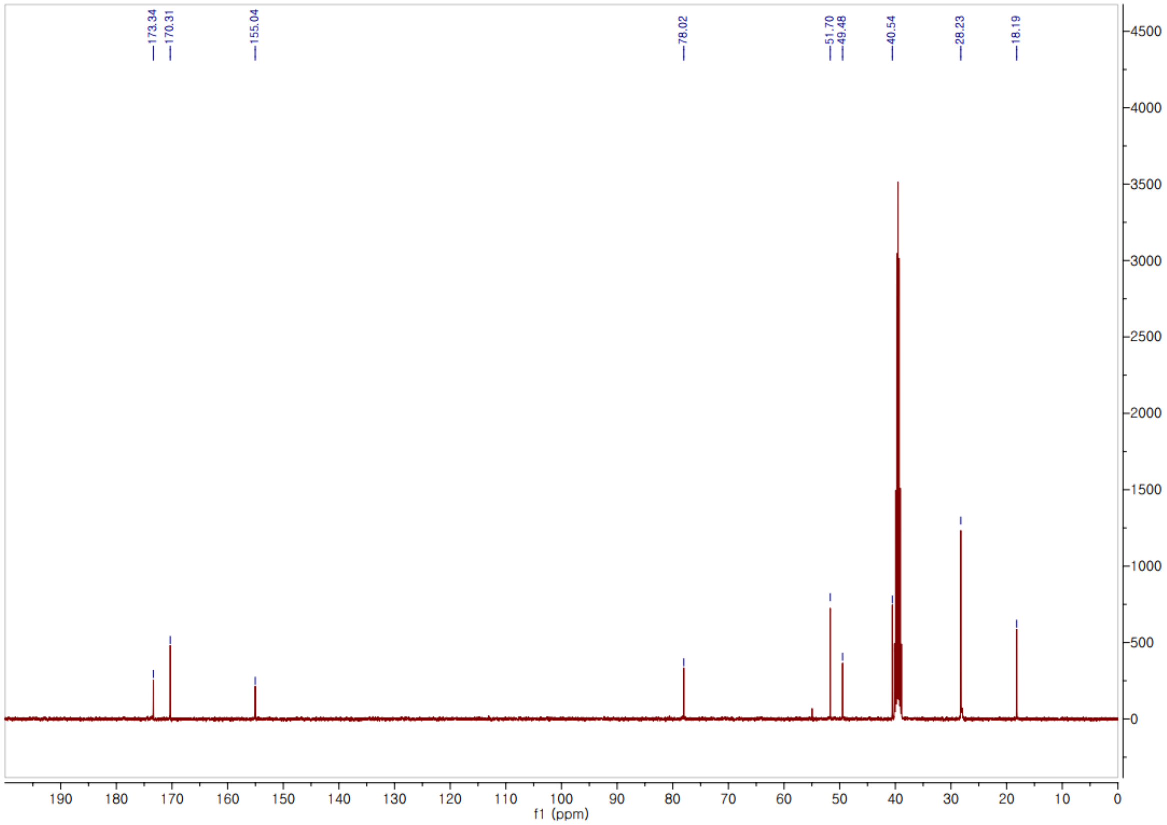

Methyl (*tert*-butoxycarbonyl)-*L*-alanylglycylglycinate **(9)**.

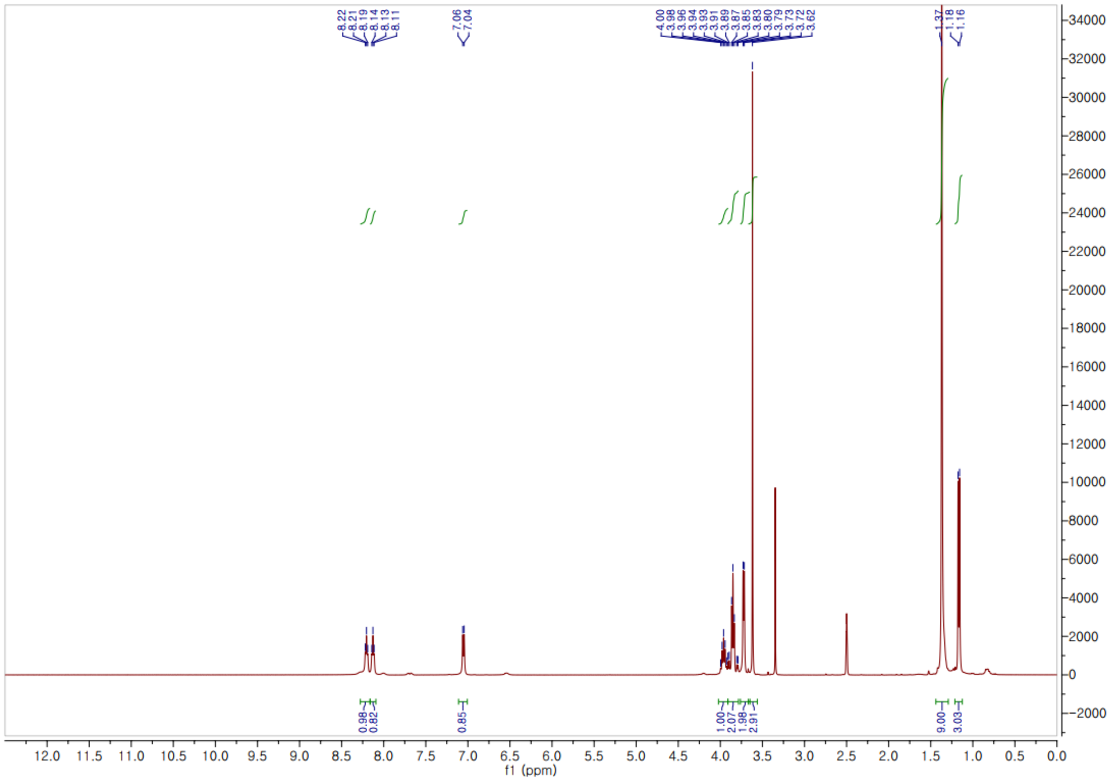


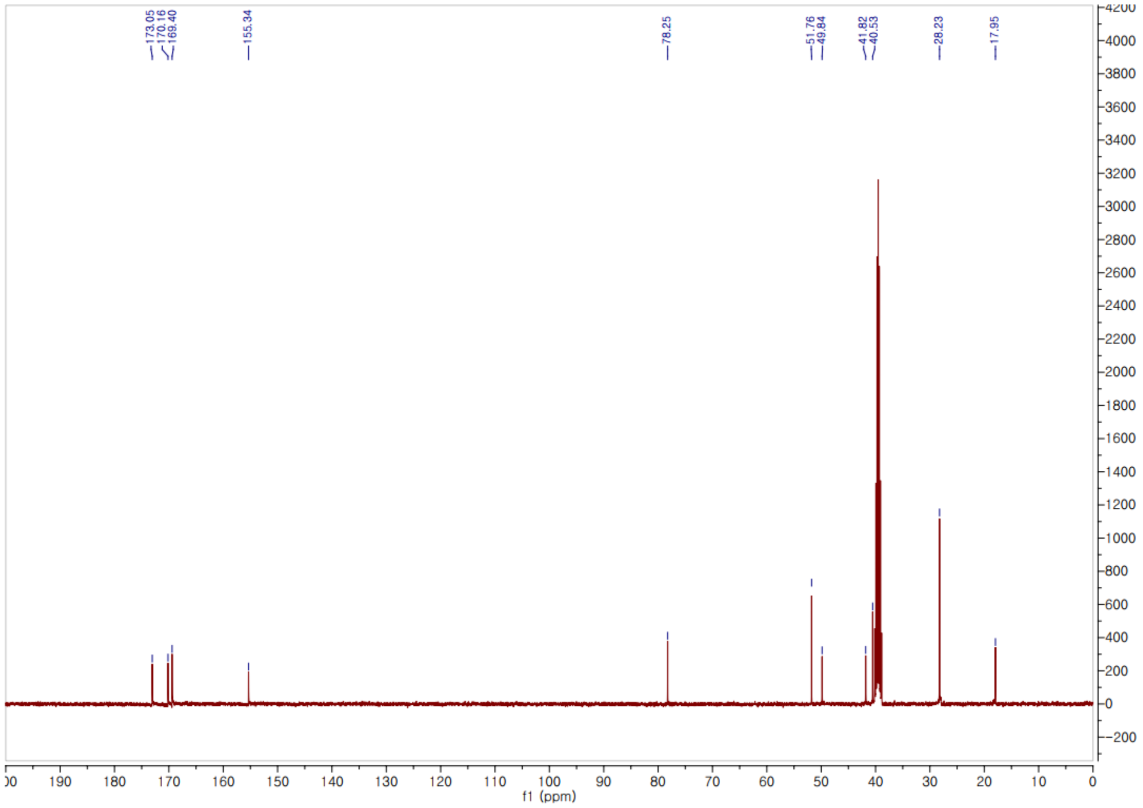

*tert*-Butyl (*S*)-2-((*S*)-2-(methoxycarbonyl)pyrrolidine-1-carbonyl)pyrrolidine-1-carboxylate **(12)**.

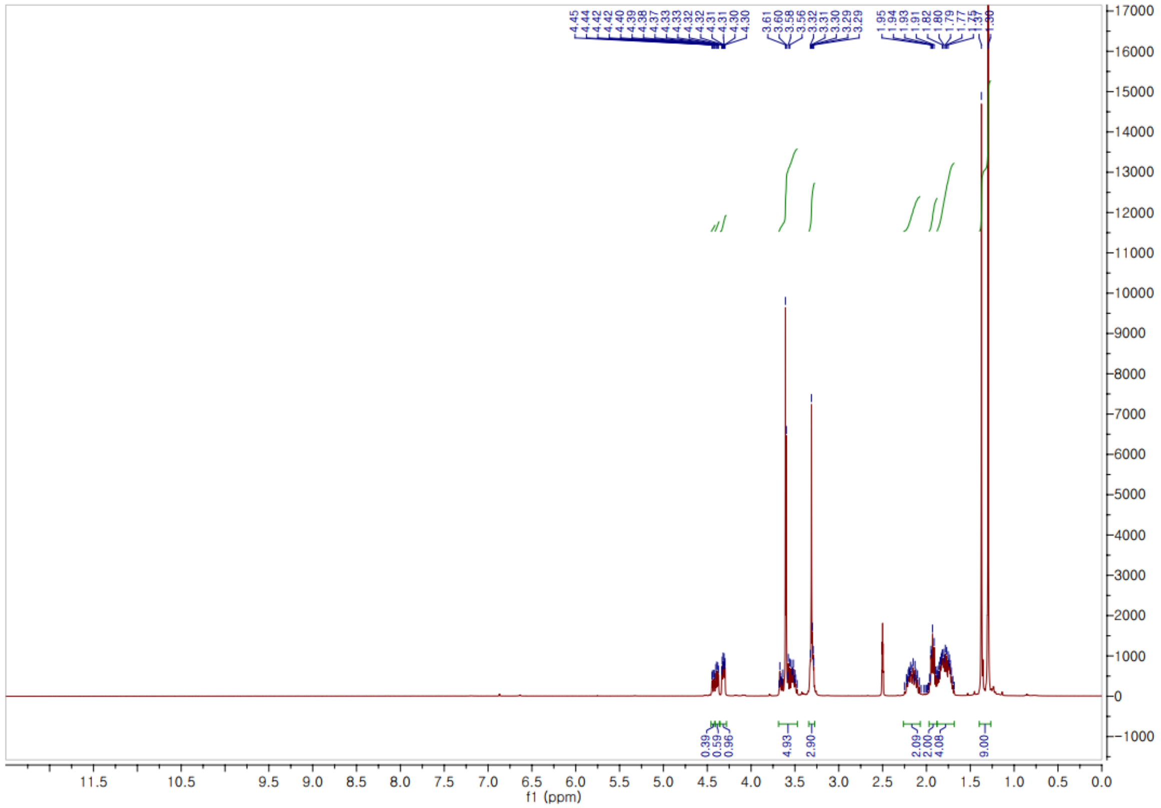


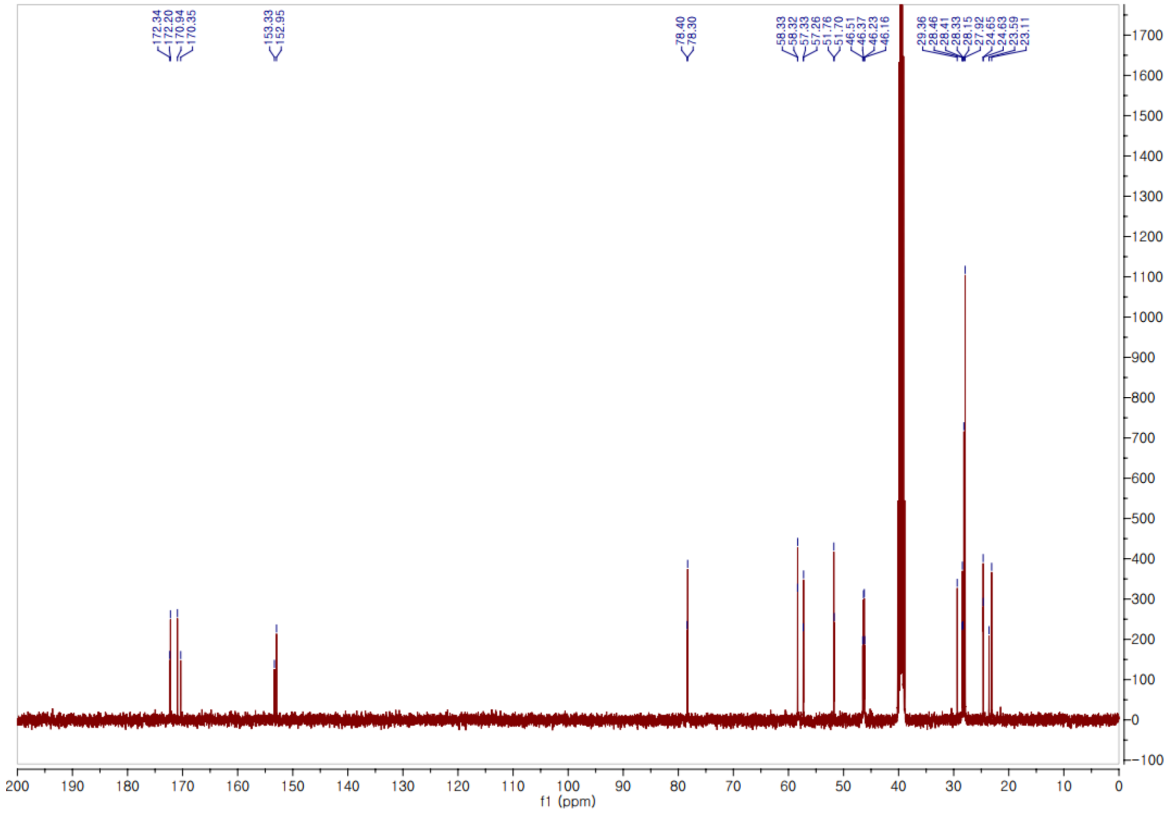

*t*ert-Butyl (*S*)-2-((*S*)-2-((*S*)-2-(methoxycarbonyl)pyrrolidine-1-carbonyl)pyrrolidine-1-carbonyl)pyrrolidine-1-carboxylate **(13)**.

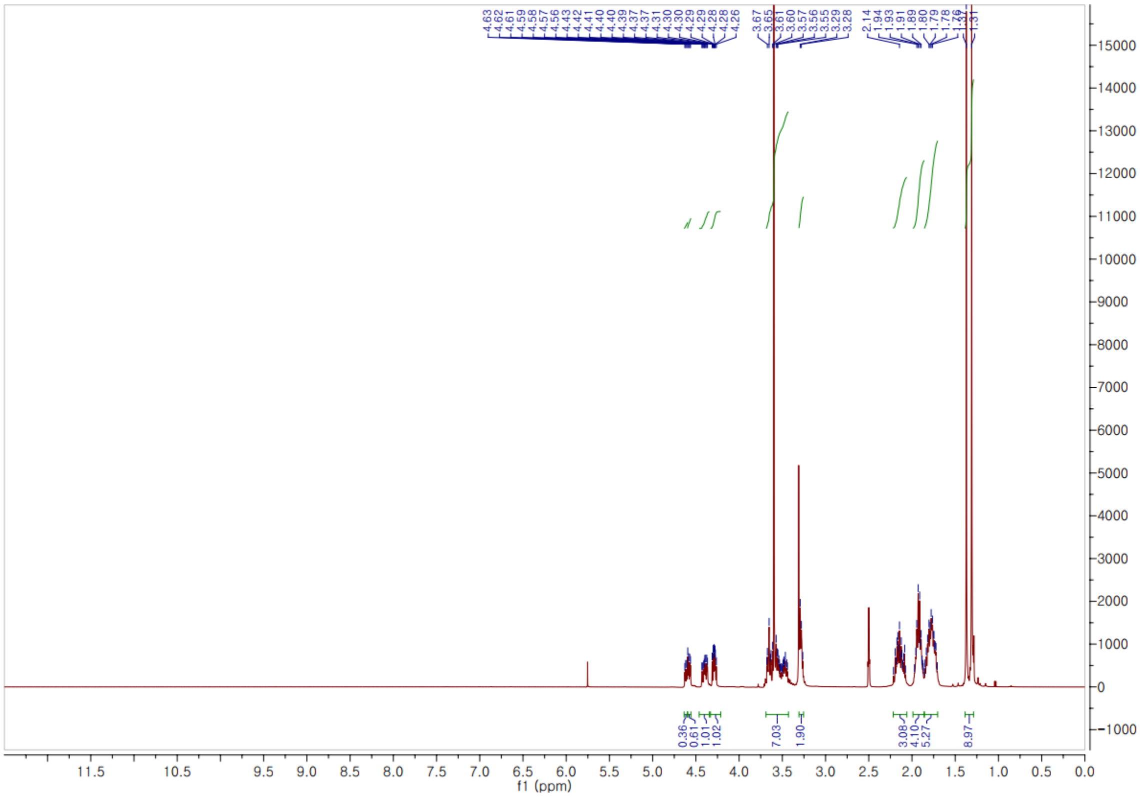


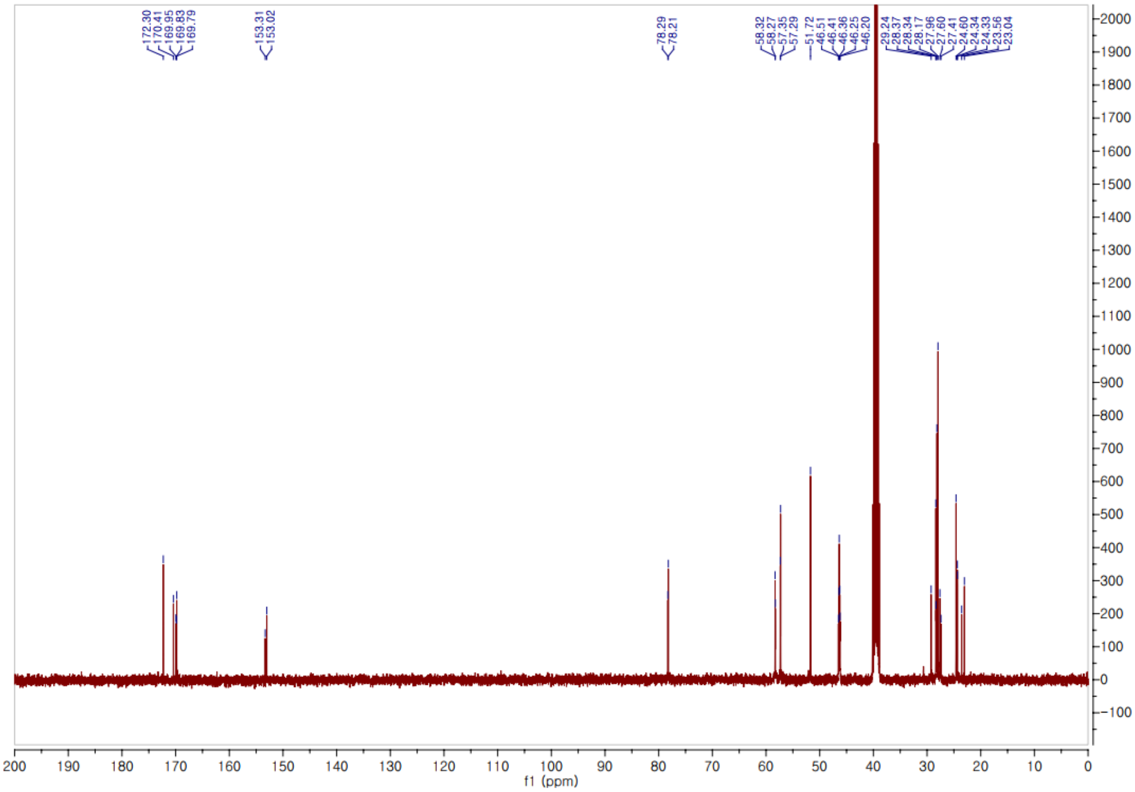

Methyl *O*-methyl-*L*-homoserinate hydrochloride **(16)**.

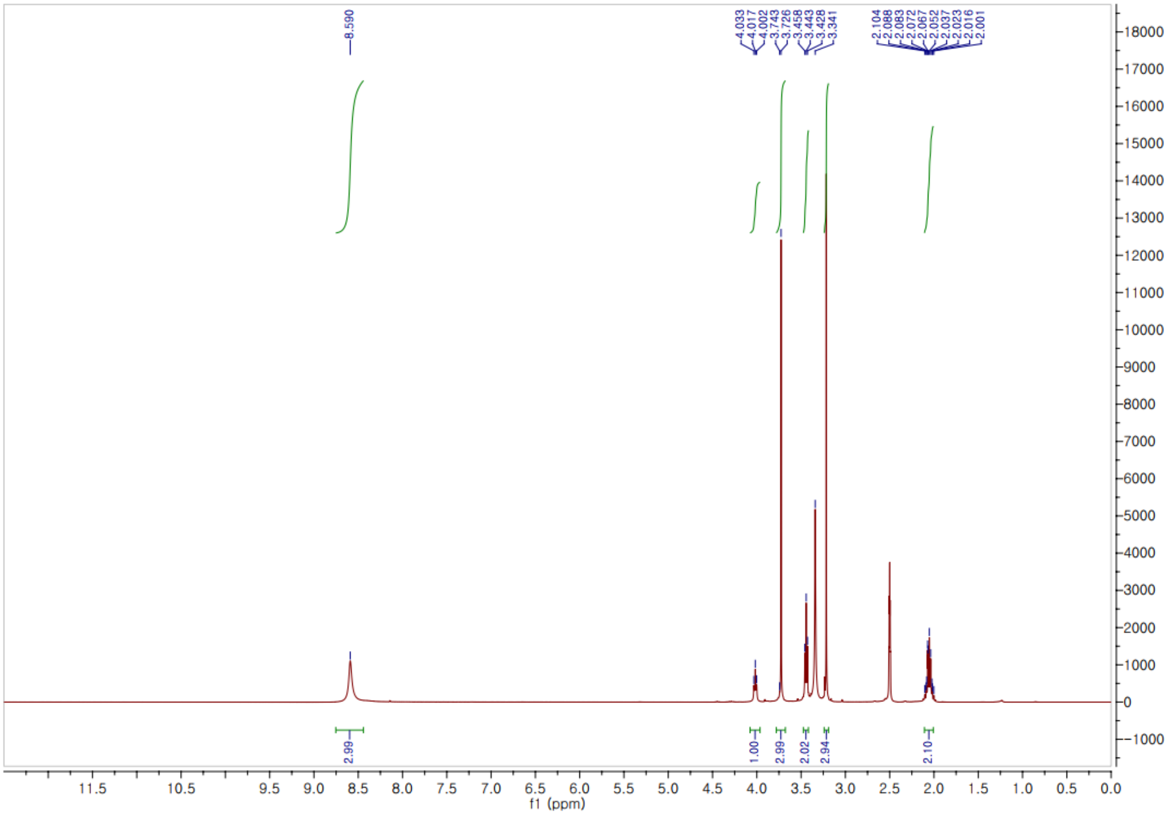


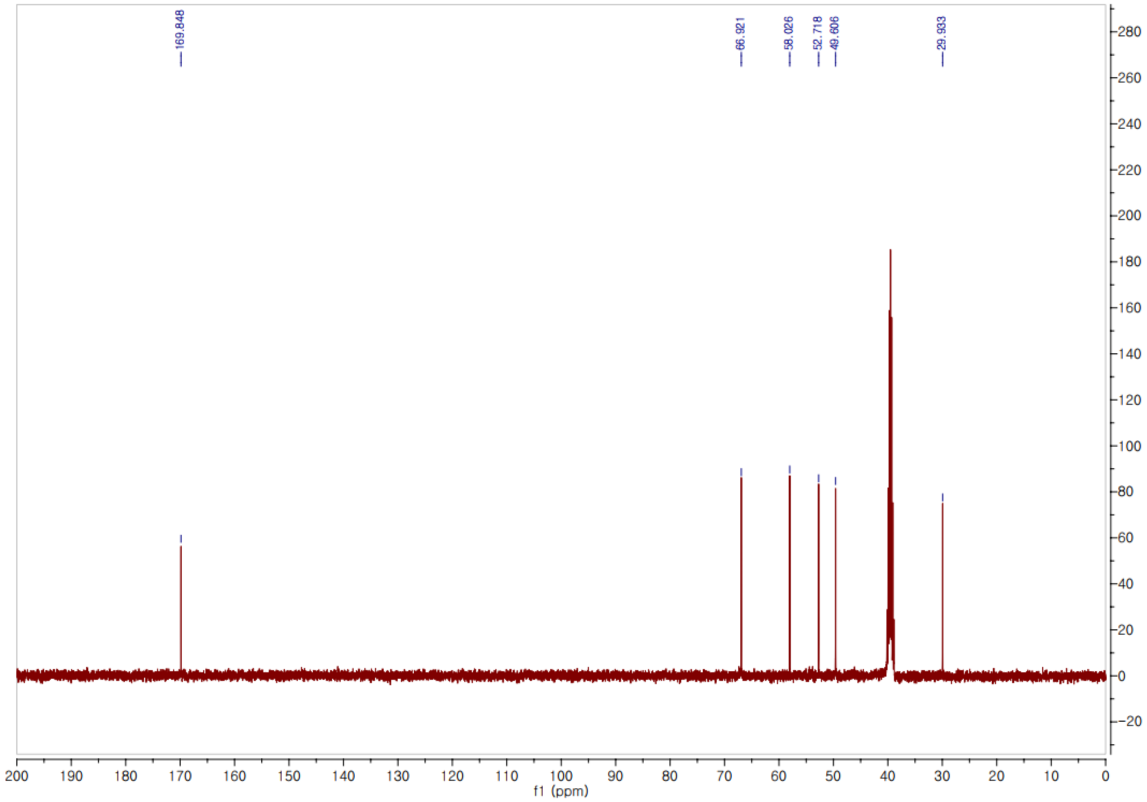

*tert*-Butyl (*S*)-2-((*S*)-2-((*S*)-2-(((*S*)-1-methoxy-4-(methylthio)-1-oxobutan-2-yl)carbamoyl)pyrrolidine-1-carbonyl)pyrrolidine-1-carbonyl)pyrrolidine-1-carboxylate **(17)**.

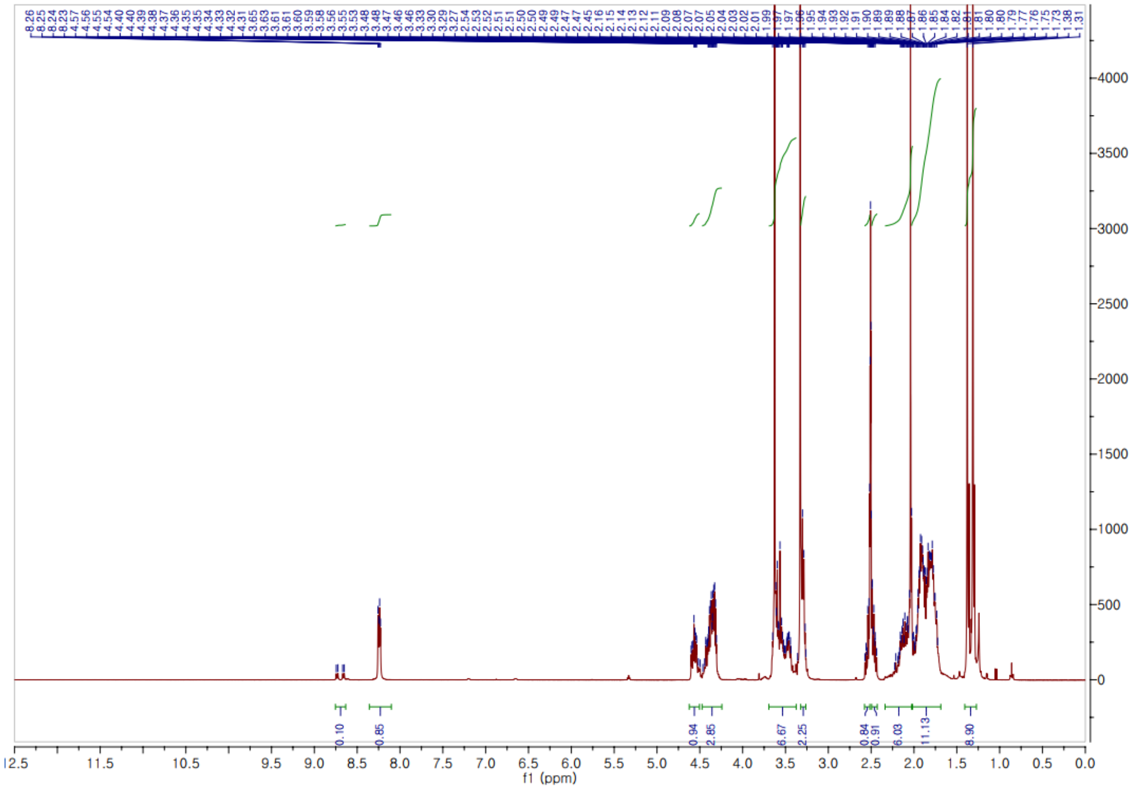


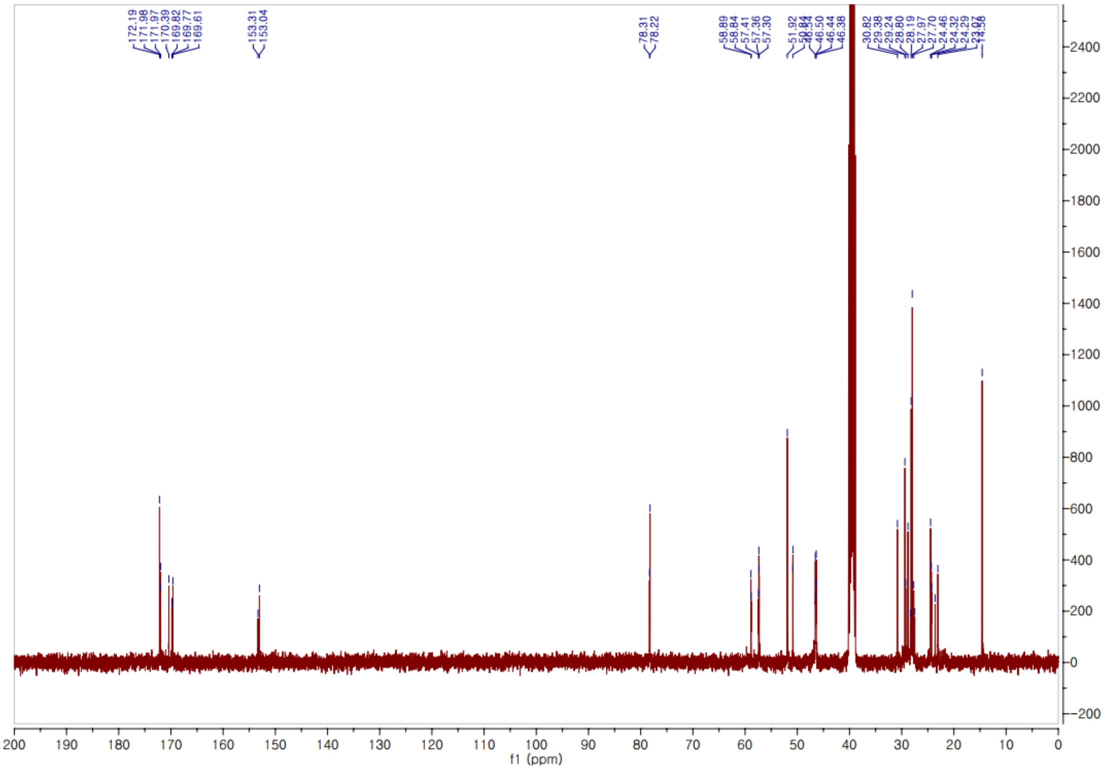

*tert*-Butyl (*S*)-2-((*S*)-2-((*S*)-2-(((*R*)-1-methoxy-4-(methylthio)-1-oxobutan-2-yl)carbamoyl)pyrrolidine-1-carbonyl)pyrrolidine-1-carbonyl)pyrrolidine-1-carboxylate **(18)**.

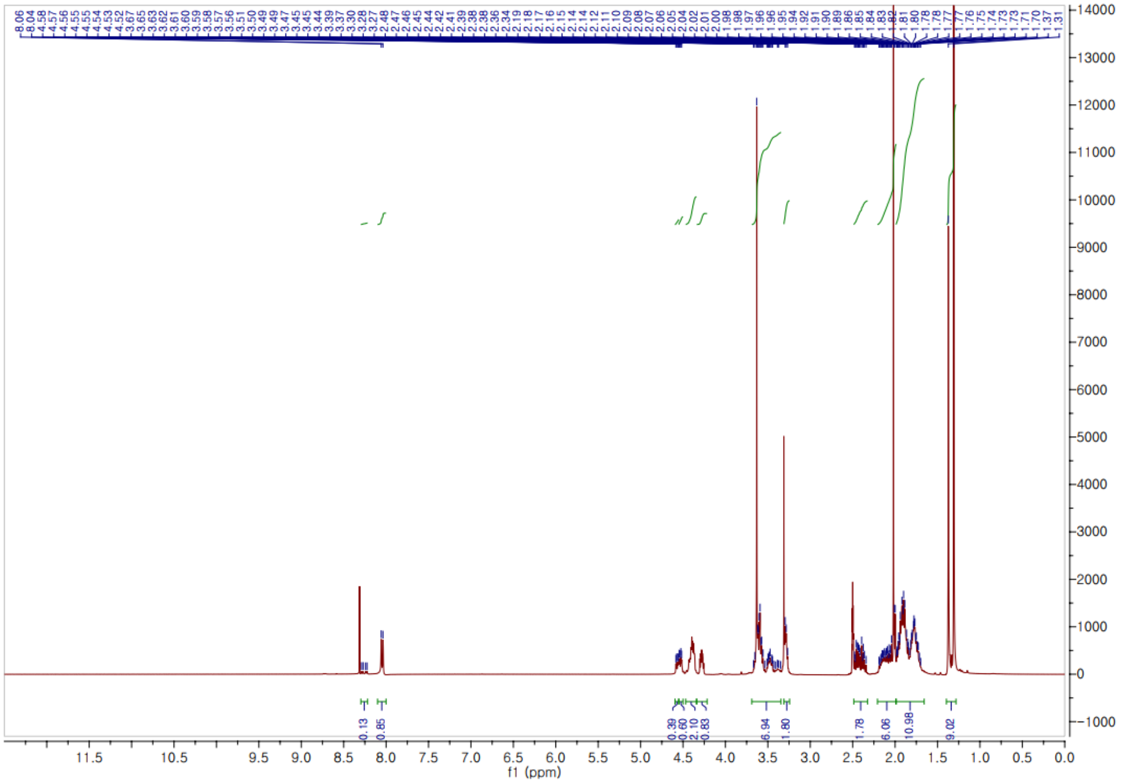

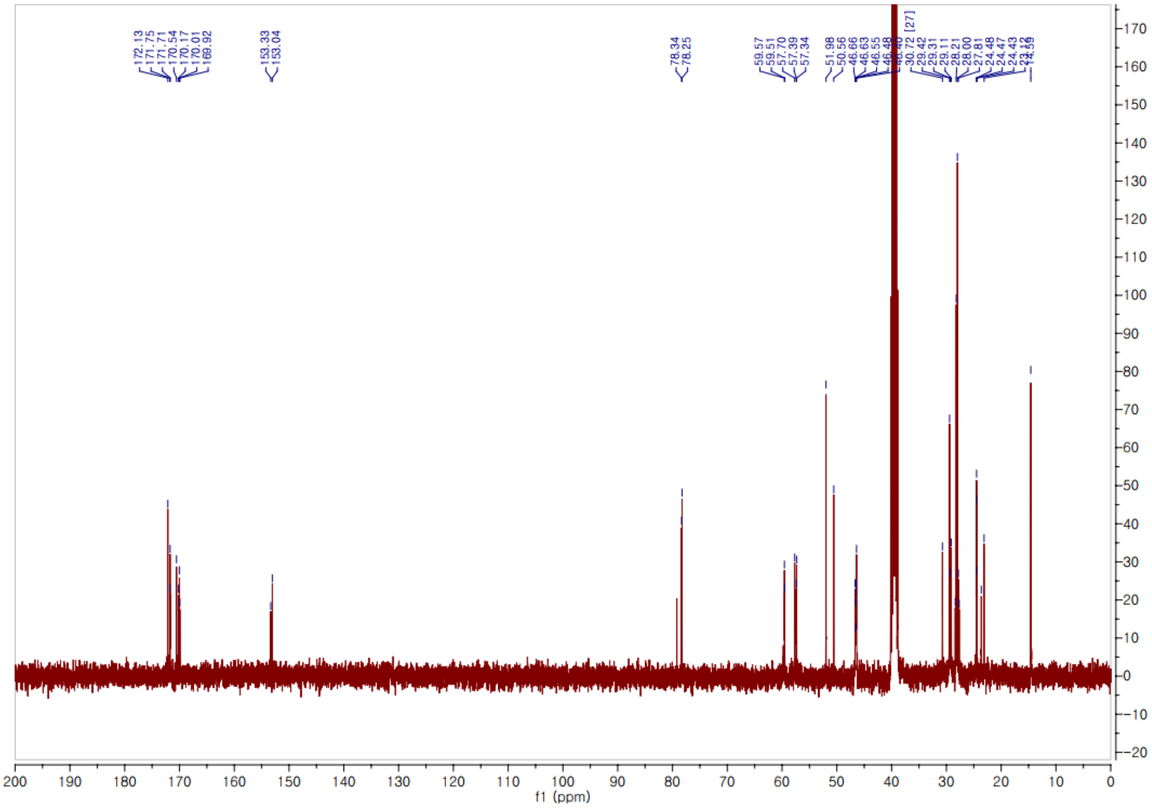

*tert*-Butyl (*S*)-2-((*S*)-2-((*S*)-2-(((*S*)-1,4-dimethoxy-1-oxobutan-2-yl)carbamoyl)pyrrolidine-1-carbonyl)pyrrolidine-1-carbonyl)pyrrolidine-1-carboxylate **(19)**.

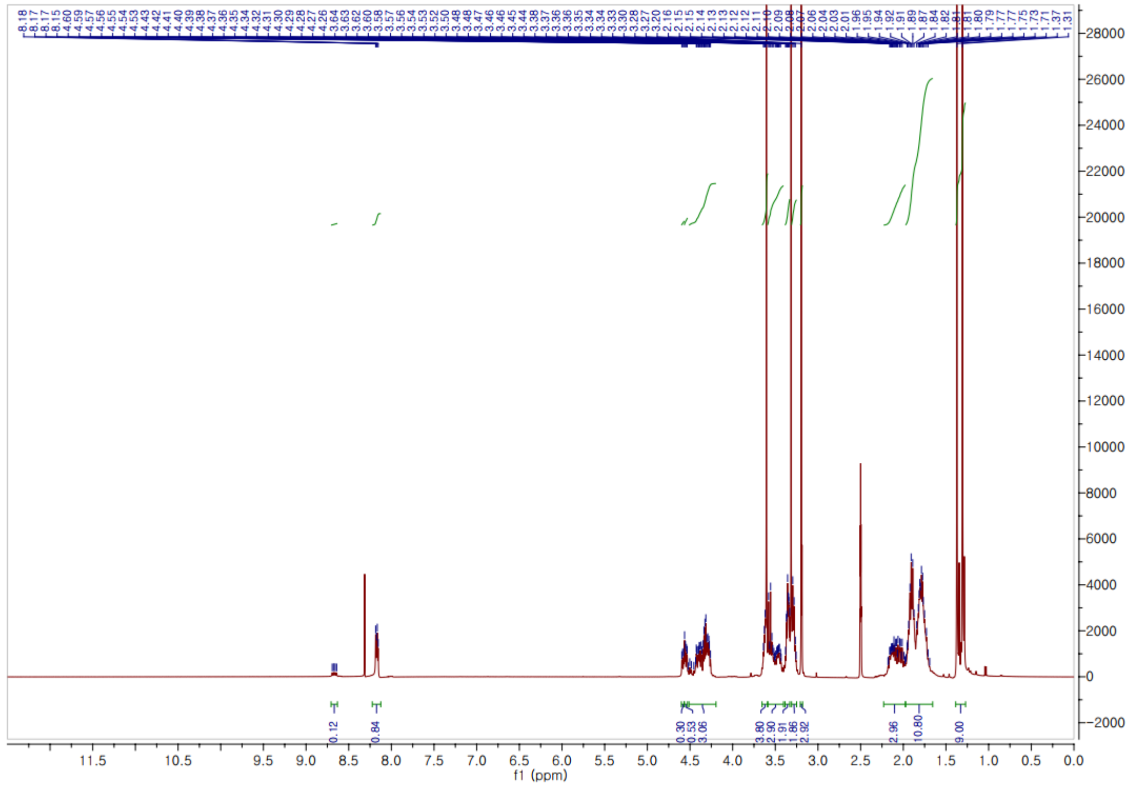


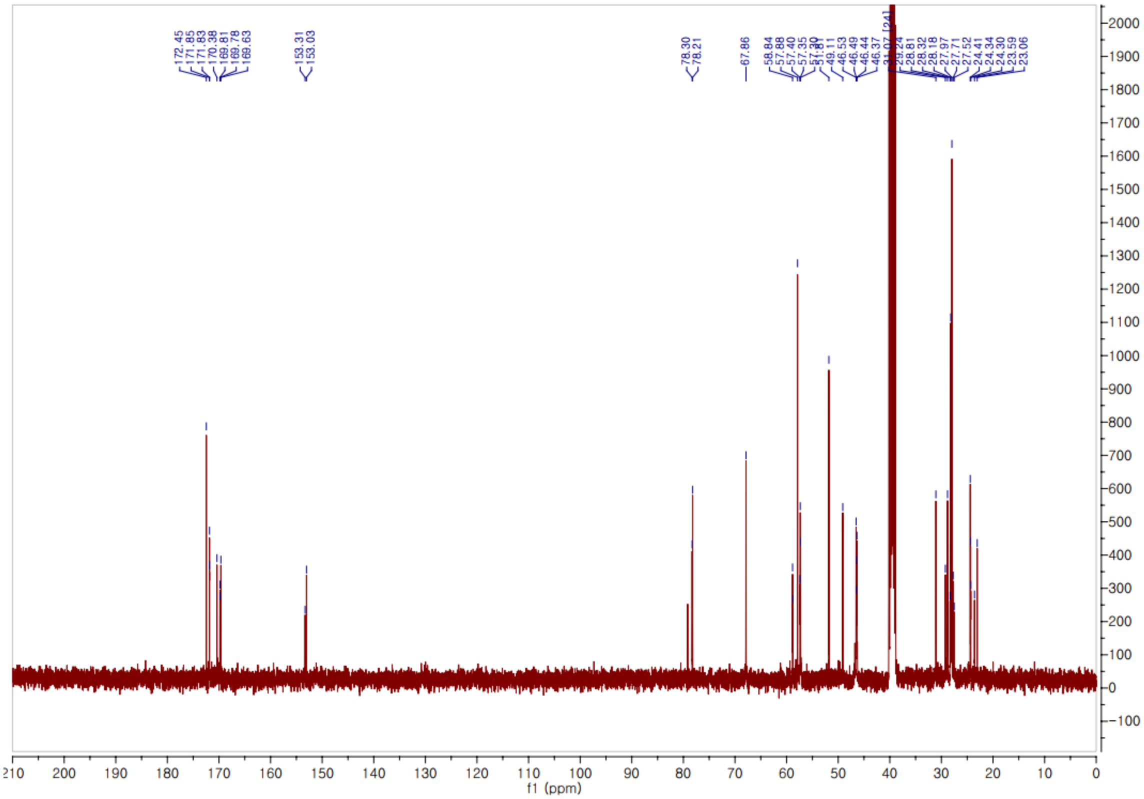

*tert*-Butyl (*S*)-2-(((*S*)-1-methoxy-4-(methylthio)-1-oxobutan-2-yl)carbamoyl)pyrrolidine-1-carboxylate **(20)**.

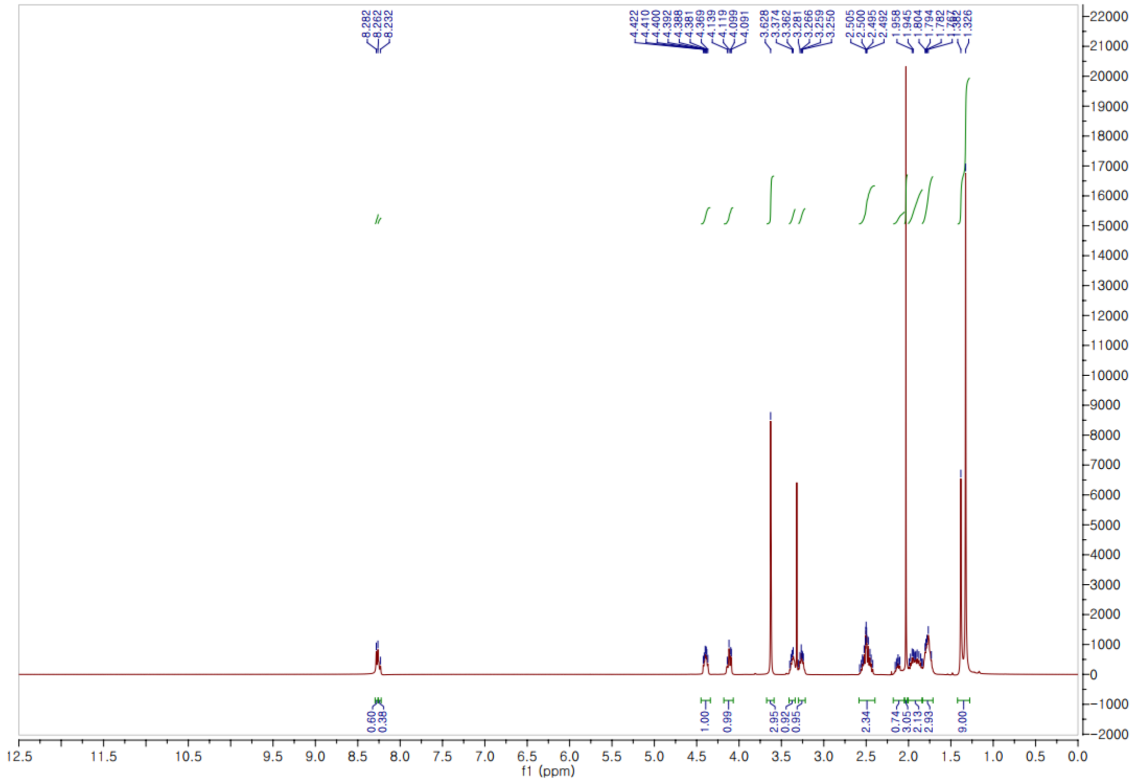


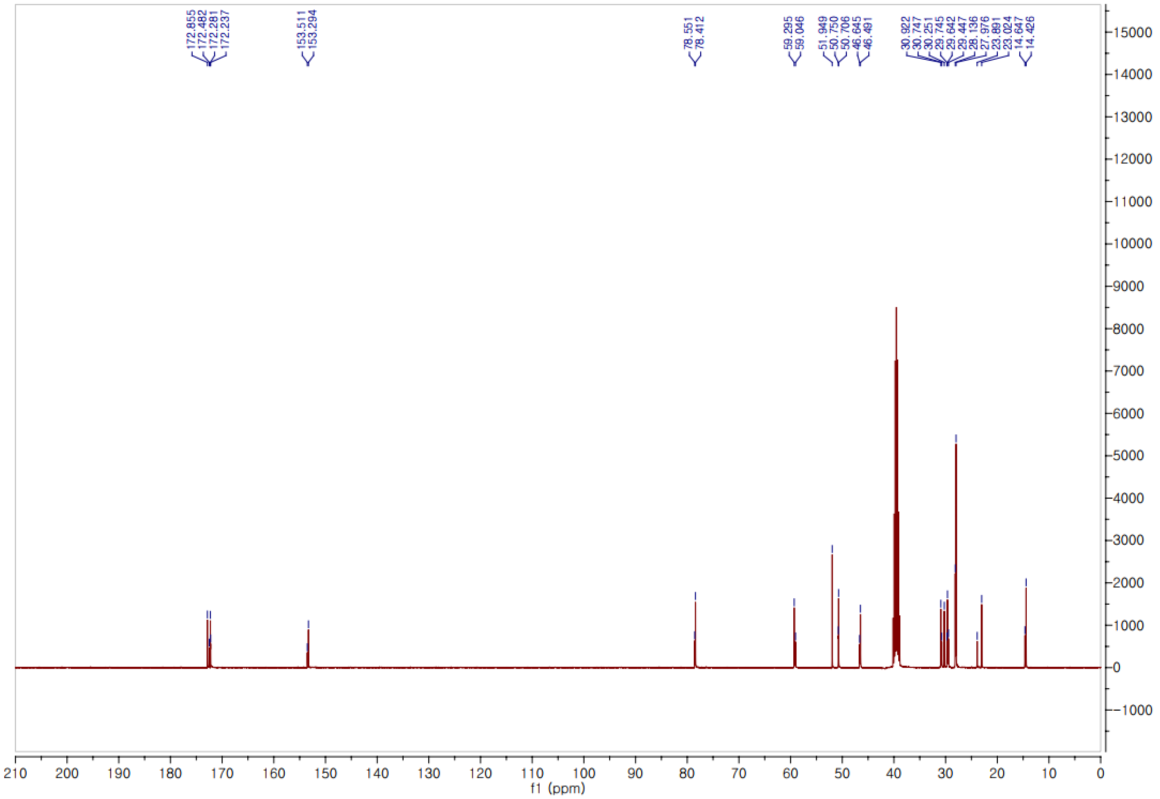

*tert*-Butyl (*S*)-2-(((10*S*,13*S*)-10-methyl-3,6,9,12-tetraoxo-2-oxa-16-thia-5,8,11-triazaheptadecan-13-yl)carbamoyl)pyrrolidine-1-carboxylate **(21)**.

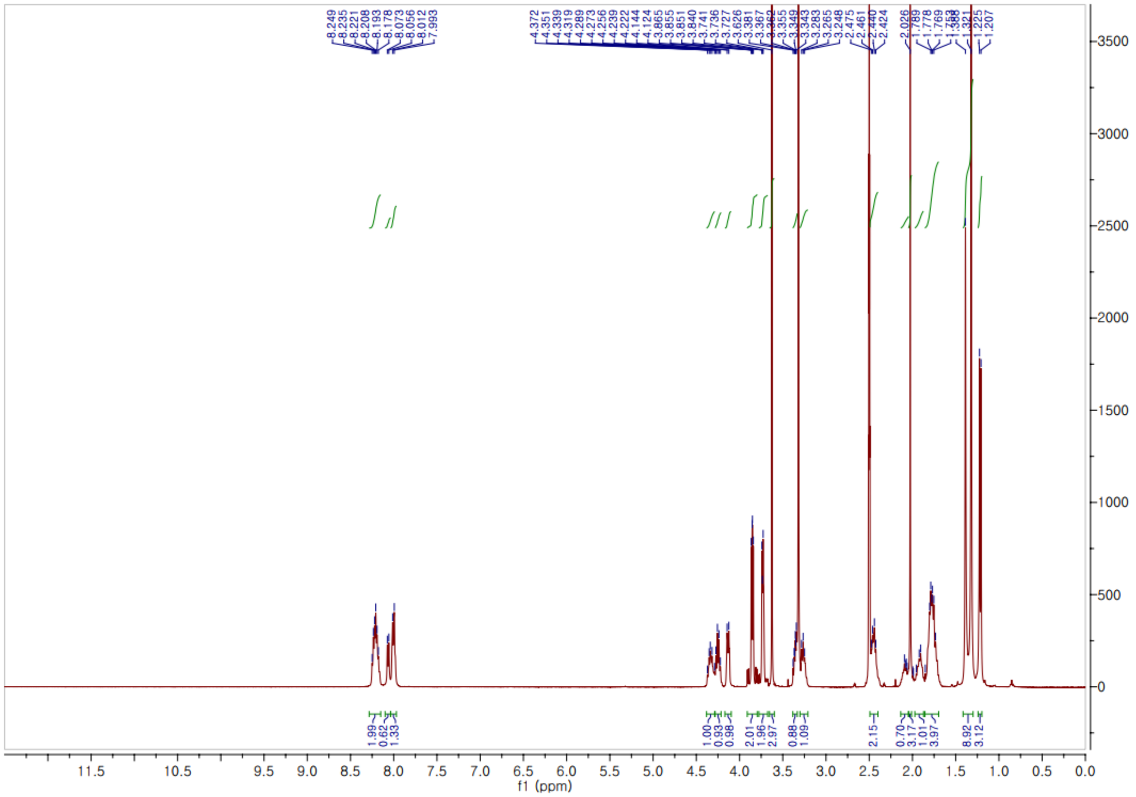


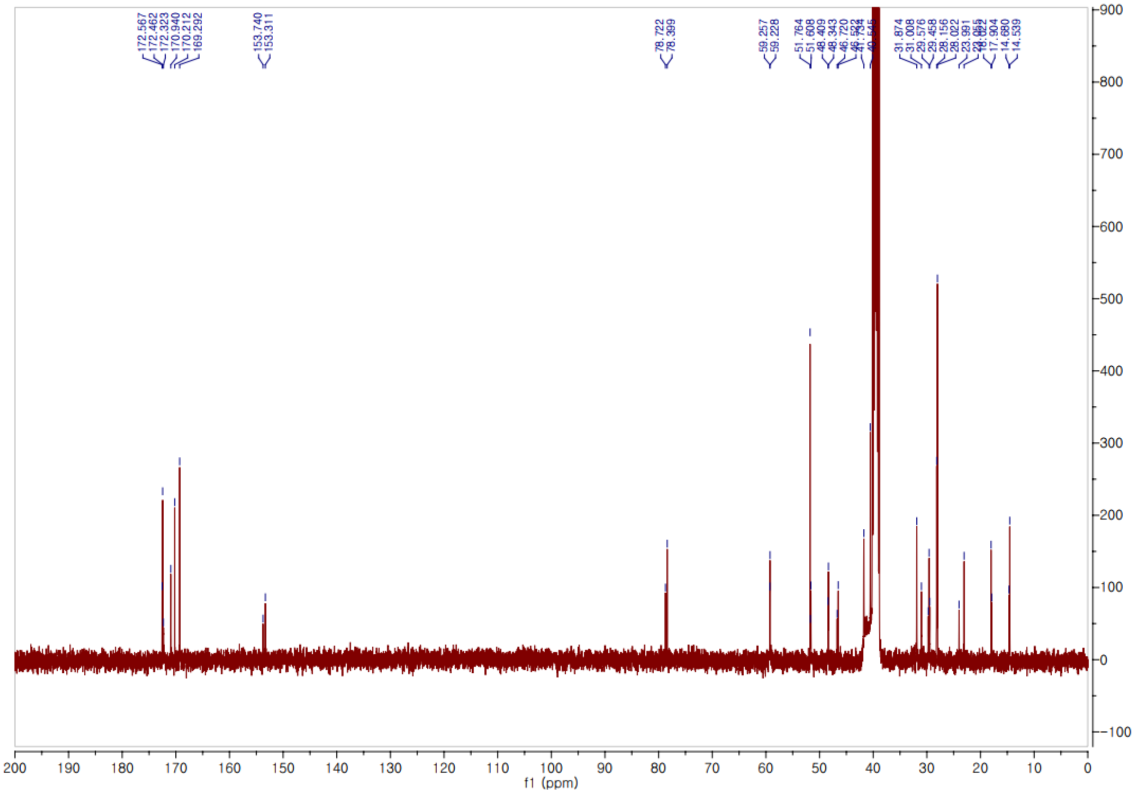

*tert*-Butyl (*S*)-2-((*S*)-2-((*S*)-2-(((10*S*,13*S*)-10-methyl-3,6,9,12-tetraoxo-2-oxa-16-thia-5,8,11-triazaheptadecan-13-yl)carbamoyl)pyrrolidine-1-carbonyl)pyrrolidine-1-carbonyl)pyrrolidine-1-carboxylate **(22)**.

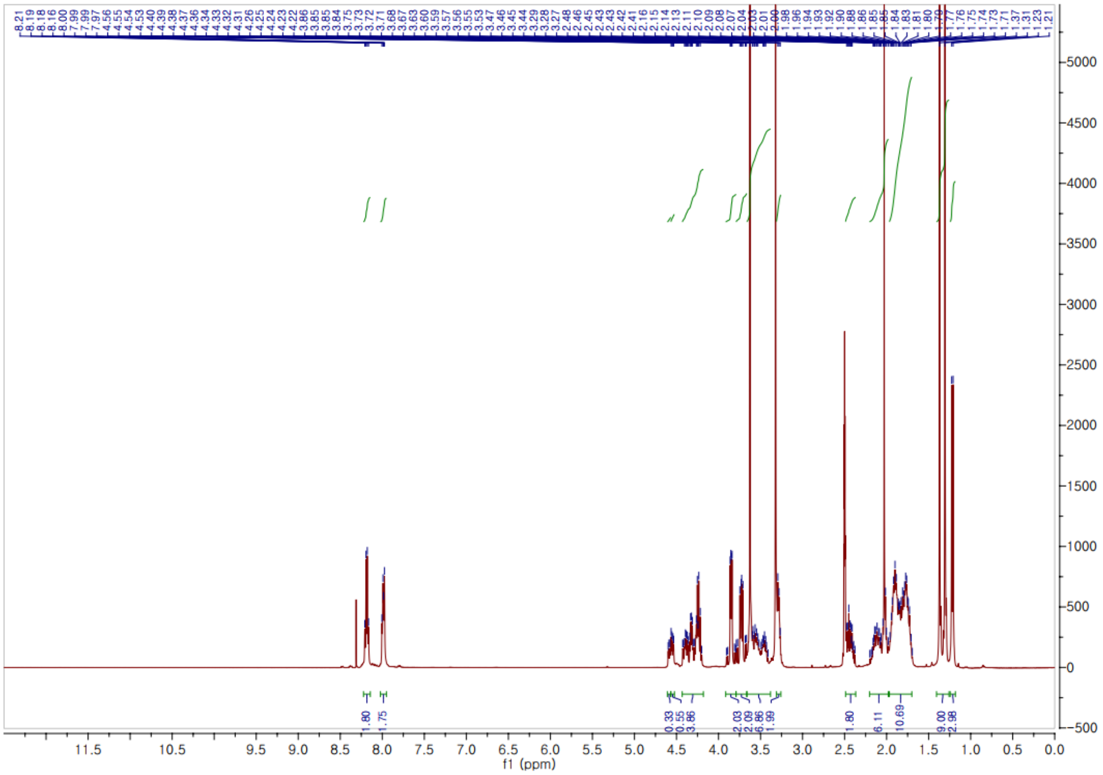


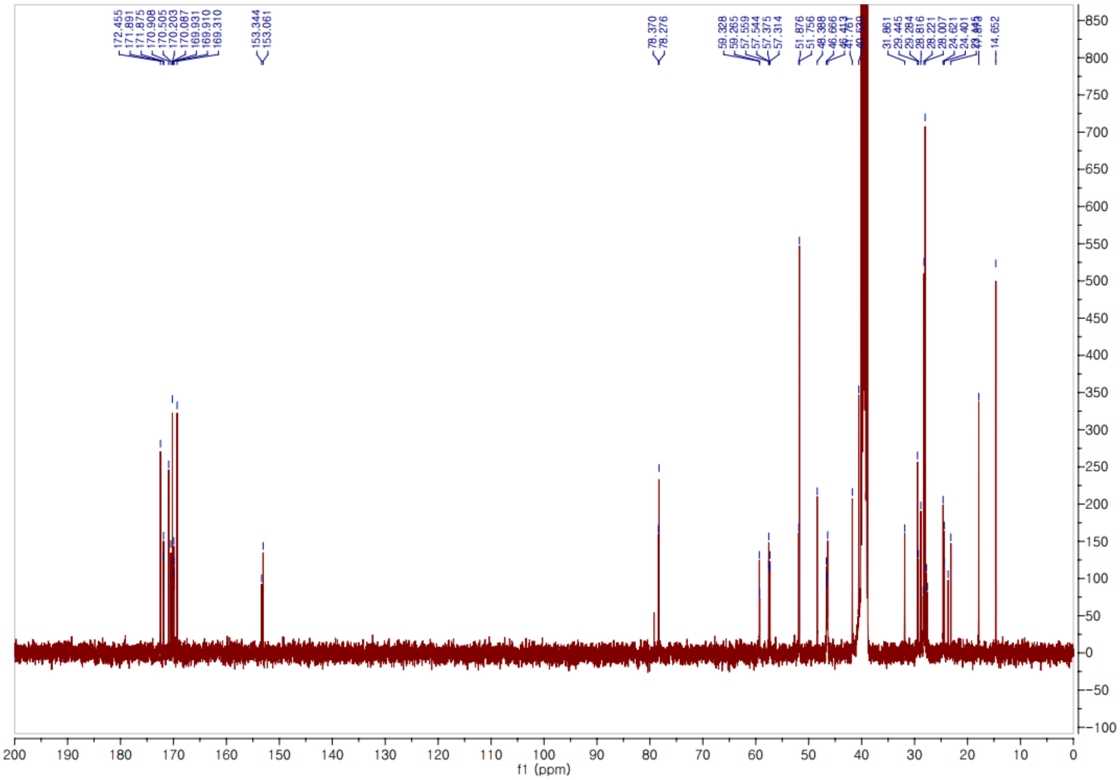

*tert*-Butyl (*S*)-2-((*S*)-2-((*S*)-2-(((10*S*,13*R*)-10-methyl-3,6,9,12-tetraoxo-2-oxa-16-thia-5,8,11-triazaheptadecan-13-yl)carbamoyl)pyrrolidine-1-carbonyl)pyrrolidine-1-carbonyl)pyrrolidine-1-carboxylate **(23)**.

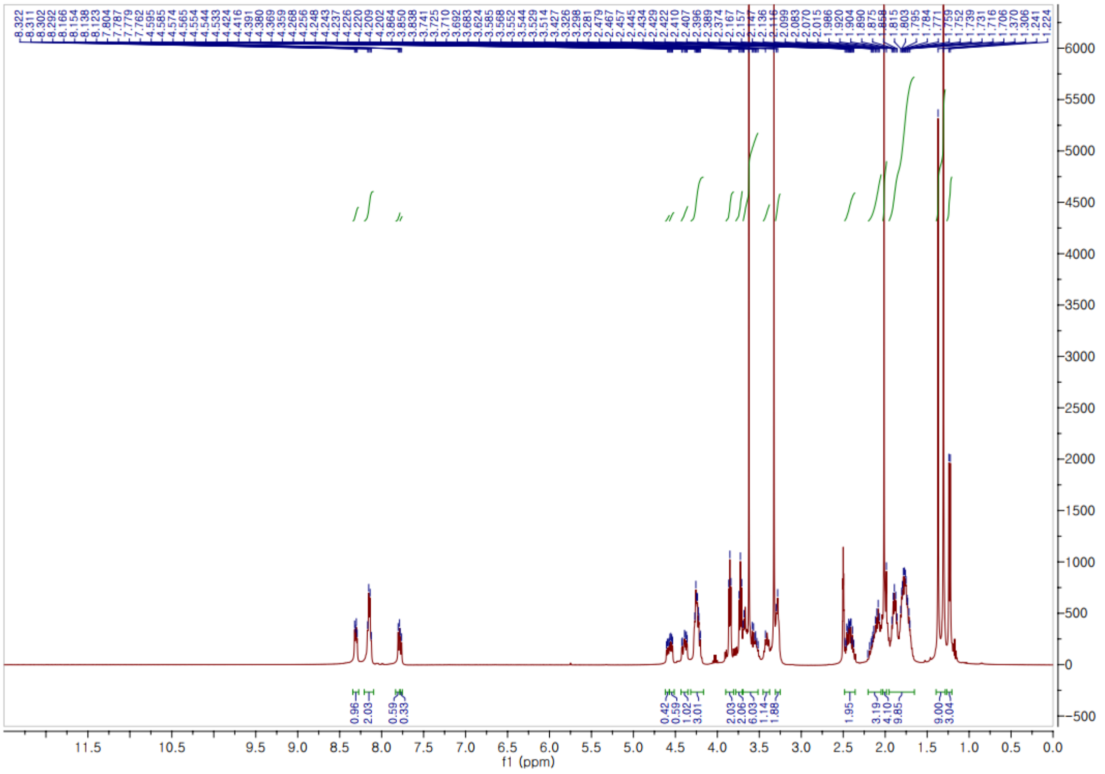


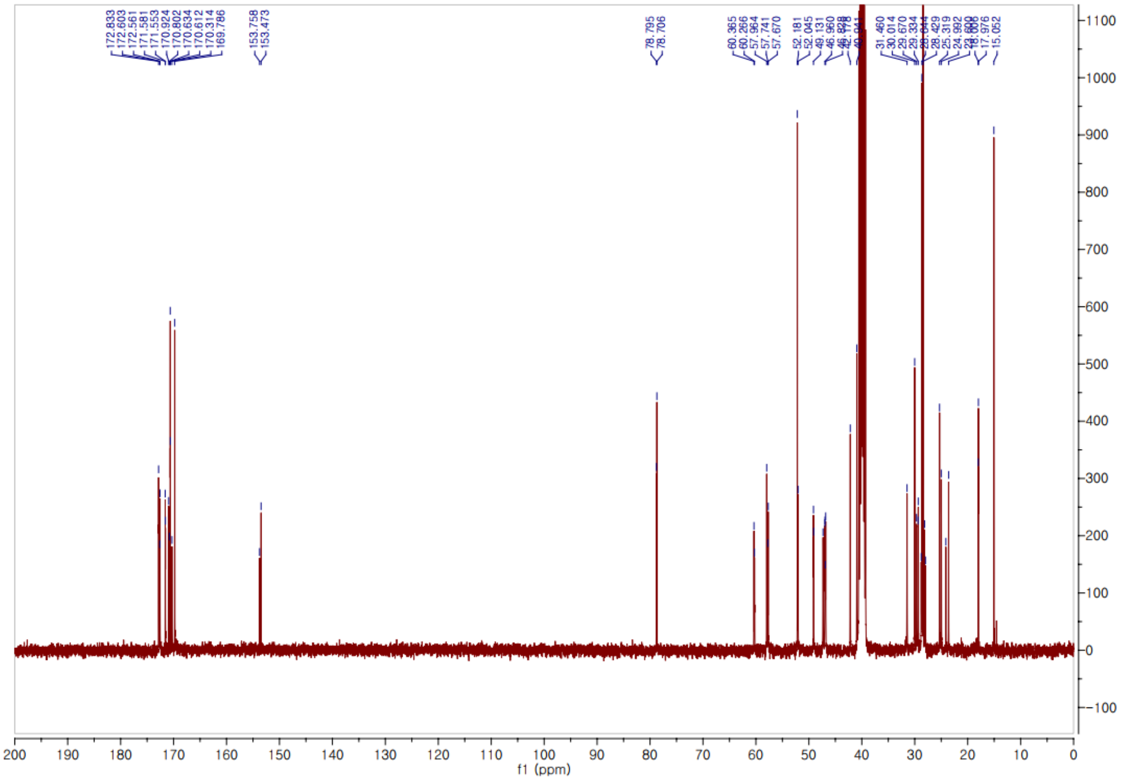

*tert*-Butyl (*S*)-2-((*S*)-2-((*S*)-2-(((10*S*,13*S*)-10-methyl-3,6,9,12-tetraoxo-2,16-dioxa-5,8,11-triazaheptadecan-13-yl)carbamoyl)pyrrolidine-1-carbonyl)pyrrolidine-1-carbonyl)pyrrolidine-1-carboxylate **(24)**.

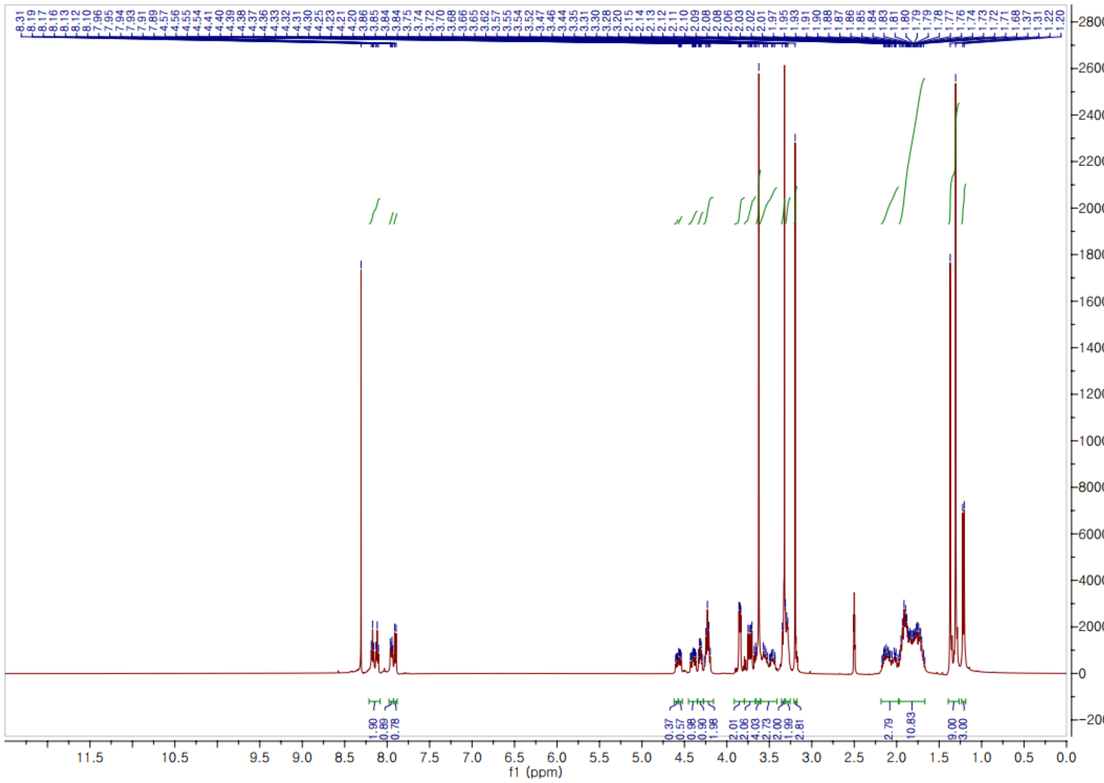


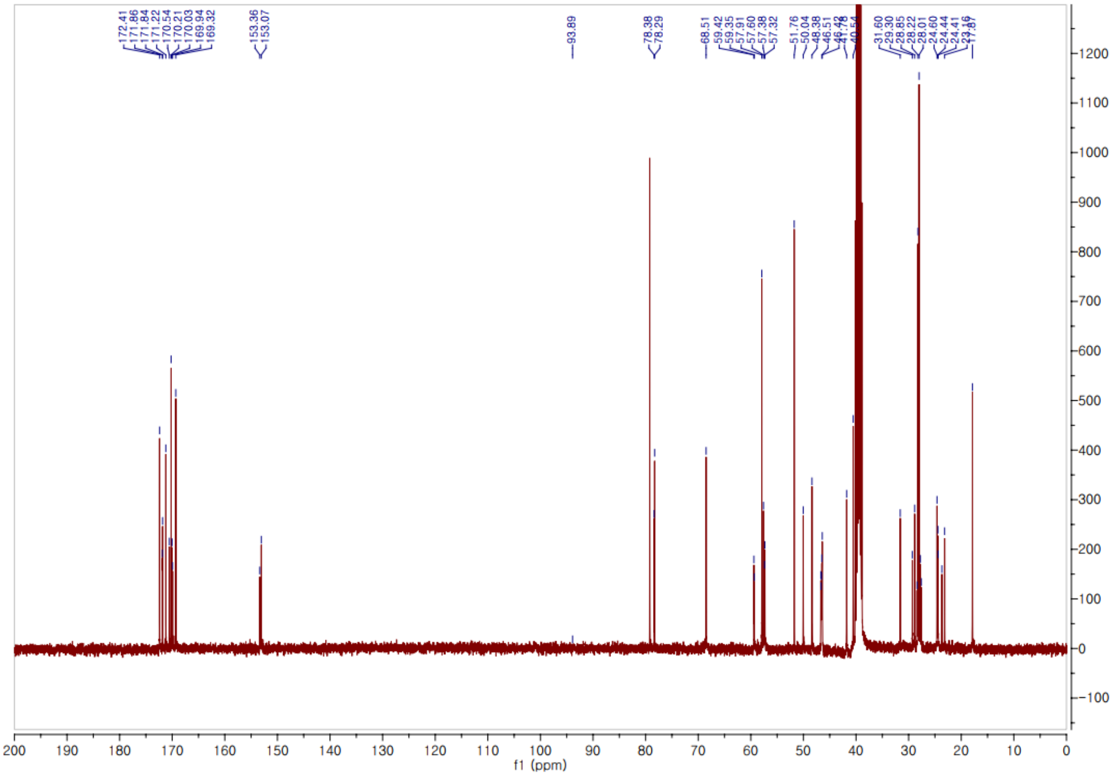

**$^{1}$H, $^{13}$C NMR, HPLC traces, and ESI-HRMS spectra of final compounds**

(3-(2-(2-(4-(4-((7-(3-(methylsulfonamido)phenyl)thieno[3,2-d]pyrimidin-2-yl)amino)phenyl)piperazin-1-yl)ethoxy)ethoxy)propanoyl)-*L*-prolyl-*L*-methionyl-*L*-alanylglycylglycine **(25)**.

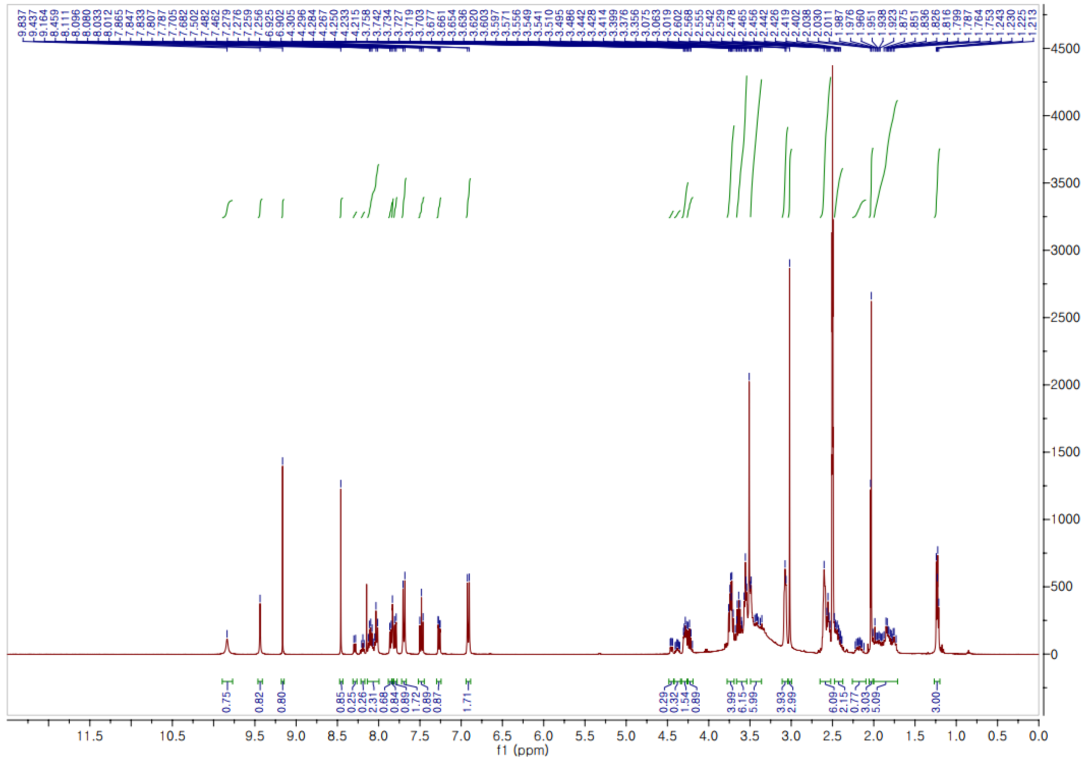


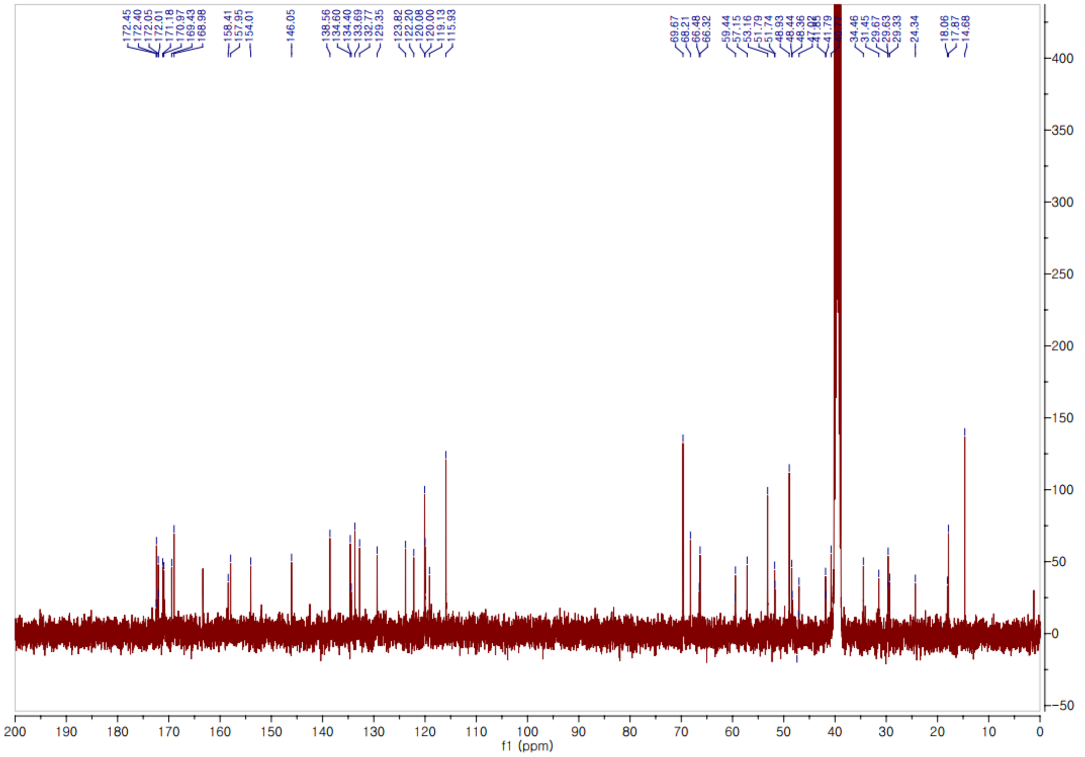

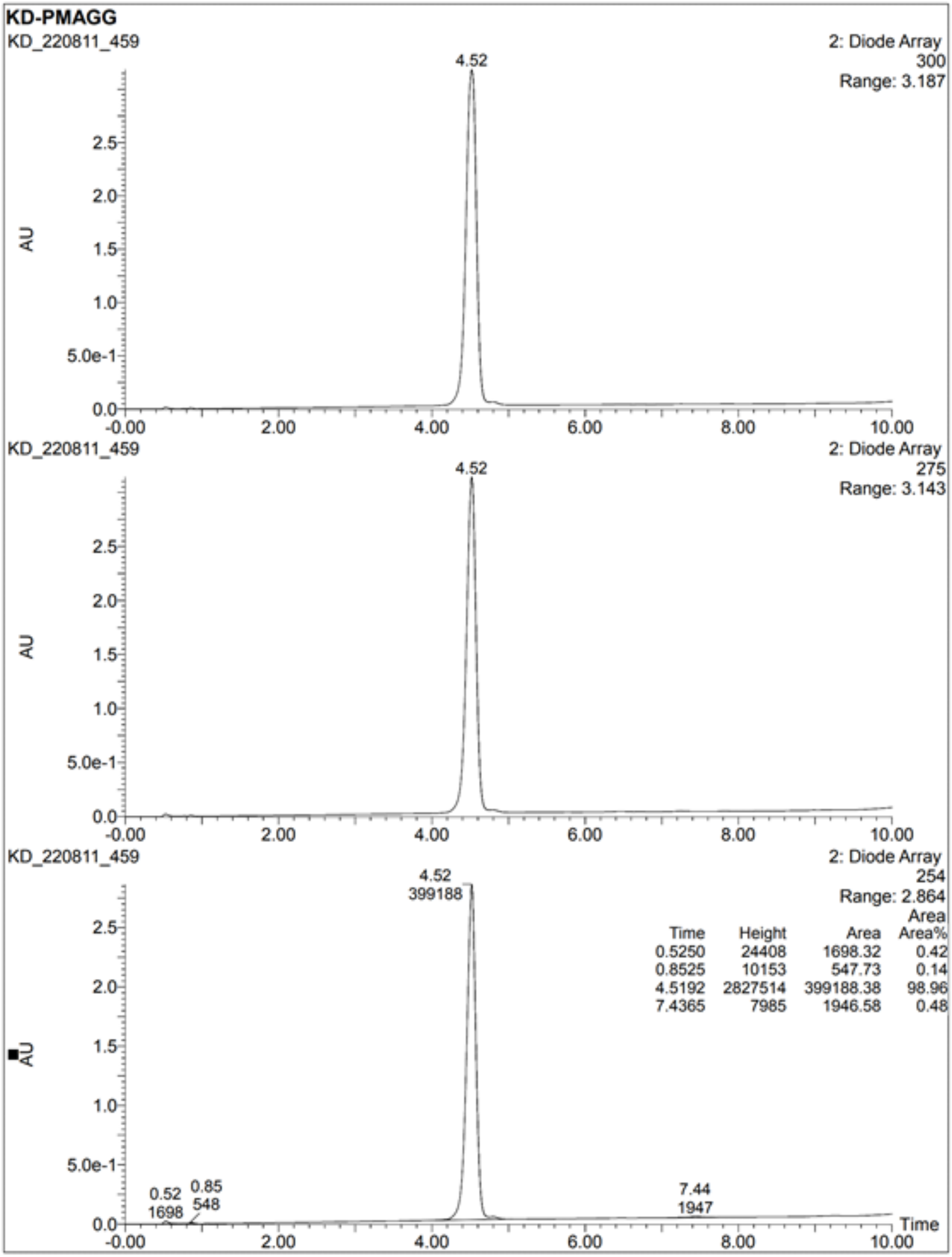
KD-PMAGG
KD_220811_459
2: Diode Array
300
Range: 3.187
4.52
AU
KD_220811_459
2: Diode Array
275
Range: 3.143
4.52
AU
KD_220811_459
2: Diode Array
254
Range: 2.864
4.52
399188
Time Height Area Area%
0.5250 24408 1698.32 0.42
0.8525 10153 547.73 0.14
4.5192 2827514 399188.38 98.96
7.4365 7985 1946.58 0.48
0.52
1698
0.85
548
7.44
1947
AU
Time

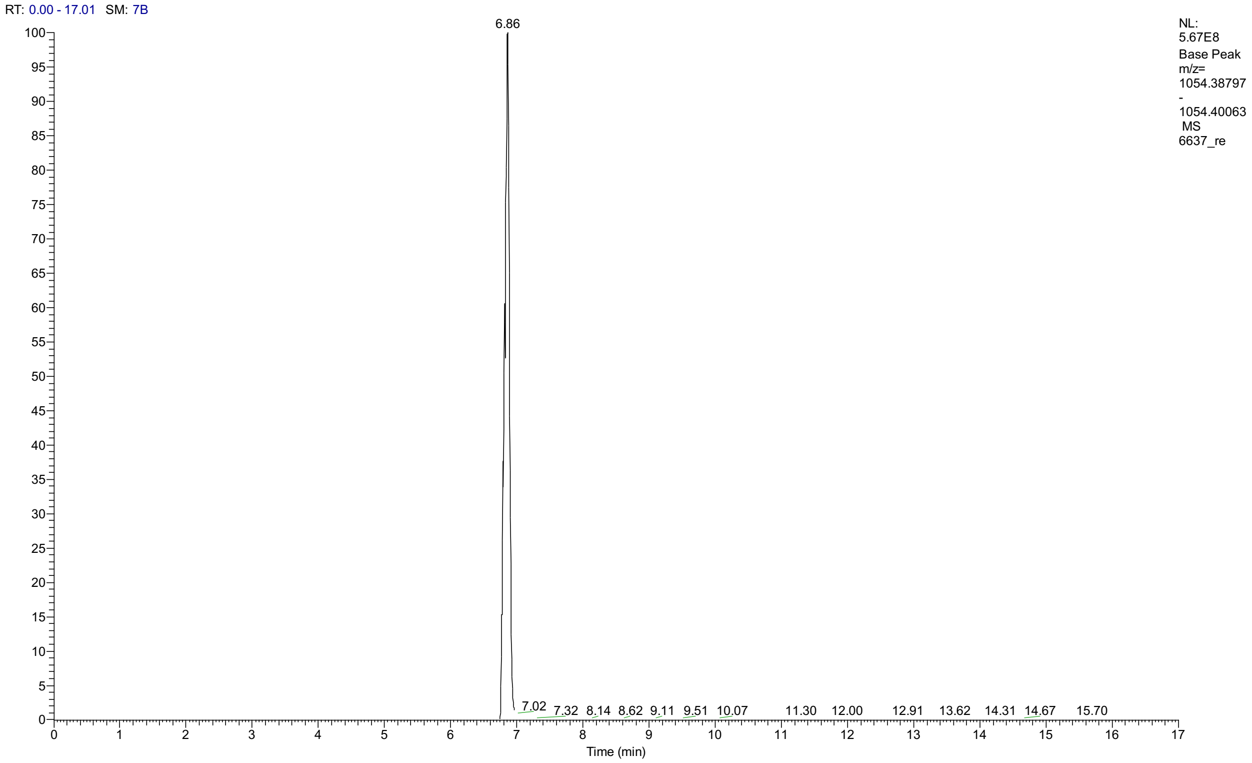
RT: 0.00 - 17.01 SM: 7B
6.86
NL:
5.67E8
Base Peak
m/z=
1054.38797
-
1054.40063
MS
6637_re
7.02
7.32
8.14
8.62
9.11
9.51
10.07
11.30
12.00
12.91
13.62
14.31
14.67
15.70
Time (min)

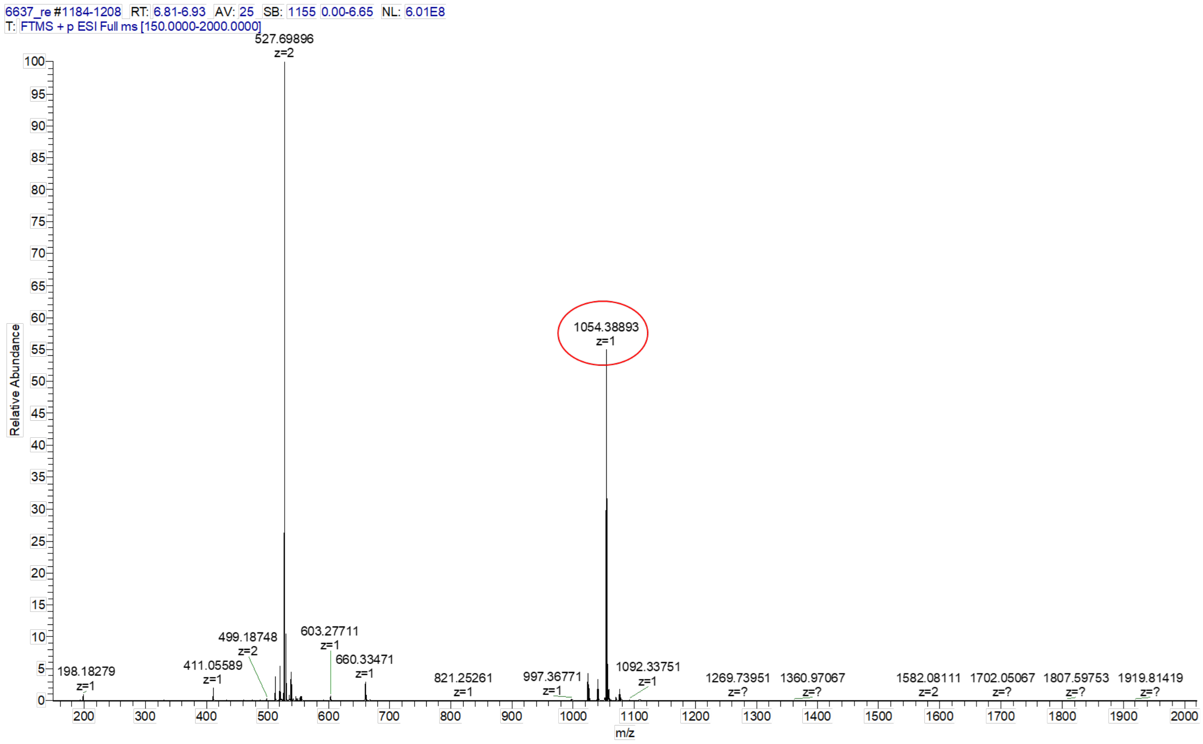
6637_re #1184-1208 RT: 6.81-6.93 AV: 25 SB: 1155 0.00-6.65 NL: 6.01E8
T: FTMS + p ESI Full ms [150.0000-2000.0000]
527.69896
z=2
1054.38893
z=1
Relative Abundance
499.18748
z=2
603.27711
z=1
660.33471
z=1
198.18279
z=1
411.05589
z=1
821.25261
z=1
997.36771
z=1
1092.33751
z=1
1269.73951
z=?
1360.97067
z=?
1582.08111
z=2
1702.05067
z=?
1807.59753
z=?
1919.81419
z=?
m/z

(3-(2-(2-(4-(4-((7-(3-(methylsulfonamido)phenyl)thieno[3,2-d]pyrimidin-2-yl)amino)phenyl)piperazin-1-yl)ethoxy)ethoxy)propanoyl)-*L*-prolyl-*L*-prolyl-*L*-prolyl-*L*-methionyl-*L*-alanylglycylglycine **(26)**.

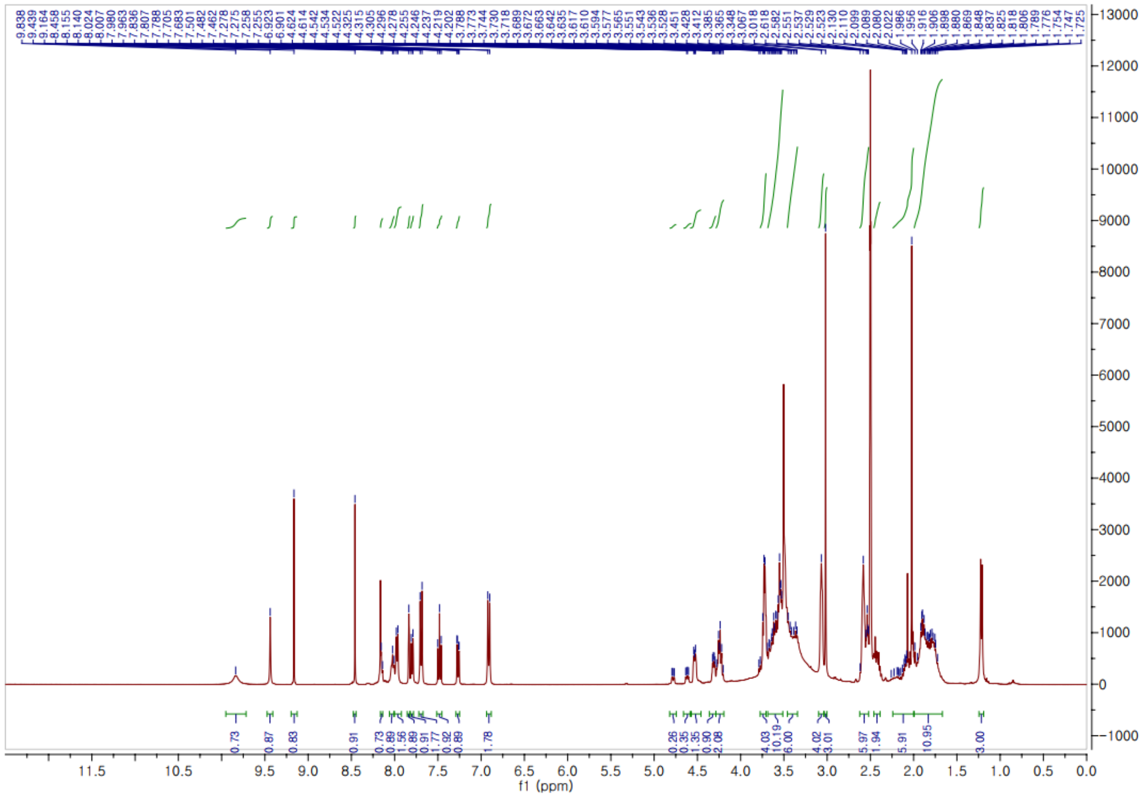

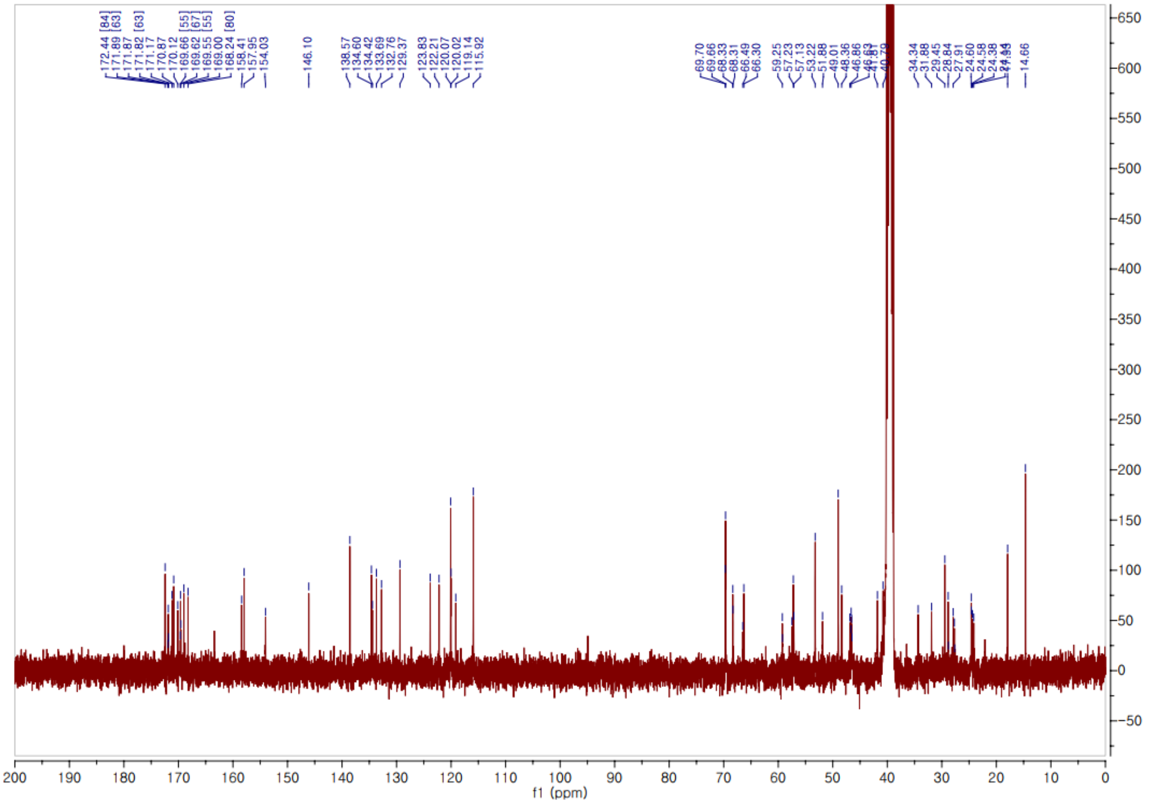

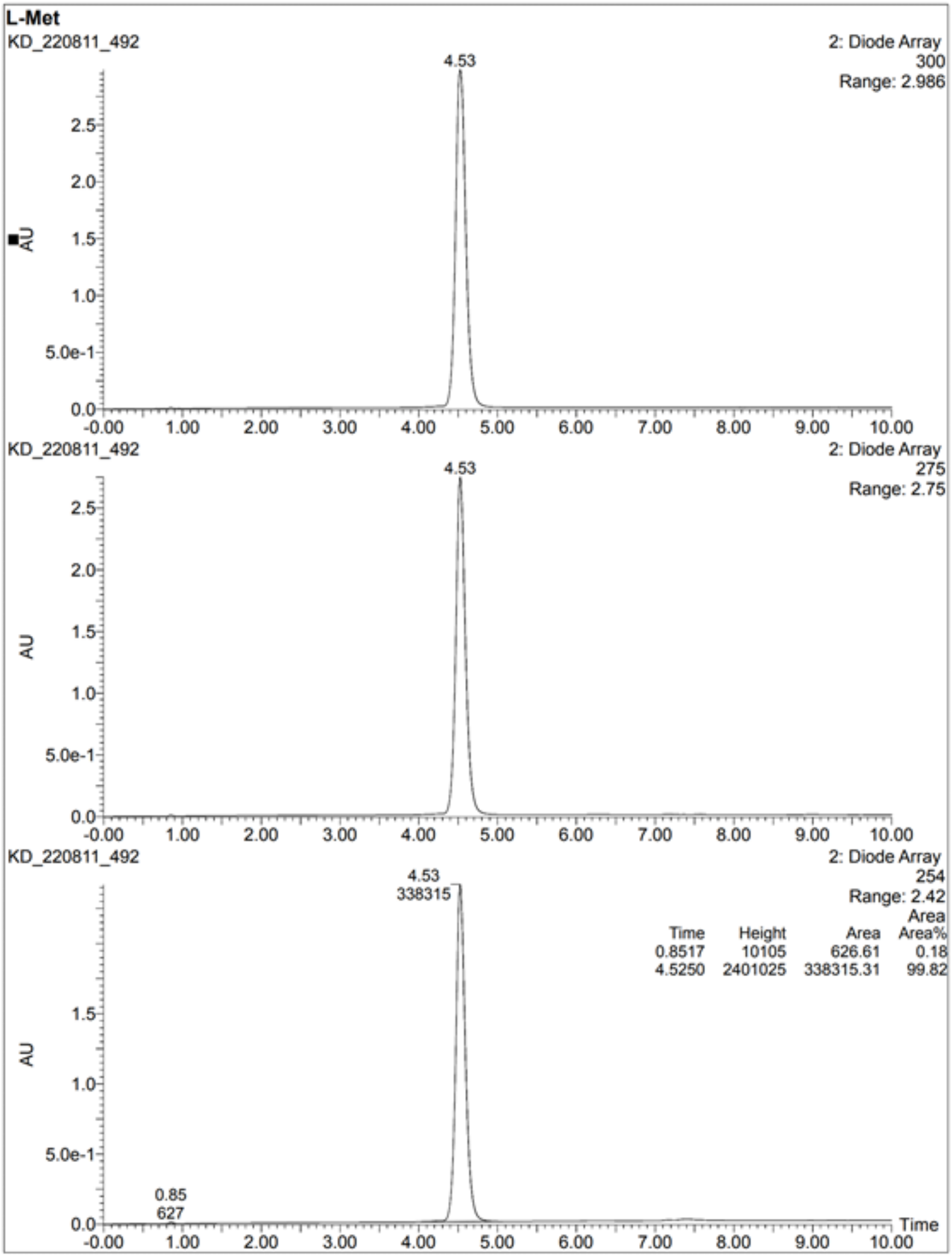
L-Met
KD_220811_492
2: Diode Array
300
Range: 2.986
4.53
AU
KD_220811_492
2: Diode Array
275
Range: 2.75
4.53
AU
KD_220811_492
2: Diode Array
254
Range: 2.42
4.53
338315
Time Height Area Area Area%
0.8517 10105 626.61 0.18
4.5250 2401025 338315.31 99.82
0.85
627
AU
Time

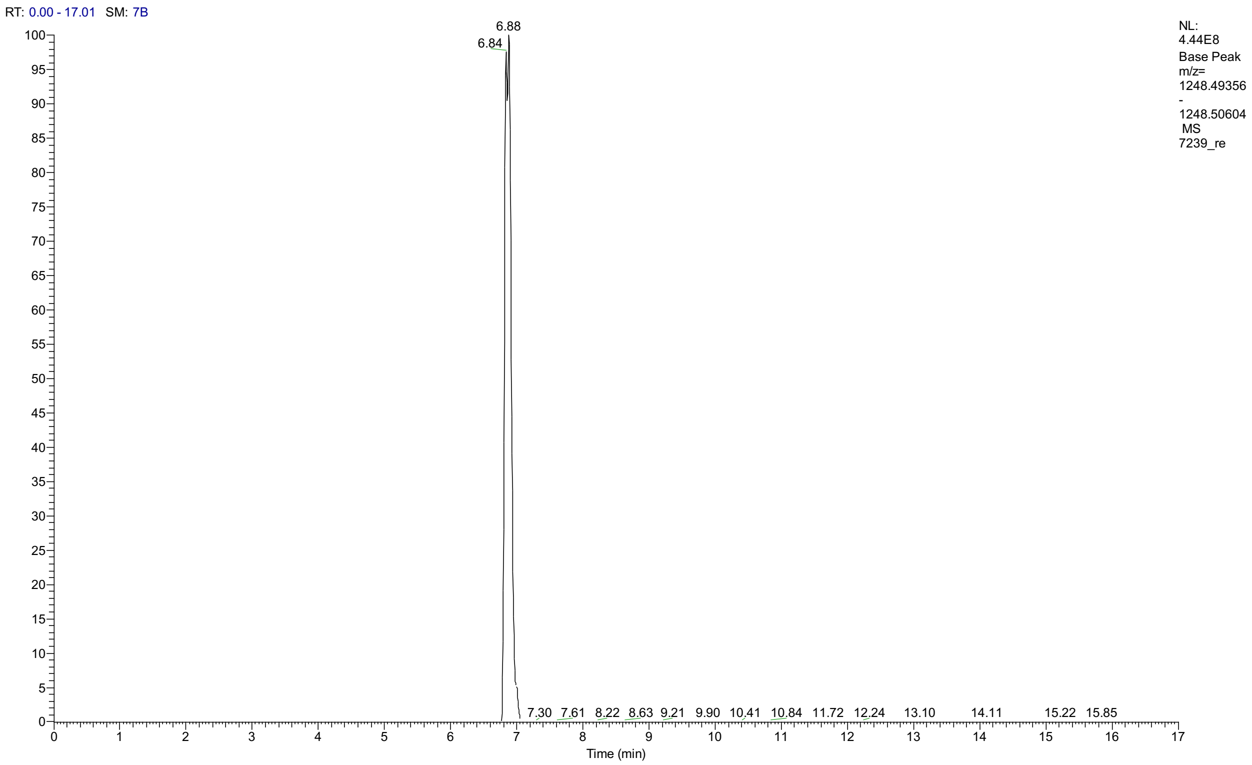
RT: 0.00 - 17.01 SM: 7B
NL: 4.44E8 Base Peak m/z= 1248.49356 - 1248.50604 MS 7239_re
6.88
6.84
7.30 7.61 8.22 8.63 9.21 9.90 10.41 10.84 11.72 12.24 13.10 14.11 15.22 15.85
Time (min)

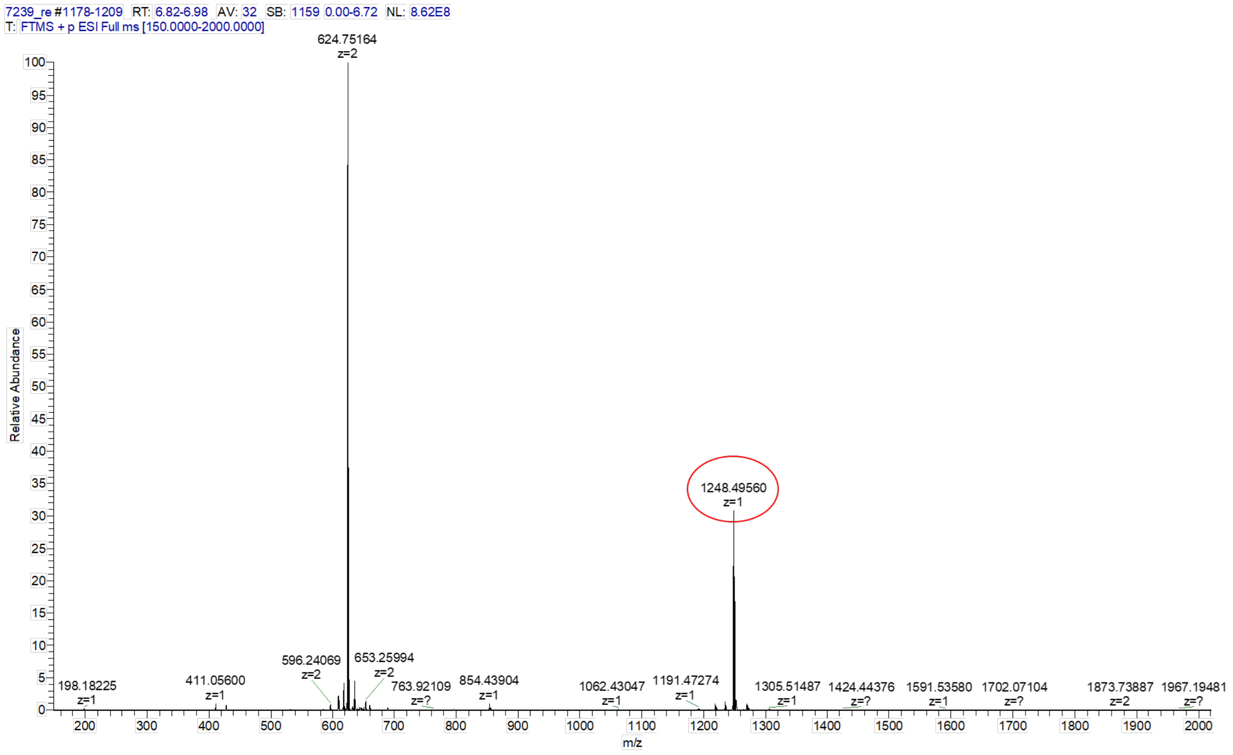
7239_re #1178-1209 RT: 6.82-6.98 AV: 32 SB: 1159 0.00-6.72 NL: 8.62E8
T: FTMS + p ESI Full ms [150.0000-2000.0000]
624.75164 z=2
1248.49560 z=1
198.18225 z=1
411.05600 z=1
596.24069 z=2
653.25994 z=2
763.92109 z=?
854.43904 z=1
1062.43047 z=1
1191.47274 z=1
1305.51487 z=1
1424.44376 z=?
1591.53580 z=1
1702.07104 z=?
1873.73887 z=2
1967.19481 z=?
Relative Abundance
m/z

(3-(2-(2-(4-(4-((7-(3-(methylsulfonamido)phenyl)thieno[3,2-d]pyrimidin-2-yl)amino)phenyl)piperazin-1-yl)ethoxy)ethoxy)propanoyl)-*L*-prolyl-*L*-prolyl-*L*-prolyl-*D*-methionyl-*L*-alanylglycylglycine **(27)**.

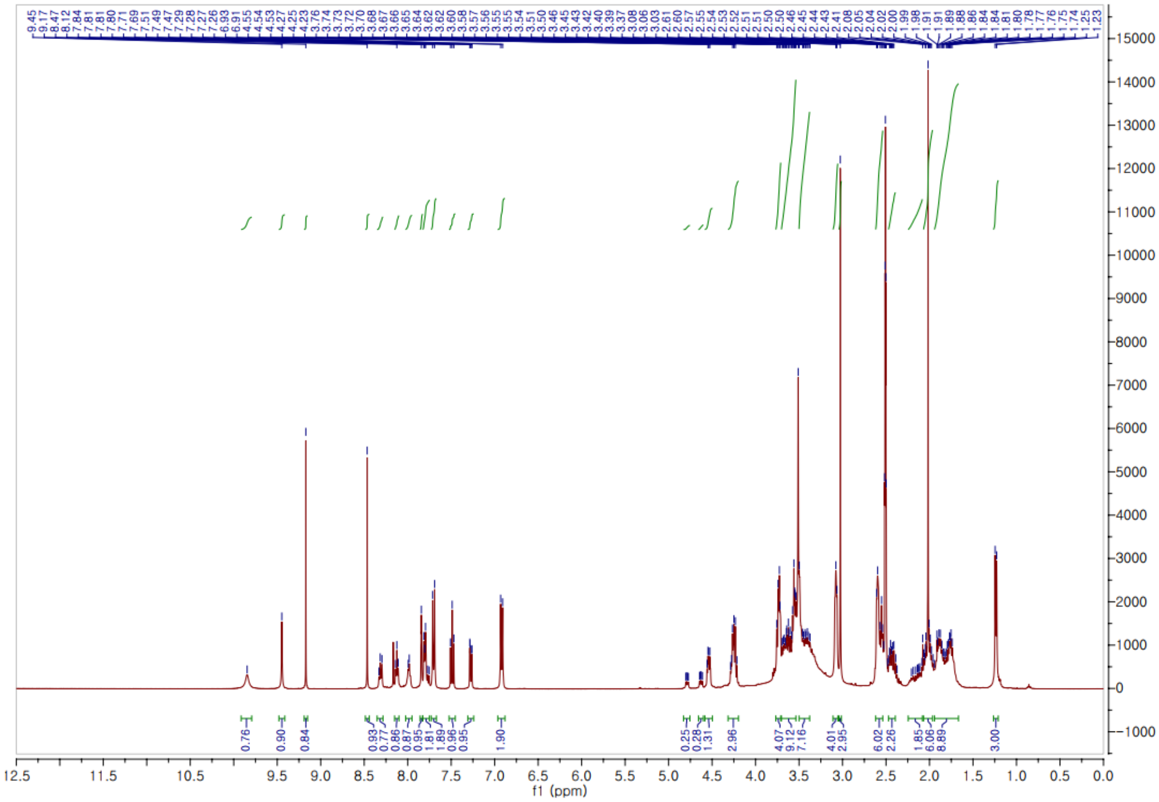

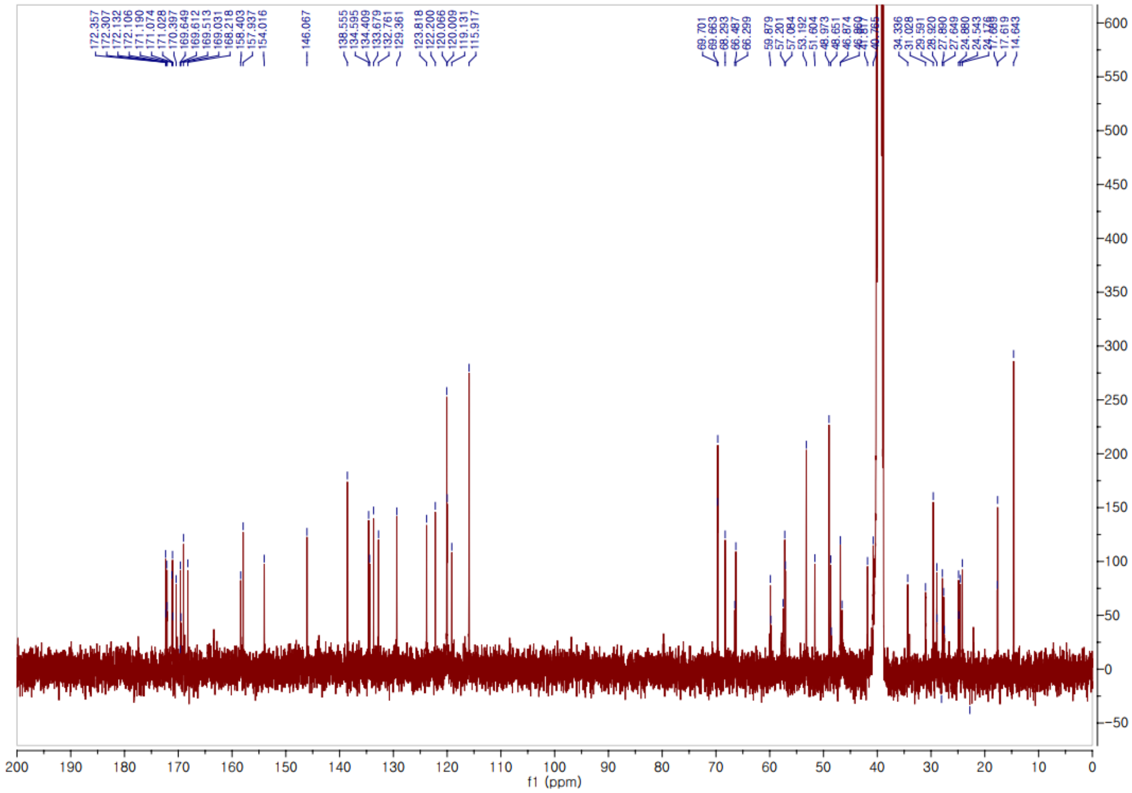

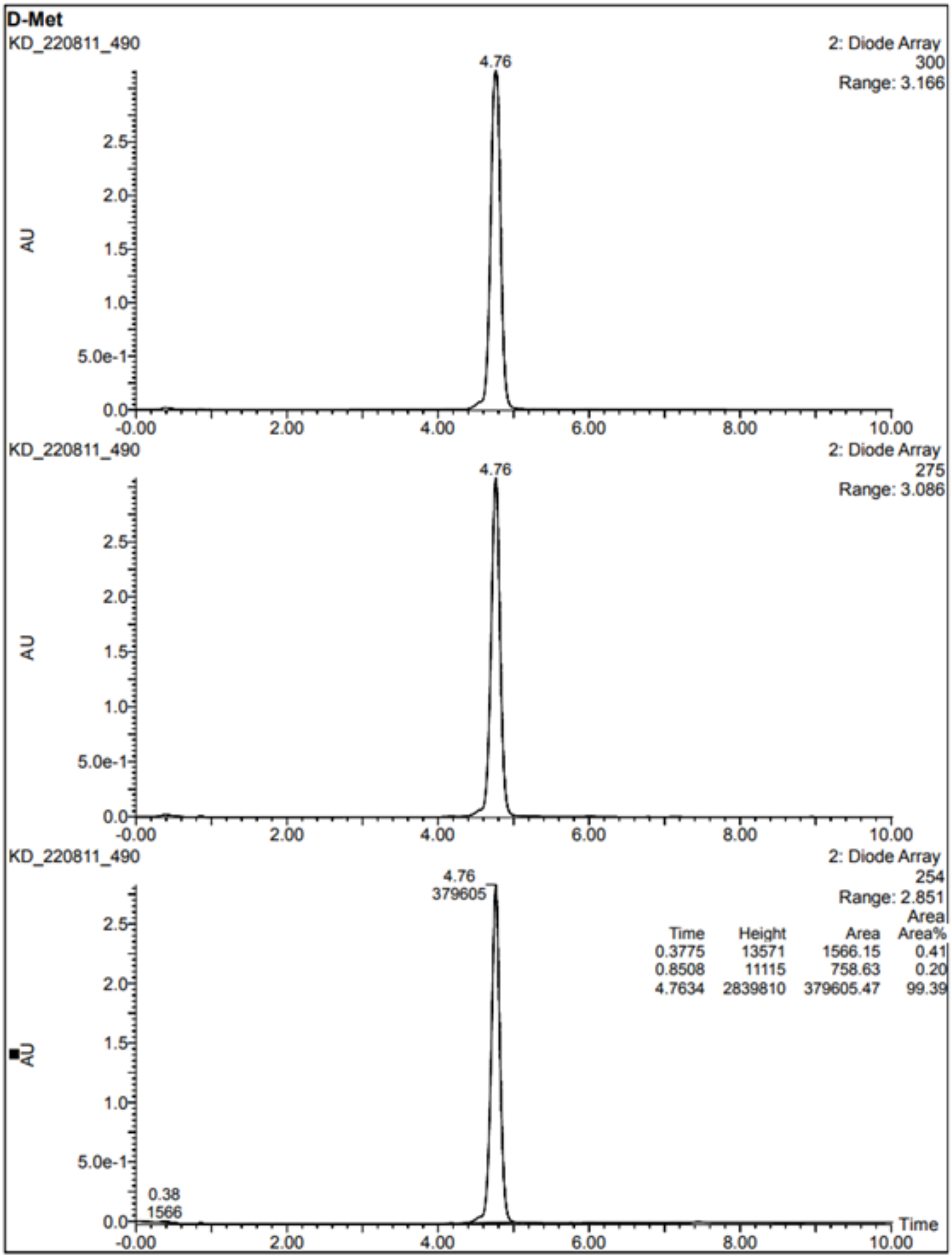
D-Met
KD_220811_490
2: Diode Array
300
Range: 3.166
4.76
AU
KD_220811_490
2: Diode Array
275
Range: 3.086
4.76
AU
KD_220811_490
2: Diode Array
254
Range: 2.851
4.76
379605
0.38
1566
Time Height Area Area%
0.3775 13571 1566.15 0.41
0.8508 11115 758.63 0.20
4.7634 2839810 379605.47 99.39
AU
Time

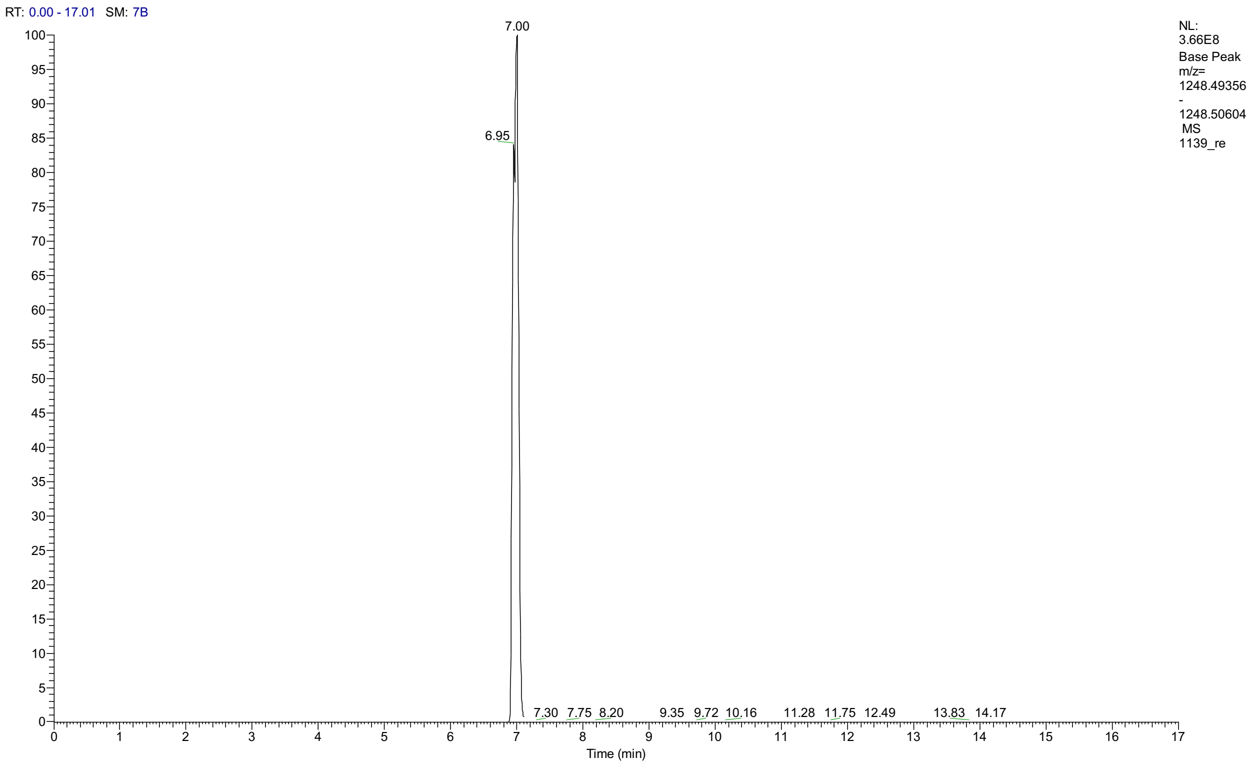

RT: 0.00 - 17.01 SM: 7B
7.00
6.95
7.30 7.75 8.20 9.35 9.72 10.16 11.28 11.75 12.49 13.83 14.17
NL: 3.66E8 Base Peak m/z= 1248.49356 - 1248.50604 MS 1139_re
Time (min)


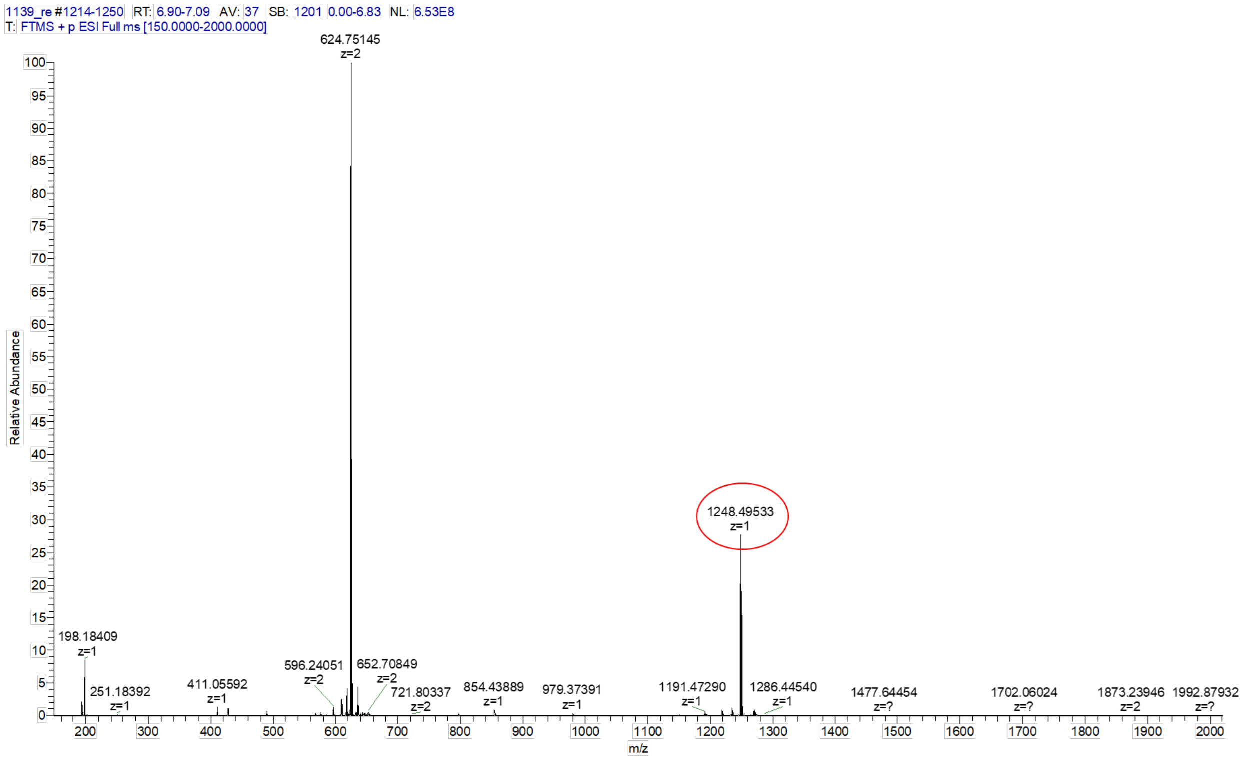

1139_re #1214-1250 RT: 6.90-7.09 AV: 37 SB: 1201 0.00-6.83 NL: 6.53E8
T: FTMS + p ESI Full ms [150.0000-2000.0000]
624.75145 z=2
1248.49533 z=1
198.18409 z=1
251.18392 z=1
411.05592 z=1
596.24051 z=2
652.70849 z=2
721.80337 z=2
854.43889 z=1
979.37391 z=1
1191.47290 z=1
1286.44540 z=1
1477.64454 z=?
1702.06024 z=?
1873.23946 z=2
1992.87932 z=?
Relative Abundance
m/z

*O*-methyl-*N*-(3-(2-(2-(4-(4-((7-(3-(methylsulfonamido)phenyl)thieno[3,2-d]pyrimidin-2-yl)amino)phenyl)piperazin-1-yl)ethoxy)ethoxy)propanoyl)-*L*-prolyl-*L*-prolyl-*L*-prolyl-*L*-homoseryl-*L*-alanylglycylglycine **(28)**.

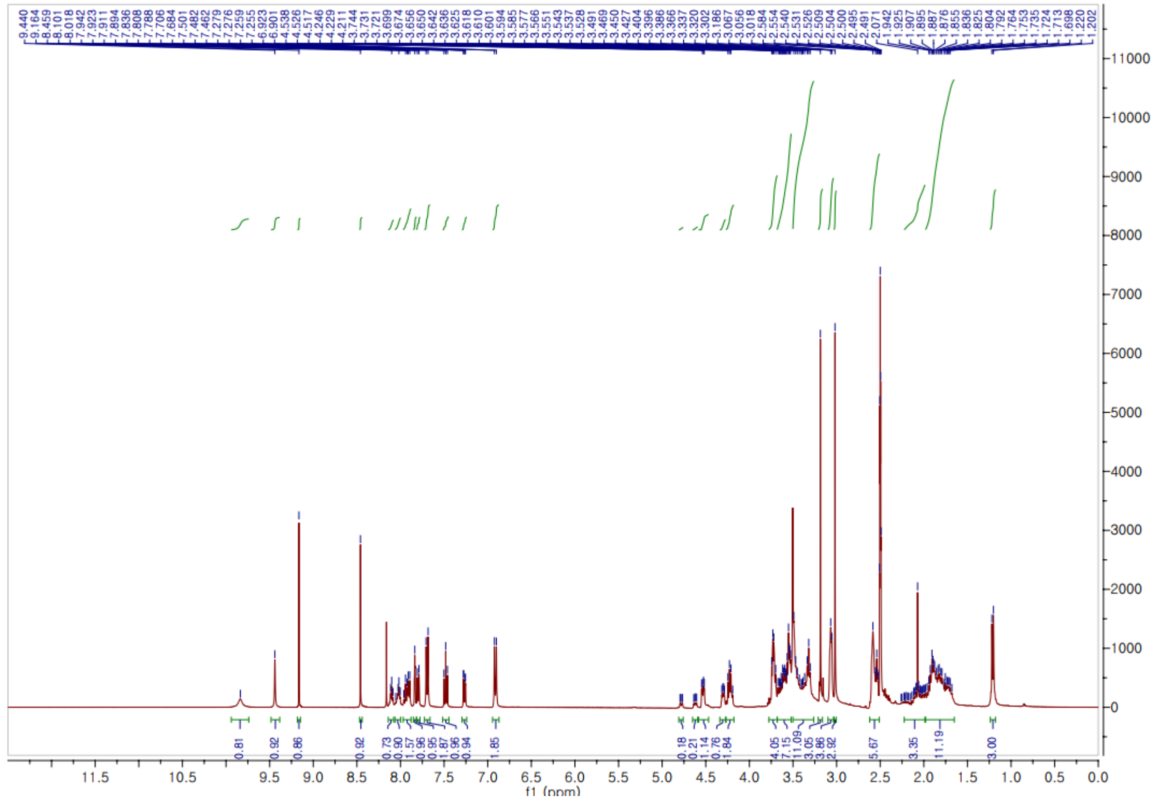


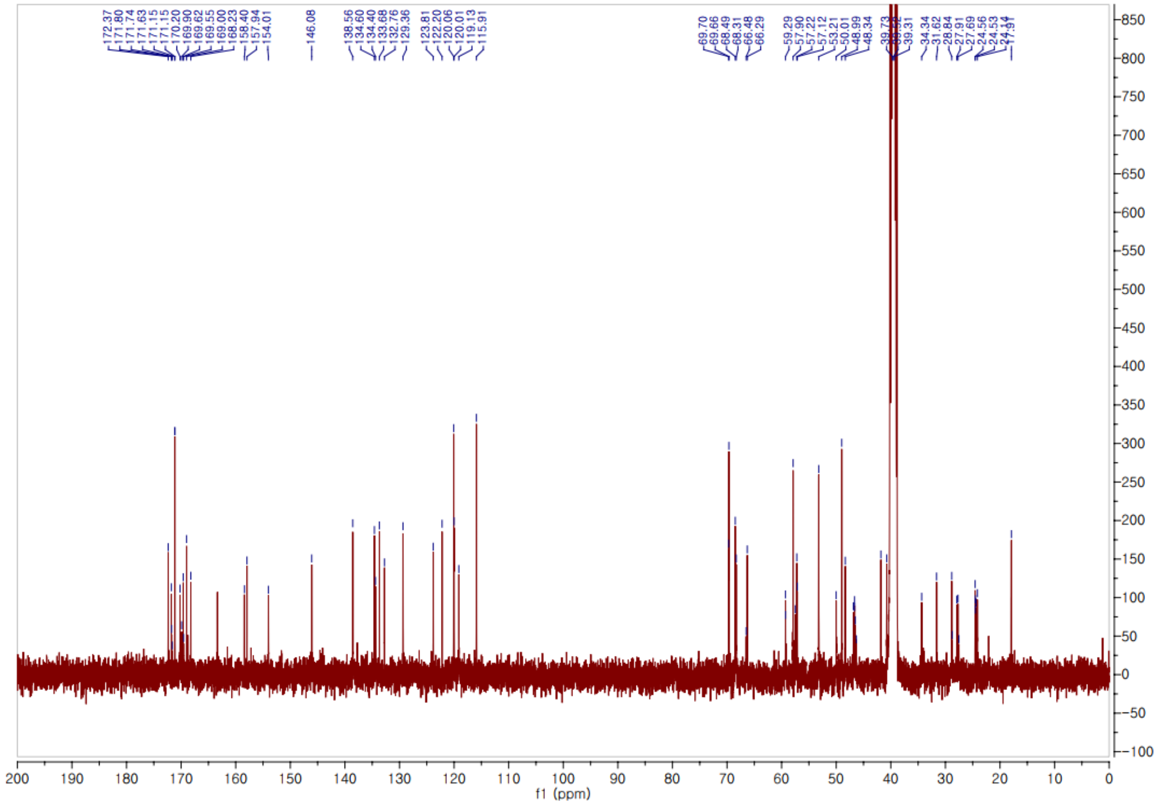

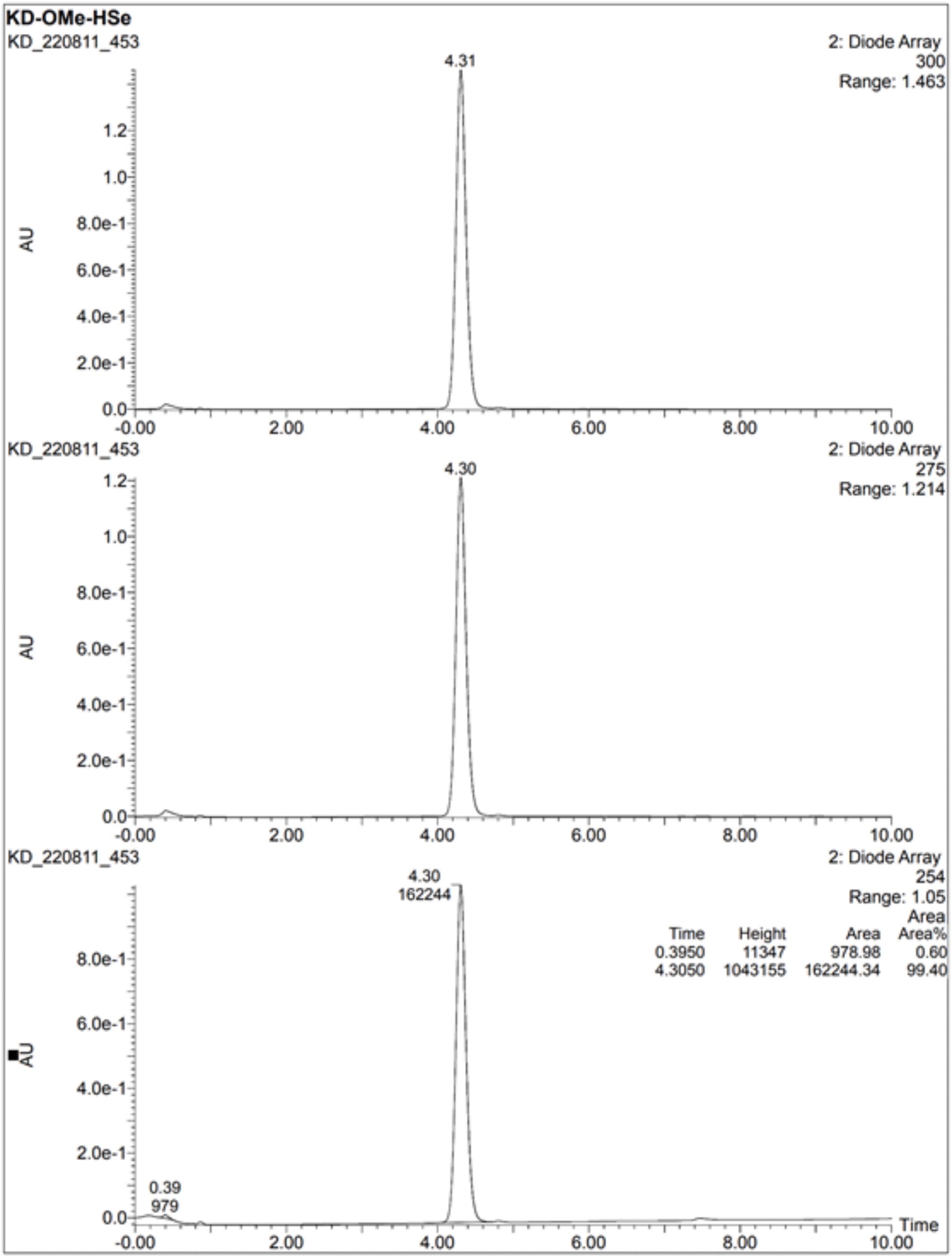
KD-OMe-HSe
KD_220811_453
2: Diode Array
300
Range: 1.463
4.31
AU
KD_220811_453
2: Diode Array
275
Range: 1.214
4.30
AU
KD_220811_453
2: Diode Array
254
Range: 1.05
4.30
162244
0.39
979
Time Height Area Area%
0.3950 11347 978.98 0.60
4.3050 1043155 162244.34 99.40
AU
Time

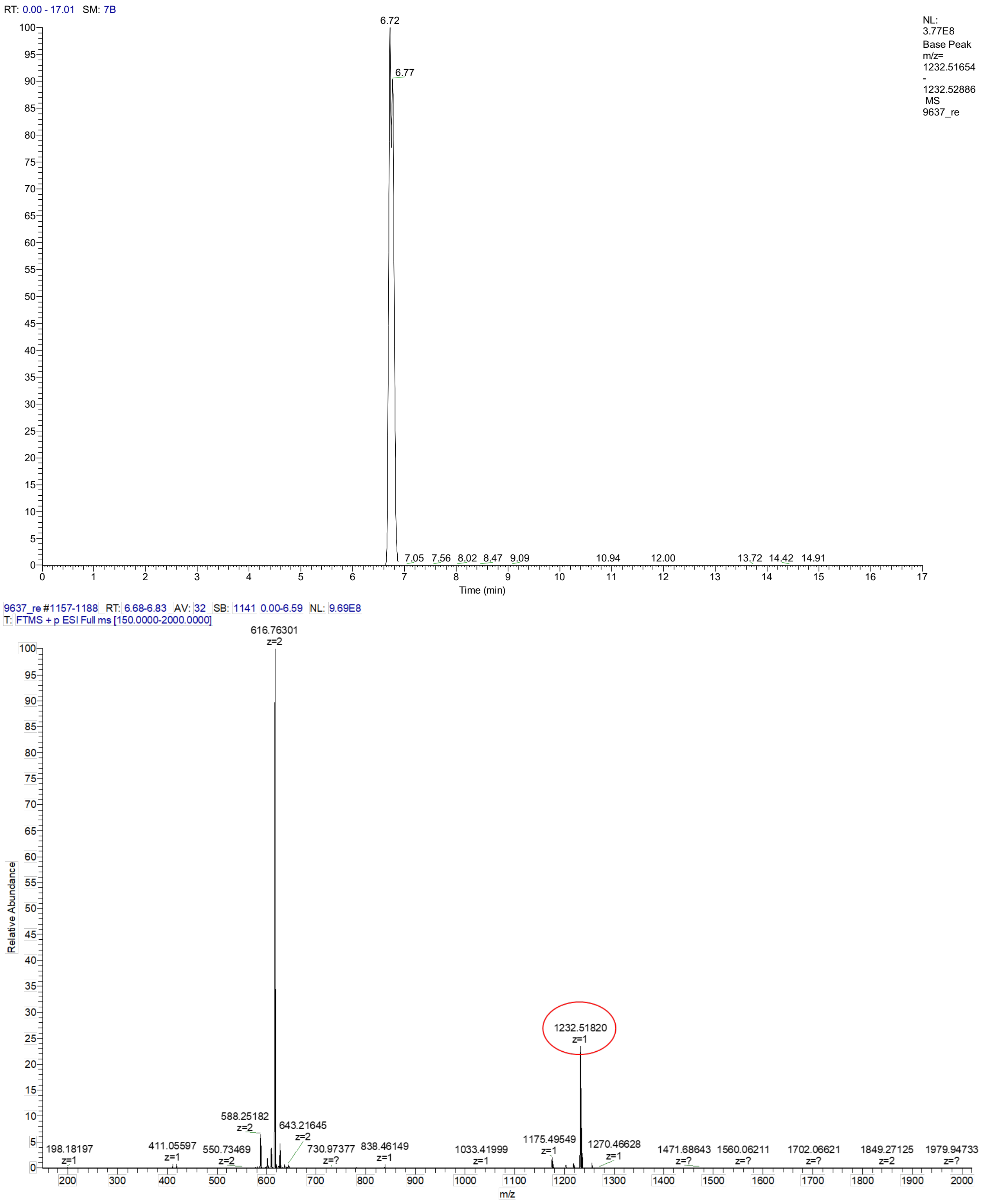
RT: 0.00 - 17.01 SM: 7B
NL: 3.77E8 Base Peak m/z= 1232.51654 - 1232.52886 MS 9637_re
6.72
6.77
7.05 7.56 8.02 8.47 9.09 10.94 12.00 13.72 14.42 14.91
Time (min)
9637_re #1157-1188 RT: 6.68-6.83 AV: 32 SB: 1141 0.00-6.59 NL: 9.69E8
T: FTMS + p ESI Full ms [150.0000-2000.0000]
Relative Abundance
616.76301 z=2
1232.51820 z=1
588.25182 z=2
643.21645 z=2
198.18197 z=1
411.05597 z=1
550.73469 z=2
730.97377 z=?
838.46149 z=1
1033.41999 z=1
1175.49549 z=1
1270.46628 z=1
1471.68643 z=?
1560.06211 z=?
1702.06621 z=?
1849.27125 z=2
1979.94733 z=?
m/z

1. H. Cho *et al.*, Identification of Thieno [3, 2-d] pyrimidine Derivatives as Dual Inhibitors of Focal Adhesion Kinase and FMS-like Tyrosine Kinase 3. *Journal of Medicinal Chemistry* **64**, 11934-11957 (2021).
2. S. Ryu *et al.*, Synthesis and structure-activity relationships of targeted protein degraders for the understudied kinase NEK9. *Current Research in Chemical Biology* **1**, (2021).
